# Shared potential metabolism trends in degraded soils and type 2 diabetes gut microbiomes

**DOI:** 10.1101/2025.03.11.642605

**Authors:** Craig Liddicoat, Bart A. Eijkelkamp, Timothy R. Cavagnaro, Jake M. Robinson, Kiri Joy Wallace, Andrew D. Barnes, Garth Harmsworth, Damien J. Keating, Robert A. Edwards, Martin F. Breed

**Author notes:** **Corresponding author:** Craig Liddicoat.

## Abstract

Global change profoundly impacts microbial systems but cascading effects on human metabolic health remain largely unexplored. Type 2 diabetes (T2D) is shaped by nutrition, host and environmental factors, with rapidly increasing global prevalence. Soil microbiomes shift with ecosystem degradation and influence metabolism through shaping food quality and gut microbiomes, including metabolite exposures without requiring colonization. Here, we explored functional overlaps between degraded soil microbiomes from five ecosystem quality gradients and gut microbiomes in T2D. We developed a method to translate metagenomic functional pathways to potential metabolism of biochemical compounds. In silico trend analyses revealed consistent shifts relevant to energy harvesting and management. Degraded soil microbiomes and T2D gut microbiomes exhibited increased potential metabolism for sugars and decreased potential metabolism for lignin and monomethyl branched-chain fatty acids. Our results advance the hypothesis that soil-ecosystem degradation may contribute to T2D pathogenesis through nutrient-depleted food and/or adverse shaping of gut microbiome functional capacities.

## Introduction

Microbiomes connect ecosystems and humans under One Health^1^. Yet, microbiome-mediated impacts of accelerating global change^2^ on human metabolic health have received little attention. Type 2 diabetes (T2D) is a major chronic multifaceted metabolic disease, characterized by insulin resistance, high blood sugar, and linked to host, nutritional and environmental factors^3, 4^. As T2D disease and economic burdens rise rapidly worldwide^5^, understanding its complex pathogenesis is critical. The gut microbiome is integral to human metabolism^6, 7^, producing a vast enzymatic repertoire that complements human digestive capabilities^8^ and metabolites that modulate appetite, gut motility, hormonal signaling and gene expression^7^. Gut microbiomes shift in T2D and with metformin treatment^9^, while fecal microbiome transplantations can improve clinical outcomes^10^. Given that long-term diet and environmental exposures have major influences on the gut microbiome^11, 12^, which is intricately linked to metabolism and T2D risk, there is a pressing need to identify upstream nutritional and environmental factors that influence this crucial interface to human metabolic health.

Beyond diet and lifestyle, living environments also shape T2D risk^3^. Microbiomes are a hypothesized link between environments and gut-associated health^13, 14^. Soils host most microbial diversity^15^, and soil exposures (for example, direct contact, foodborne and airborne dust), especially from a young age, help shape gut microbiomes^12, 13, 16^. Wild primate gut communities vary with soils^17^. Mice gut microbiomes form distinct functional composition when raised in desert, grassland or forest soils^16^. Soil microbiomes also shape food quality through nutrient acquisition^18^, incorporating microbial metabolites into plants^19^, and influencing gut microbiota of ruminants^20^. Beyond microbes per se, microbiomes also comprise metabolites^21^ and potentially transferable genetic material^22^ with structural, functional and nutritional roles^6, 19^. Microbial activity and turnover in soils and the gut may accumulate, deplete and/or transfer a vast range of metabolites with varying bioactivities^6, 23^. This means microbiomes can extend their influence via metabolites and necromass without requiring colonization. For example, soil sterilized to eliminate viable microorganisms has been found to reshape mice gut communities^24^. While the survival of soil microbes in the gut may be low or uncertain, the shaping of gut microbiomes by soil microbiome exposures is well-recognized^1, 16^. Moreover, soil microbiome composition and functional capacities may shift in generalizable ways with ecosystem degradation, for example, via urbanization and land-use change, with potential to impact human health^25, 26^. Together, these phenomena create pathways through which degraded environments may adversely shape soil microbiomes, food quality, and gut microbiomes, and potentially contribute to metabolic dysfunction in T2D.

In ongoing work to define ‘healthy microbiomes’, there is a need to advance from taxonomic to functional assessments^27^. Community-scale processes can also involve assemblages using shared resources in extracellular space^6^. Striving for deeper functional insights and interpretability, metabolite-or compound-oriented approaches^26^ aim to move beyond metabolic pathways towards measures related to specific chemical compounds (such as carbohydrates, lipids, proteins, often linked to microbial feedstocks or by-products; Supplementary Fig. 1). Compound-scale insights should offer more foundational information, exploit knowledge of biochemistry-health links, and point to mechanistic hypotheses and therapeutic targets. However, existing metabolite prediction frameworks^28, 29, 30^ rely on organism-specific and environment-specific approaches unsuited to highly biodiverse, poorly characterized and varying environments, such as soils^26^. Therefore, there is a need for new, compound-oriented methods to examine normal versus aberrant functioning in comparative analyses across soil and gut microbiomes.

In this integrative, hypothesis-generating study, we aimed to explore potential links between ecosystem degradation and metabolic alterations in the T2D gut. Specifically, we investigated shared functional potential trends in degraded soil microbiomes from five ecosystem quality gradients (‘soil datasets’): post-mining restoration^31^, disturbed versus natural soils^26, 32^, prairie restoration^33^, plantation succession^34^, and vegetation succession^35^; together with gut microbiomes in two case studies of T2D versus normal healthy controls from China^36^ and Sweden^37^ (Extended Data Table 1). We translated metagenomic functional gene-associated biochemical pathways to ‘compound processing potential’ (CPP, %), reflecting potential metabolism of individual compounds. While not measuring compounds directly, our method quantified the capacity of metagenomes to metabolize or process them. Examining (1) 30 example compounds relevant to soil and gut health (Extended Data Table 2) and (2) via compound-wide trend analyses across the soil and T2D datasets, we identified shared trends relevant to energy harvesting and management.

The CPP method provides compound-associated functional potential data by reallocating functional pathway relative abundances across all hypothetically linked chemical reactions and compounds, without directionality or thermodynamic constraints. The pipeline does not discern anabolism, catabolism, reactants or products. Strictly speaking, CPP values do not imply net production versus consumption, do not quantify in vivo/in situ compound concentrations or fluxes, and are sensitive to annotation choices and reaction-mapping structure. Notwithstanding these limitations, CPP represents a transformation in the fundamental concept of interest from ‘a given potential functional pathway (%)’ to ‘potential metabolism involving a given compound (%)’. Microbiomes adapt over time to the availability, suitability and variability of feedstock compounds and environments. Analogously, a weightlifter’s physical capacity (∼CPP) is shaped by personal development (∼community assembly), exercise routines (∼functional pathways), time, gym environment, and the weights (∼compounds) they become equipped to use, which are typically, but not always, close by. Therefore, we interpret changes in CPP values as shifts in functional capacity, linked to altered capacity for reaction sequences and hence indicative shifts in either feedstocks, intermediates, or products of microbial activity and turnover. We use the term ‘potential metabolism’ for brevity when describing CPP data, to reflect a hypothetical share of functional potential associated with particular compounds, distributed across many interconnected biochemical reactions.

## Results

### Trends with soil-ecosystem degradation

Soil metagenome functional beta diversity ordinations based on CPP values capture a substantial portion of compositional variance, largely explained by ecosystem quality, in each soil dataset (Fig. 1; PCo1 + PCo2 mean ± s.d. = 72.5 ± 6.2 % variance explained; PERMANOVA R^2^ mean ± s.d. = 0.62 ± 0.25). Approximately consistent, contrasting ATP/ADP ratios^38^ based on CPP values suggest metabolically associated divergence of metagenomes with ecosystem quality, with reduced values (ATP/ADP ∼1.65) in degraded (younger, disturbed) soils and increased values (∼1.70) in older, natural and reference soils (Fig. 1).

**Fig. 1.**
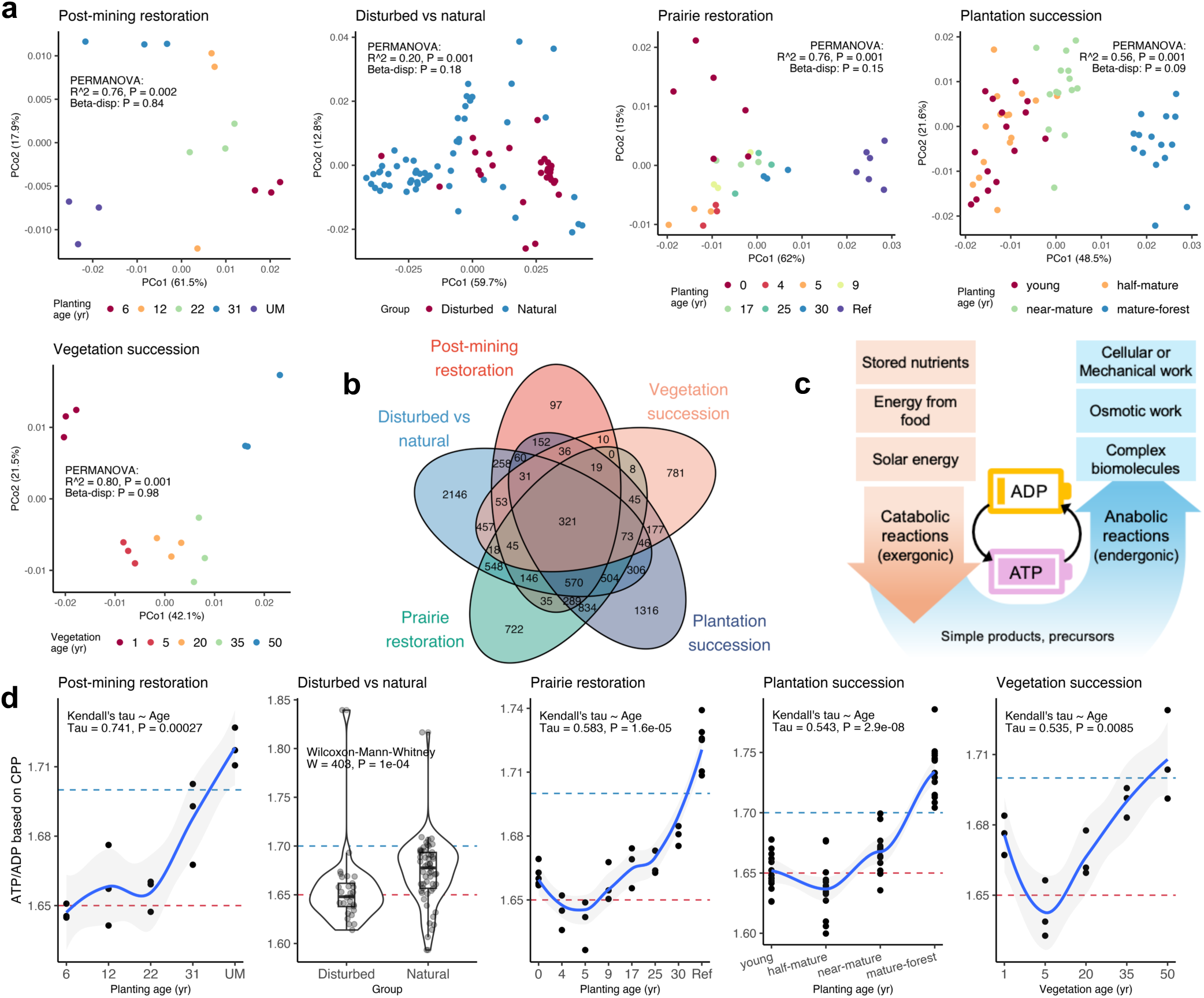
Ecosystem quality gradients. a, Beta diversity ordinations of compound processing potential (CPP) values show clustering of soil metagenomes with ecosystem quality. **b,** Venn diagram of overlapping directional CPP trends with degradation. **c,** ATP-ADP energy cycle (adapted from^39^) illustrating ATP’s universal role as energy currency driving anabolic reactions. **d,** ATP/ADP ratios based on CPP values show similar divergence between degraded/younger (∼1.65) versus older/reference (∼1.7) soils. Samples are described in Extended Data Table 1.

From the 30 example compounds, we found several directional CPP trends with soil degradation that were shared in three or more datasets (Extended Data Table 2; Supplementary Figs. 2-31). We found increased potential metabolism with degradation for (5 datasets) mannose, galactose and glucosamine; also (4 datasets) L-arabinose and chitin; and (3 datasets) D-fructose, sucrose, D-glucose, xylose and hydrogen sulfide. We found decreased potential metabolism for (5 datasets) propionate, butyrate, trimethylamine N-oxide (TMAO), and trimethylamine (TMA); also (4 datasets) lignin, glycogen, methane, carbon dioxide; and (3 datasets) amylopectin, acetate and p-cresol.

Compound-wide trend analyses identified 2,256 compounds with consistent directional (i.e., increasing vs. decreasing) CPP trends with degradation in three or more soil datasets (shared trends numbered *n* = 321 across 5 datasets, *n* = 738 across 4 datasets, *n* = 1197 across 3 datasets; Fig. 1b, trend data are in Supplementary Material). All compound-wide soil trends were *P*-adjusted for multiple testing. Carbon-containing compounds showing CPP trends with degradation were mapped using chemical formula ratios oxygen:carbon (O:C), hydrogen:carbon (H:C) and nitrogen:carbon (N:C) to visualize energetically and chemically similar compounds that trend together (post-mining restoration CPP trends are in Fig. 2; other soil datasets in Supplementary Figs 32-36, including an example showing all carbon-containing compounds within a single sample).

**Fig. 2.**
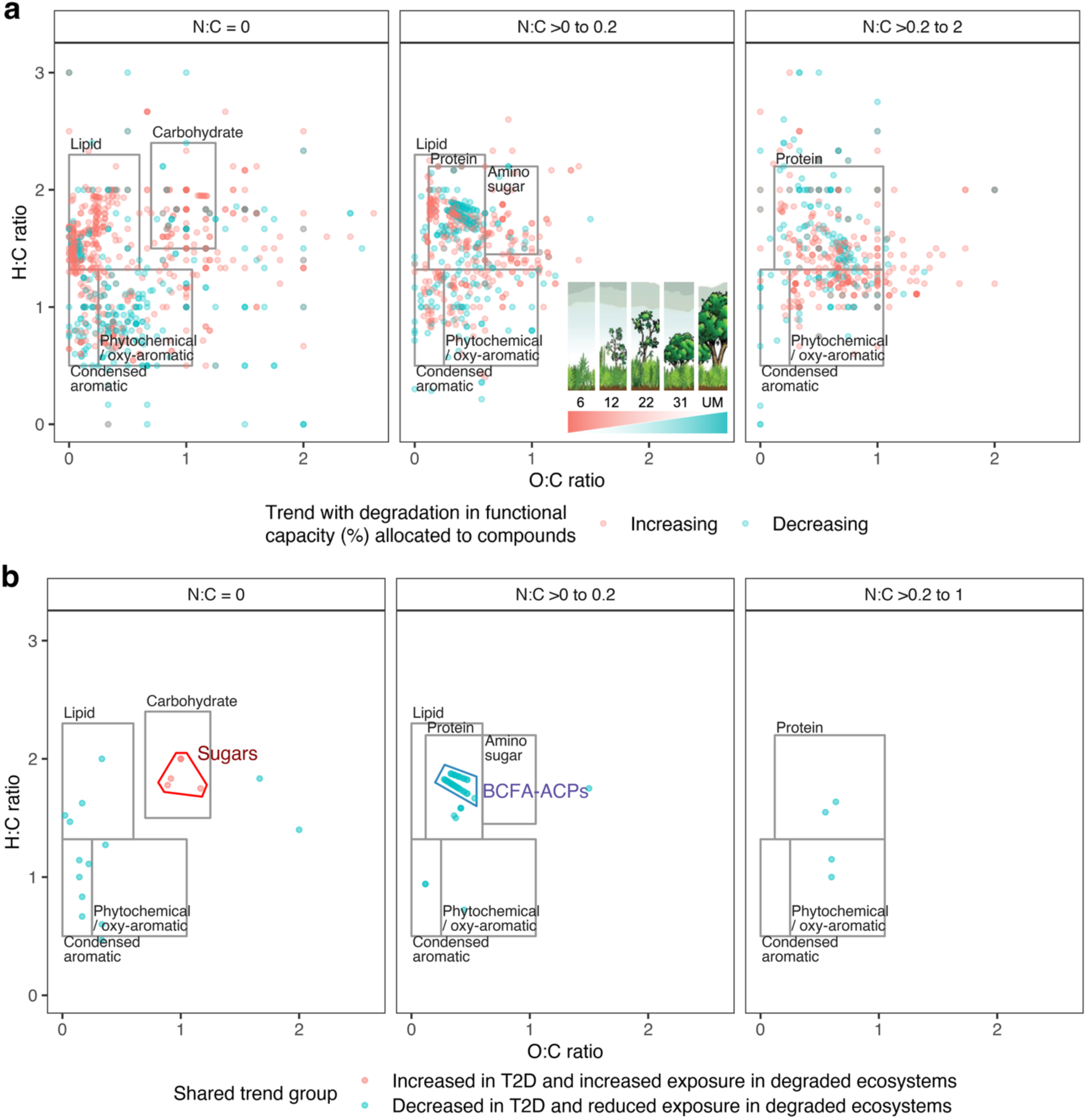
Visualizing CPP trends via chemical ratio mapping. a, Compound-wide trend analysis in post-mining restoration soils found *n* = 2,122 compounds with *P*-adjusted compound processing potential (CPP) trends (*n* = 1365 increased, *n* = 757 decreased). Points represent the carbon-containing compounds with increased (red, *n* = 1,255) or decreased (blue, *n* = 720) potential metabolism with ecosystem degradation. Each compound is mapped via its chemical formula O:C, H:C and N:C elemental ratios, where clusters of points indicate energetically and chemically similar compounds. The O:C by H:C panel extents emphasize data-rich regions. Rectangular zones denoting compound classes offer a visual guide only, adapted from^40, 41^. Samples^31^ spanned planting ages 6, 12, 22, 31 years and unmined (UM), *n* = groups of 3. **b,** Chemical ratio mapping of shared directional CPP trends with degradation and T2D (*n* = 73 compounds; 68 containing carbon) – from the combination of three or more soil datasets, T2D-CHN (*P*-adjusted) and T2D-SWE (unadjusted) results – display clustering of compounds that were grouped into ‘Sugars’ and ‘BCFA-ACPs’ (branched-chain fatty acid – acyl carrier proteins; expanded views of these two groups are in Supplementary Figs. 39-40). Lignin does not appear due to its undefined chemical formula in ModelSEED.

### Trends in type 2 diabetes

From the 30 example compounds, we found consistent trends in the Chinese (T2D-CHN) and Swedish (T2D-SWE) cohort datasets (Extended Data Table 2; Supplementary Figs. 2-31) for increased potential metabolism in T2D for D-fructose (CHN: *P* < 0.01, SWE: *P* ≤ 0.05), L-arabinose (CHN: *P* < 0.001, SWE: *P* ≤ 0.05), and amylopectin (CHN and SWE: *P* ≤ 0.05); and decreased potential metabolism in T2D for lignin (CHN and SWE: *P* ≤ 0.05). Numerous trends were only found in one T2D cohort or the other. Increased potential metabolism was found in T2D in the Chinese cohort for mannose (*P* < 0.001); galactose, xylose and glucose (all *P* < 0.01), and in the Swedish cohort for sucrose, starch and amylose (*P* ≤ 0.05). Decreased potential metabolism was found in T2D in the Chinese cohort for acetate, propionate, butyrate, carbon dioxide, hydrogen sulfide, indole (all *P* < 0.001); menaquinone (Vitamin K2), glucosamine (*P* < 0.01); methane, p-cresol and serotonin (all *P* ≤ 0.05). We found conflicting potential metabolism trends for ammonia in T2D, which decreased in T2D-CHN (*P* < 0.001), but increased in T2D-SWE (*P* < 0.01).

In the compound-wide analysis of T2D-CHN data (with *P*-adjustment) we found a total of 3,741 compounds showed trending potential metabolism with T2D (*n* = 255 increased, *n* = 3,486 decreased). Similar correction for multiple testing across >7,000 compounds did not yield *P*-adjusted trends in the Swedish (T2D-SWE) dataset. However, to explore possible overlaps within gut ecosystems (recognized as highly dynamic^11, 27^ and burdened by high dimensionality for multiple testing^42^), including indications that might support *P*-adjusted T2D-CHN findings, we considered directional CPP patterns suggested by unadjusted *P*-values in the T2D-SWE dataset. Later analyses of defined groups of compounds also relied on selecting the most interesting and/or potentially significant compounds in each dataset. Proceeding on this basis, the T2D-SWE data had 1,269 compounds with (unadjusted) trending potential metabolism (*n* = 358 increased, n = 911 decreased). From visual inspection, the chemical ratio mapping of carbon-containing compounds (Supplementary Figs. 37-38) showed limited commonality in CPP trends in the vicinity of carbohydrates, with widespread reduced CPP in the T2D-CHN dataset (compounds and trend data are in Supplementary Material).

### Consistent trends with ecosystem degradation and T2D

For the 30 example compounds, we found consistent CPP trends with degradation and both T2D datasets for increased potential metabolism in D-fructose (three soil datasets) and L-arabinose (four soil datasets), and decreased potential metabolism in lignin (four soil datasets; Extended Data Table 2).

From the exhaustive compound-wide trend analyses in all datasets, we then looked for shared trends. From soils, we considered the 2,256 directional CPP trends previously identified in three or more datasets. Then, looking for simultaneous matches in compound identifiers and directional CPP trends from T2D-CHN (*P*-adjusted) and T2D-SWE (unadjusted) results, we found 73 compounds with shared CPP trends in degraded soils and T2D (*n* = 5 increased, *n* = 68 decreased; listed in Extended Data Table 3). From visualizing the subset of compounds that contained carbon in chemical mapping space (Fig. 2b; *n* = 5 increased, *n* = 63 decreased) we observed two prominent groupings: ‘Sugars’ (L-arabinose, D-fructose, melibiose, melitose and 6-phosphosucrose) showed increased CPP in degradation and T2D; and ‘BCFA-ACPs’ comprising 36 compounds dominated by branched-chain fatty acid – acyl carrier proteins (Extended Data Table 4) showed decreased CPP in degradation and T2D. The BCFA-ACPs group were all involved in fatty acid biosynthesis and dominated by monomethyl BCFAs that were not found elsewhere in the datasets beyond this trend group. Other compounds with decreased CPP in degradation and T2D included: lignin and precursors sinapyl alcohol and p-coumaryl alcohol; ubiquinone precursors^43^ (2-Octaprenylphenol and 3-Octaprenyl-4-hydroxybenzoate); and molybdate. Summary mean patterns across soils and T2D for shared CPP trends are shown in Fig. 3, and trends across all datasets in summed CPP values for highlighted groups of BCFA-ACPs, sugars, and lignin and precursors are shown in Fig. 4. Thirty-five of 36 BCFA-ACPs, three of the five sugars, lignin and precursors, and molybdate each had decreased CPP in four out of five soil datasets. The ubiquinone precursors had decreased CPP in all five soil datasets.

**Fig. 3.**
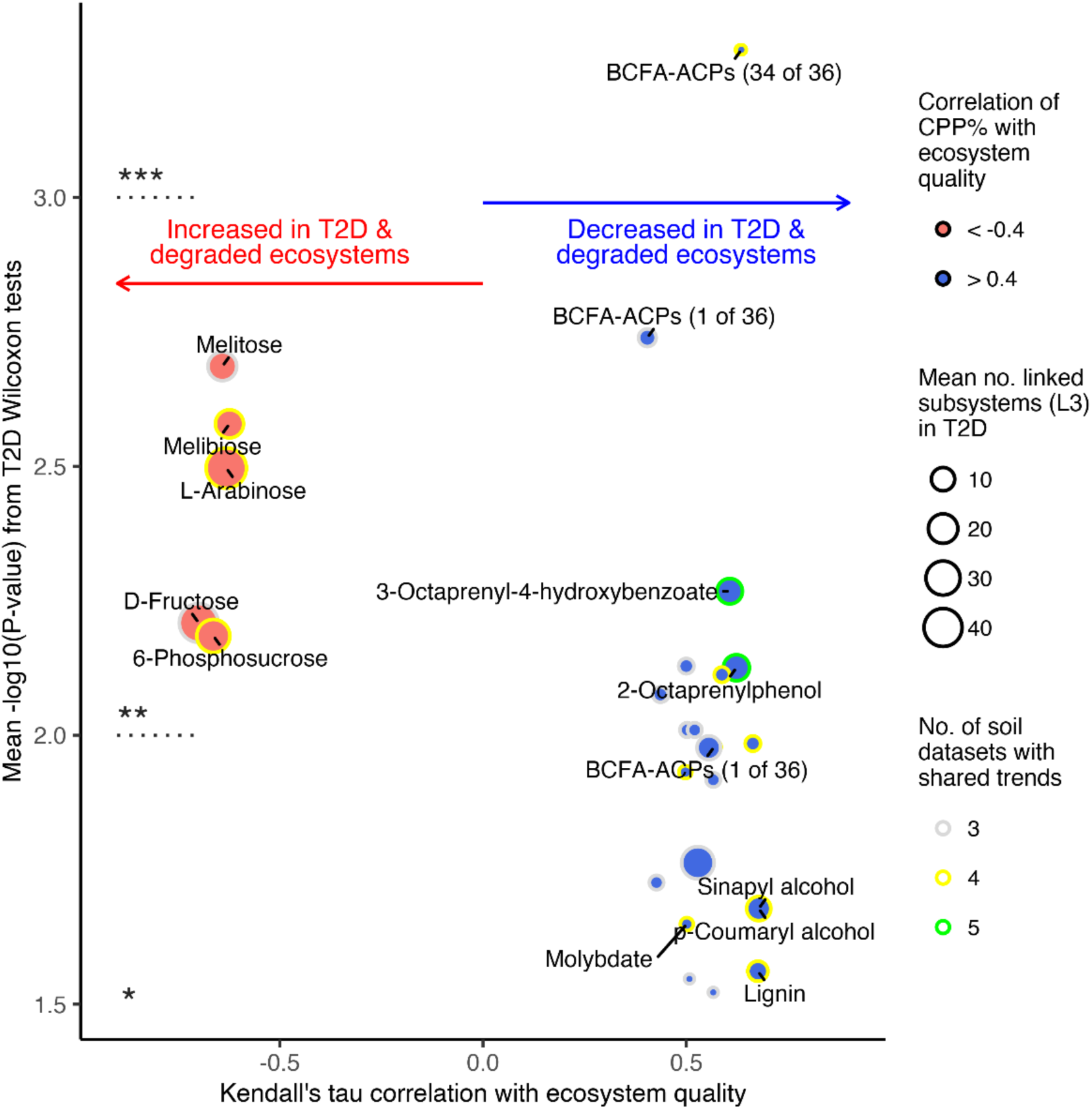
Shared potential metabolism trends in degradation and T2D. Points indicate compounds with increased (red, *n* = 5) or decreased (blue, *n* = 68) compound processing potential (CPP) based on shared trends in 3 or more soil datasets, *P*-adjusted trends in Chinese (CHN), and unadjusted trends in Swedish (SWE) type 2 diabetes (T2D) cohort datasets. The plot combines mean correlations with soil ecosystem quality (x-axis), mean-log10(*P*-values) from Wilcoxon difference testing of T2D versus normal subjects (y-axis), mean numbers of linked functional themes (level 3 subsystems) from the T2D datasets, and the number of soil datasets exhibiting the shared trends. BCFA-ACPs are compounds (*n* = 36) dominated by branched-chain fatty acid-acyl carrier proteins and involved in fatty acid biosynthesis. Compounds 2-Octaprenylphenol and 3-Octaprenyl-4-hydroxybenzoate are precursors in ubiquinone biosynthesis. Indicative *P*-value thresholds are: *** = –log10(*P*) > 3, ** = 3 2≥ –log10(*P*) > 2, * = 2 2≥ –log10(*P*) 2≥ 1.301. All compounds are listed in Extended Data Table 3.

**Fig. 4.**
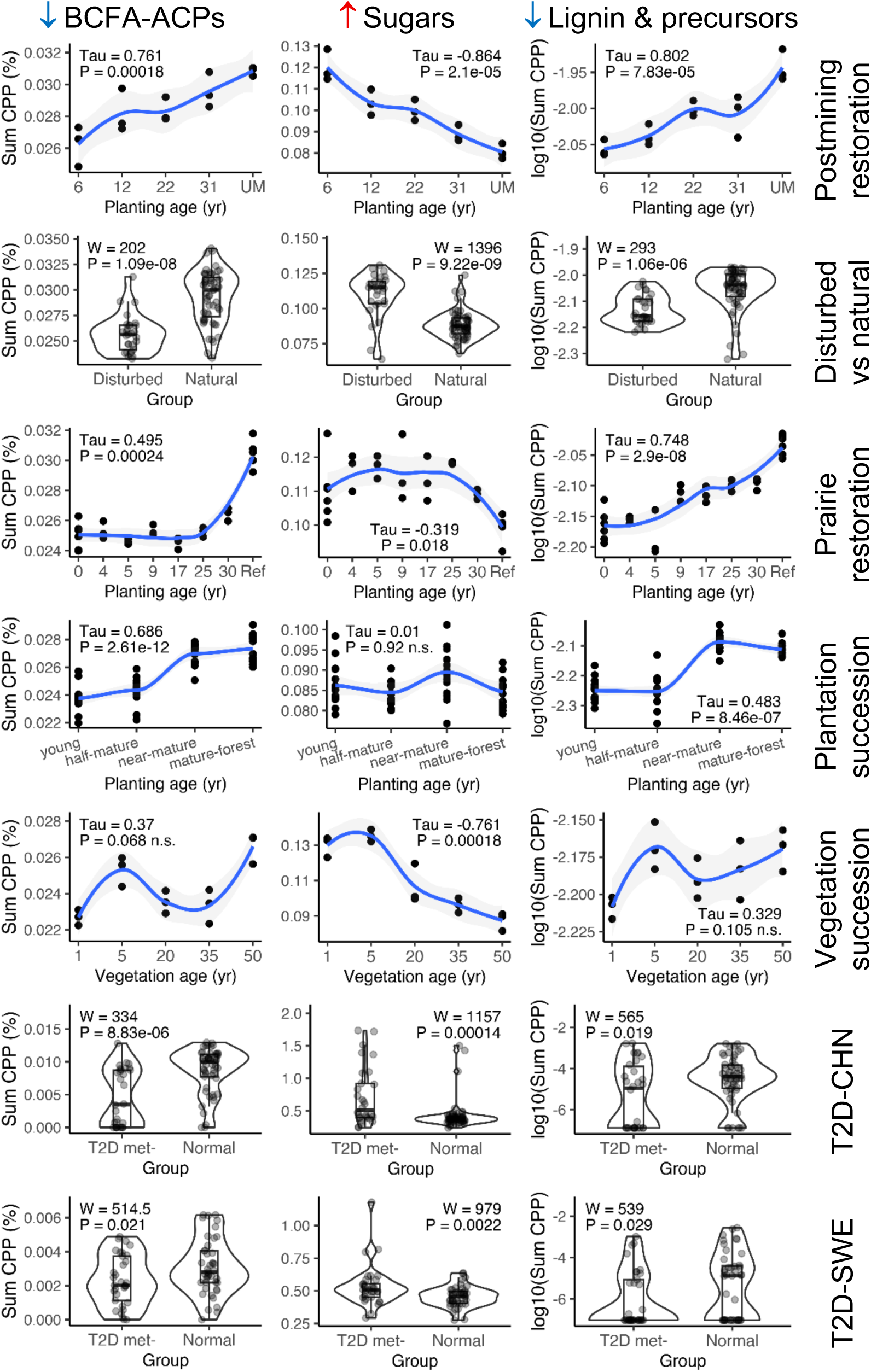
Potential metabolism trends in compound groups. The plots show trends in summed compound processing potential (CPP, %) in the groups BCFA-ACPs (*n* = 36 compounds, Extended Data Table 4), sugars (comprising D-fructose, L-arabinose, melibiose, 6-phosphosucrose, and melitose), and lignin plus precursors (sinapyl alcohol and p-coumaryl alcohol), identified from individual compound trends occurring in 3 or more of 5 soil datasets, *P*-adjusted trends in Chinese (CHN) and unadjusted trends in Swedish (SWE) type 2 diabetes (T2D) cohorts. Testing is based on Kendall tau correlation with increasing age or ecosystem quality, and differences (Wilcoxon) between T2D versus normal. Lignin CPP values were log_10_-transformed to improve display. Sample sizes are in Extended Data Table 1.

## Discussion

We identified biologically relevant shared trends in the potential metabolism of compounds in soil and human gut microbiomes, aligning with the hypothesis that nutrient-depleted food (i.e., low BCFA and lignin content, high sugar content) and/or excessive microbiome exposures associated with degraded soil-ecosystems may contribute to development of T2D. Similar, yet separate, ecological stresses and dysfunction could shape convergent functional traits in soil and gut ecosystems. However, potential linkages across these ecosystems also warrant attention due to the rising burden of T2D and increasing global change soil impacts – because soil microbiomes underpin food quality and shape gut microbiomes. Our compound-oriented approach facilitates access to a wealth of evidence connecting chemical compounds with metabolism, health, and broader biological processes in humans and the environment. Also, mapping compounds into the bioenergetically informed O:C, H:C, and N:C coordinate space revealed patterns of simultaneously trending compounds (i.e., sugars, BCFA-ACPs) with soil degradation and T2D. Our results align with the One Health framework, suggesting functional links between ecosystem quality and human metabolic health. This contributes to research aimed at improving theoretical models of nature–human health relationships, and identifying synergies and co-benefits in advancing the UN Sustainable Development Goals.

### Do branched-chain fatty acids connect healthy ecosystems and healthy people?

Our results suggest that microbiomes in degraded ecosystems and T2D have reduced capacity for monomethyl BCFA (mmBCFA) biosynthesis. This may reflect a shift in the biochemical composition of bacterial assemblages, as mmBCFAs are key components that enhance bacterial membrane fluidity, and mmBCFA-synthesis capacity enables adaptive responses for survival and proliferation under environmental stresses^44, 45^. BCFA levels vary among bacterial taxa, commonly being higher in Gram-positive bacteria, present in some Gram-negative species, and especially high in *Bacillus* and *Bifidobacterium*^44, 46^. Within ecosystems, BCFAs are present in lipid/waxy plant compounds, animal (including human) tissues and fluids, ruminant products (milk, cheese, meat), fish, organs, blood serum, skin, hair, adipose tissue, colostrum, breast milk, *vernix caseosa* of newborns, and at high levels in saprotrophic fungi such as *Conidiobolus* spp. found in soil and decaying organic matter^45, 47, 48^. Therefore, variation in BCFA synthesis capacity could result from changes in microbial taxonomic composition as ecosystems degrade.

Monomethyl BCFAs possess multiple beneficial bioactivities for human health and link bacterial and human physiology^47^. Beneficial bioactivities of mmBCFA include lipid-lowering, reducing risk of metabolic disorders, maintaining insulin-producing β-cell function and insulin sensitivity, anti-inflammatory effects, cytotoxicity against cancer cells, and regulation of development^45, 46, 47^. Bacterially-derived mmBCFAs facilitate direct biochemical uptake and cross-domain metabolism between the gut microbiome and animal hosts^49, 50^. Nematode and mammalian tissue culture studies show that mmBCFA are critical for triggering activation of the host mechanistic target of rapamycin complex 1 (mTORC1)^50^, a central regulator of cell growth and metabolism^51^. The host mTORC1 responds to environmental signals (for example, amino acid availability, growth factors, energy levels, stress) to coordinate cellular status. However, mTORC1 dysregulation is closely linked to diabetes, cancer, and neurodegenerative disorders^51^. Ref.^52^ reported high capacities for human fetal intestinal epithelial cells (enterocytes) to incorporate long-chain BCFA into membrane phospholipids. Following uptake by enterocytes, further interactions between BCFA and host endogenous BCFA synthesis and metabolism may occur^45, 47^, including post-translational modifications of cellular proteins that alter intestinal cell function^53^. While precise mechanisms are uncertain, it is possible that mmBCFA in the gut microbiome may modulate host cellular metabolism and physiological functioning via these interactions.

Microbial sources play a dominant role in supplying health-associated mmBCFA^46, 47^. Key inputs to BCFA synthesis include branched-chain amino acids, which are only made in bacteria, plants, and fungi, but not animals^54^; or shorter BCFA, which can be elongated. Microbes (for example, rumen bacteria) also contribute to BCFA synthesis in animals^47^. Therefore, both diet and gut microbiome synthesis are major sources of BCFA for humans^46, 47^. Within soil and gut microbiomes, BCFA exchange between substrates and microbes is expected because fatty acids represent high-value energy, and bacteria assimilate environmental BCFA into their bodies^55, 56^. In the gut, the constant release of mmBCFAs from cell membranes occurs during bacterial turnover from competitive interactions^57^ and the secretion of fatty acids and their derivatives^49^. A portion of these gut bacteria may be supplied from external ecosystem exposures^14^, while BCFA-synthesizing bacteria may be further shaped by diet and other factors (for example, high BCFA-content *Bifidobacterium* spp. proliferate in breastfed infants^58^).

As ecosystems degrade, there may be several reasons why BCFA-synthesizing organisms decline. Ecosystem degradation is widely associated with reductions in soil organic matter^2, 59^. Soil organic matter comprises a continuum of progressively decomposed organic compounds, commonly including amino acids and fatty acids^60^. Amino acids are released from proteins in the soil at rates correlated with soil organic matter pools and soil protein concentrations^61^, and increasing with ecosystem succession^62^. Natural decomposition and recycling networks of saprotrophic fungi (including high-BCFA-content spp.) may also be degraded, likely reducing BCFA content and turnover within soil microbiomes. Diverse plant litter typically contributes to increased soil carbon, higher microbial abundance and diversity, and greater decomposition and turnover, strengthening over time^63^. Together, these influences may reduce access to and recycling of both branched-chain amino acids and fatty acids for soil microbiomes in degraded ecosystems, thereby potentially reducing ambient human exposure to health-promoting microbial mmBCFA synthesis capacity (Fig. 5). Degraded agricultural soils might also produce crops and ruminant products (for example, meat, dairy) with reduced mmBCFA content, which impact gut microbiomes via diet.

**Fig. 5.**
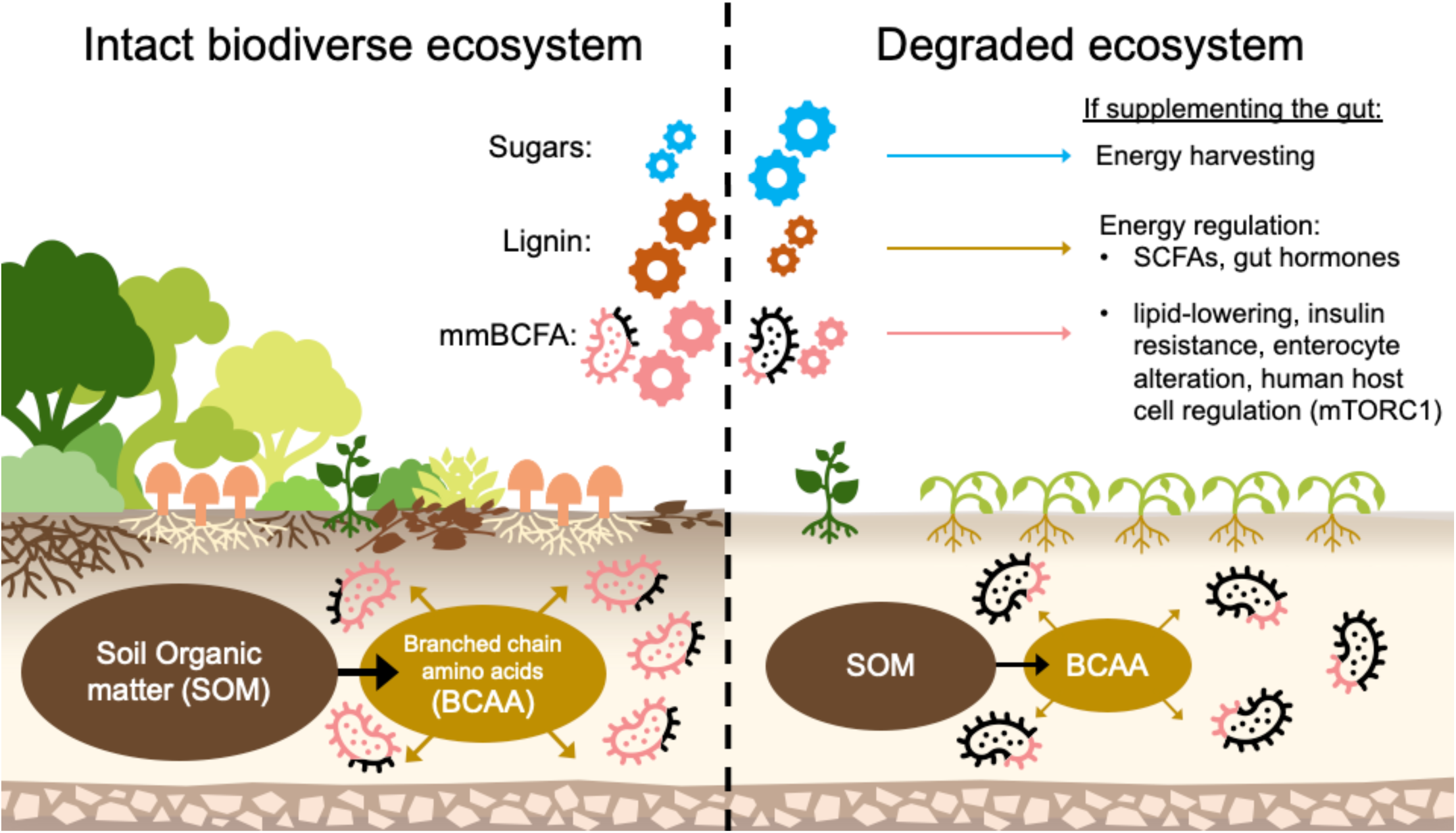
Hypothesis linking ecosystem degradation and type 2 diabetes. Intact ecosystems typically have greater biodiversity, woody plant debris (including lignin), fungal recycling and soil organic matter (SOM) pools. SOM releases branched-chain amino acid (BCAA) precursors to precious monomethyl branched-chain fatty acids (mmBCFA) used in bacterial cell membranes, stored by fungi, and needed to regulate animal cell metabolism via the mechanistic target of rapamycin complex 1 (mTORC1) among other pathways. With ecosystem degradation, soil organic matter decomposes into simple sugars. Degraded soil microbiomes have increased capacity to metabolize simple sugars, decreased capacity to metabolize lignin (potentially impacting production of short-chain fatty acids, SCFAs), and decreased mmBCFA synthesis capacity (potentially linked to mmBCFA content in microbial bodies, necromass and the wider ecosystem). If people consume nutrient-depleted food (low mmBCFA and lignin content, high sugar content) and/or are increasingly exposed to soil microbiomes associated with degraded ecosystems, it may contribute to metabolic shifts in the gut microbiome consistent with type 2 diabetes.

### Compounds linked to energy harvesting

We found the potential metabolism of sugars, L-arabinose, D-fructose, melibiose, melitose, and 6-phosphosucrose, increased with ecosystem degradation and T2D. Additionally, the potential metabolism of mannose, galactose, sucrose, glucose and xylose increased in more degraded ecosystems. Sugars are the most abundant organic compounds in the biosphere, representing key carbon and energy sources for soil microbes^64^. Sugars in soil can be derived from decomposition of plant litter, rhizodeposits, root exudates, microbes and their residues^64^. Previous work reports mannose, galactose, arabinose, glucose, and xylose comprise major sugar residues in soil organic matter^65^. Our detection of increased potential for sugar metabolism (largely monosaccharides, disaccharides, and melitose – a plant-based trisaccharide comprising galactose, glucose, and fructose) likely reflects enhanced microbial decomposition of soil organic matter, including fresh plant inputs, in more degraded ecosystems^59^.

Gut microbiome carbohydrate metabolism produces up to 10% of host energy extraction and contributes to insulin resistance, obesity and prediabetes^66^. Ref.^66^ found increased gut microbiome metabolism of monosaccharides, including fructose, galactose, mannose and xylose, correlated with insulin resistance. Interestingly, in soils and humans, enhanced sugar metabolism is associated with increased abundance of opportunistic bacteria^67^. Increased proportions of opportunists have been associated with disturbance in urban greenspace soils^25^ and human-altered land uses^68^. If microbiome components that promote enhanced sugar metabolism are transferred from degraded ecosystems to the human gut, they may foster opportunistic or pathogen-like traits, potentially contributing to increased energy harvest and metabolic imbalances associated with T2D.

### Compounds linked to energy management

We found altered potential metabolism of compounds with plausible links to energy storage via fatty acids, gut hormone secretion, and appetite regulation, which decreased in both degraded ecosystems and T2D. This raises the question whether nutrient-depleted foods and/or excessive exposures to degraded soils with deficient CPP related to energy management may shape gut microbiomes that are also deficient in this way. We discussed above the potential roles of gut microbiome mmBCFA content and biosynthesis capacity in metabolism. Dietary fiber, such as lignin, is known to decrease plasma ghrelin, the ‘hunger hormone’^69^. Short-chain fatty acids (SCFAs) are produced when gut microbes ferment non-digestible fibers, and these SCFAs bind to receptors in the colon, leading to the production of hormones that regulate food intake, body weight, and energy metabolism^7, 69^. Our findings from soil and T2D datasets of decreased potential metabolism of ubiquinone precursors (ubiquinone is central to metabolism and redox balance of cells across bacteria to vertebrates^70^), and molybdate (essential for nearly all life^71^) warrant further investigation. Additional compounds that we identified remain to be explored (Extended Data Table 3).

### Disentangling dietary and environmental influences

Altered potential metabolism for compounds detected within gut microbiomes may arise from diet, environmental exposures, gut environment and functions, neurological influences, among other factors^72^. The shared CPP trends in degradation and T2D (for example, BCFA-ACPs, sugars, lignin and precursors), are consistent with aligned environmental exposures and dietary inputs. Examples such as ammonia, with inconsistent CPP trends in T2D and a lack of apparent ecosystem quality trends, might be explained largely by diet. Similarly, increased potential metabolism for amylopectin in both T2D datasets could be related to higher consumption of starch-rich foods. For many compounds, we found alignment in potential metabolism between the ecosystem condition and T2D-CHN cohort, but not the T2D-SWE cohort – which may point to greater environmental exposures and detrimental health influence in some populations. Longitudinal and controlled experimental data are ultimately needed to discern environmental and diet influences in dynamic gut microbiomes – where health vs. disease may depend on thresholds and durations of compound-related exposures. Noteworthy implications for community connection to soils and ecosystems, and study limitations, are in Supplementary Material (Supporting Information).

## Conclusions

We found shared trends in the potential metabolism of compounds involved in energy harvesting and management between degraded soil microbiomes from five ecosystem quality gradients (from the United States, China, Australia) and gut microbiomes in T2D versus normal health (in cohorts from China and Sweden). Our results support existing studies implicating roles for microbiomes, sugars, lignin (dietary fiber) and mmBCFA in metabolism and T2D. Additionally, we raise the hypothesis that degraded soil-ecosystems may contribute to metabolic imbalances in T2D via nutrient-depleted food (i.e., low mmBCFA and lignin, high sugar) and/or microbial metabolite exposures that reshape gut microbiome functional capacities. Our method for examining microbiome potential metabolism at compound-scale raises biologically interpretable and testable hypotheses that could inform new microbiome-mediated T2D interventions.

## Methods

### Case study data

We examined previously published soil and human gut metagenome case study datasets (detailed in Extended Data Table 1). T2D datasets were from Swedish^37^ (T2D *n* = 33, Normal *n* = 43) and Chinese^36^ (T2D *n* = 30, Normal *n* = 52) cohorts. We used metadata from ref.^9^ to exclude subjects treated with metformin or diagnosed with prediabetes. From the Chinese cohort, we included only subjects >50 years old to compare with the mature-age (69-72 years) Swedish subjects. Australian Microbiome Initiative (AMI) disturbed versus natural soils were previously described in ref.^26^.

### Metagenome functional profiling via compound processing potential (CPP)

Our CPP analysis approach used a purpose-built bioinformatics workflow (Fig. 6) that redistributes values of functional potential relative abundances (%), from the scale of functions (i.e., biochemical pathways that often comprise multiple linked chemical reactions), down to the level of individual chemical compounds. Here, the concept of the collective functional capacity of a metagenome is transformed, without directional or thermodynamic constraints, into a hypothetical array of chemical reactions in which CPP captures the potential participation of compounds.

**Fig. 6.**
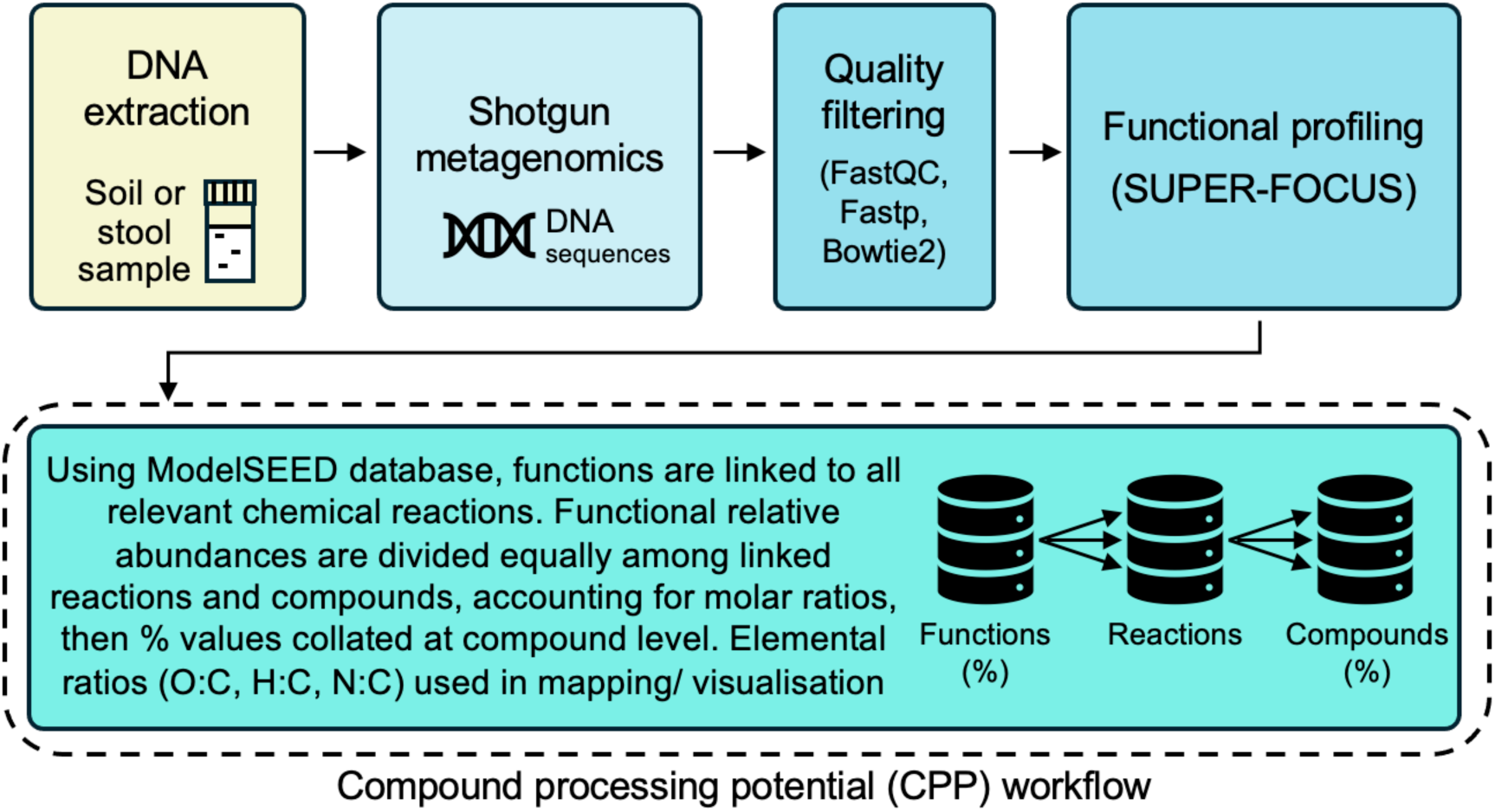
Overview of compound processing potential (CPP) workflow. Metagenomic functional potential pathway relative abundances were translated to quantify hypothetical potential metabolism for individual compounds. CPP % values for carbon-containing compounds can be mapped using chemical formula ratios to visualize clusters of bioenergetically similar compounds that trend together.

Measures of CPP were derived via steps outlined in Fig. 6. Firstly, following ref.^26^, using high performance computing resources^73^ raw metagenomic sequences were accessed, trimmed for quality control, and human case study sequences that mapped using Bowtie2^74^ to the human genome (GRCh38.p14/hg38) were excluded (low levels were observed; Supplementary Material, Supporting Information). Then good quality read 1 sequences were classified to SEED subsystem^75^ functional annotations and relative abundances using SUPER-FOCUS software^76^ (SUPER-FOCUS does not integrate paired reads, so R1-only were used for computational efficiency). Every SUPER-FOCUS function (output row) was translated to one or more corresponding chemical reaction(s) using a purpose-built R-script algorithm based on ModelSEED database lookup tables (from https://github.com/ModelSEED/ModelSEEDDatabase; accessed 10 Aug 2022). Linking of functions (and sub-functions where present) to chemical reactions was based on either: full matching of functional hierarchies (using subsystem-class,-subclass,-name and-role); detection of Enzyme Commission (EC) number; or matching of SUPER-FOCUS function name within ModelSEED lookup tables for reactions (reaction name or alias), subsystems (role), or reaction-pathways (external reaction name). Reactions were then linked to corresponding compounds using ModelSEED database tables (‘reactions.tsv’, ‘compounds.tsv’).

Advancing from the previous method used in ref.^26^, here we derived CPP measures at the resolution of individual compounds (all software parameters and supporting R code are available at: https://github.com/liddic/cpp_t2d):

(1) Function level relative abundances from SUPER-FOCUS (100% in each sample) were allocated to compounds by dividing values equally across all sub-functions and reactions, and dividing again across all corresponding compounds, with weightings to account for stoichiometric coefficients, and equal consideration to reactants and products.
(2) Relative abundances (%) for respective unique compounds (which often occur across multiple reactions and functions) were collated and summed to provide a summary CPP relative abundance (%) for each compound. Not all functions can be linked to chemical reactions, and conversion to CPP data format successfully recovered typical totals of 65-70%, which were retained on the same 0-100% scale, in the context of the initial 100% function potential per sample (Extended Data Table 1).
(3) For mapping visualization of CPP patterns for carbon-containing compounds, elemental ratios of O:C, H:C, and N:C were calculated. Such chemical mapping frameworks have been used to display and interpret groupings of energetically and chemically similar compounds^26, 40^.

## Statistical Analysis

We examined CPP trends in 30 example compounds, and from compound-wide trend analyses performed separately in each soil and T2D dataset, then looked for shared trends across the soil and T2D datasets. Further detail is in Supplementary Material and at https://github.com/liddic/cpp_t2d:

(1) ***Thirty example compounds*** were chosen due to their relevance to soil and/or gut health (Extended Data Table 2). CPP trends were evaluated using Kendall’s tau rank correlation tests based on ordinal age or ecosystem quality classes; or Wilcoxon-Mann-Whitney difference tests between disturbed vs. natural soils, and T2D versus normal subjects. We interpreted CPP trends with degradation as the inverse of calculated trends with increasing age or ecosystem quality. All data were plotted to assist interpretation.

(2) ***Compound-wide association tests*** were performed to exhaustively scan CPP trends for all compounds, using the same approach as above, with steps to automate analyses. In correlation tests, compounds that lacked relative abundance data for at least a quarter of the total sample size were excluded due to insufficient data. In Wilcoxon difference tests, compounds were only included if at least one or other of the groups contained at least 50% non-zero CPP values. Testing was based on log_10_-transformed CPP % values, with zero values replaced by small positive values equal to half the minimum non-zero CPP % across all relevant samples. One-sided (directional) difference tests were based on whether the median value in disturbed/T2D samples was greater or less than the median value for natural/normal samples. In the soil and T2D-CHN datasets, we used significant results based on Benjamini and Hochberg adjusted *P*-values. As described, for exploratory purposes, we retained results from unadjusted P-values (≤ 0.05) in the T2D-SWE dataset.

(3) ***Overlap analysis.*** We identified shared CPP trends that occurred simultaneously in 3 or more soil datasets, in T2D-CHN (*P*-adjusted) and T2D-SWE (unadjusted) results, with matches based on ModelSEED compound identifiers and directionality (increased/decreased), with an emphasis on degradation and T2D.

(4) ***Visualising compound-wide association test results*** Compounds with CPP trends identified separately in all soil and T2D datasets, and overlaps (i.e., shared trends as above), were mapped using their chemical formula elemental ratios of O:C, H:C, and N:C. From this mapping space, prominent localized groupings of consistently trending compounds (i.e., sugars, BCFA-ACPs) were observed and further analyzed to determine trends in their collective (summed) CPP % and to interpret their possible metabolic importance.

(5) ***Trend consolidation***. Trend and contextual information for compounds with shared trends were reported and visualized to represent mean correlations and significance (-log_10_[*P*-value]; as used in genome-wide association study Manhattan plots) of associations with degradation and T2D, numbers of functional themes (level 3 subsystems) they are linked to (for gut samples), and numbers of soil datasets showing trends.

(6) ***Beta diversity ordinations*** Supplementary analyses examined community-scale compositional differences in all datasets comparing conventional functional potential data versus CPP data (Supplementary Figs. 41-47; Fig. 1). This used principal coordinate analysis (PCoA) ordinations based on Bray-Curtis distances, and testing for differences between groups via permutational analysis of variance (PERMANOVA) and beta dispersion testing for homogeneity of groupings using the vegan R package^77^.

### Method validation and robustness to data processing

Supplementary analyses are provided to demonstrate our CPP method is robust to samples with varying sequence effort (within normal expectations for metagenome dataset quality), and that trend analysis signals are drawn widely from available compounds and not biased towards compounds that feature in many more reactions and functional pathways (Supplementary Material, Supporting Information, Figs. S48-54). As the CPP method is new, additional data analyses are provided for preliminary validation. We show the CPP method (1) recovers previously published gut microbiome functional beta diversity separation of mice raised in different soil environments; (2) tracks the dynamic response of soil microbiomes following glucose addition to soil; and (3) shows differentiation and clustering of diverse gut bacteria from single-species culture metagenome standards (Supplementary Material, Supporting Information, Figs. S55-57).

## Data availability

The datasets used are available from NCBI Sequence Read Archive (accessions PRJEB1786, PRJNA422434, PRJNA1215775, PRJNA1215778, PRJNA1215780, PRJNA1215781, PRJNA1080685), MG-RAST (project mgp16379), JGI IMG Study ID Gs0144357, and soil metagenomes from the Australian Microbiome Initiative data portal (https://data.bioplatforms.com/organization/australian-microbiome).

## Code availability

Code used to support this study is available at https://github.com/liddic/cpp_t2d.

## Acknowledgments

This work was supported by funding from the New Zealand Ministry of Business, Innovation and Employment (grant UOWX2101), and the Australian Research Council (DE260101877, LP240100073). We acknowledge and thank the authors of previously published metagenomics datasets used here. This includes the Australian Microbiome initiative, supported by funding from Bioplatforms Australia and the Integrated Marine Observing System (IMOS) through the Australian Government’s National Collaborative Research Infrastructure Strategy (NCRIS), Parks Australia through the Bush Blitz program funded by the Australian Government and BHP, and CSIRO. We acknowledge Indigenous peoples and the traditional custodians of the lands on which the case study datasets originate, and our authors live and work.

---

**Extended Data Table 1.** Description of case study metagenome datasets. **Notes for Extended Data Table S1:** *^a^*Ref.^9^ applied metformin treatment data in a reanalysis of metagenome data from Swedish^37^ and Chinese^36^ cohorts. We analyzed a subset of each cohort. From the Swedish cohort, we excluded subjects who were treated with metformin or diagnosed with impaired glucose tolerance (prediabetes). From the Chinese cohort, we only included subjects >50 years old to compare with the mature age (69-72 years) Swedish subjects. *^b^*AMI soils were surface depth (0–10 cm), from the temperate climate zone, with 7.5–45 % clay content (i.e., avoiding very sandy and very clayey soils) as previously studied in Ref.^26^. AMI Data portal URL: https://data.bioplatforms.com/organization/australian-microbiome.

| Case study | Main comparison groups (and sample/subject numbers) | Metagenome functional profile characteristics (sample mean ± s.d.) | Source data reference; sequencing platform | Study nation of origin |
| --- | --- | --- | --- | --- |
| **Human gut** |  |  |  |  |
| T2D-CHN | Chinese (CHN) cohort of subjects with T2D met- ($n = 30$; females $n = 17$, aged 51-71 years; males $n = 13$, aged 51-75 years). Normal ($n = 52$; females $n = 24$, aged 51-67 years; males $n = 28$, aged 52-74 years). Total $n = 82$. | <u>Functions</u> (SUPER-FOCUS): $n = 19,352$ ($8650 \pm 4214$). Functional % all sum to 100%. <u>Compounds</u> (CPP profiling): $n = 7,213$ ($5,311 \pm 1670$). CPP (%): ($65.7 \pm 4.1$ %) | SRA accession PRJNA422434<sup>9,36</sup><br><sup>a</sup>; Illumina Genome Analyzer II | China |
| T2D-SWE | Swedish (SWE) cohort of subjects with T2D without metformin treatment (T2D met-, $n = 33$), vs. normal healthy ($n = 43$). Subjects were females only, aged 69-72 years. Total $n = 76$. | <u>Functions</u> (SUPER-FOCUS): $n = 17,912$ ($7834 \pm 1699$). Functional % all sum to 100%. <u>Compounds</u> (CPP profiling): $n = 7,024$ ($5,287 \pm 585$). CPP (%): ($70.5 \pm 2.9$ %) | Sequence Read Archive (SRA) accession PRJEB1786<sup>9,37</sup> <sup>a</sup>;<br>Illumina HiSeq | Sweden |
| **Soil** |  |  |  |  |
| Post-mining restoration | Samples from a forest ecosystem restoration chronosequence span planting ages of 6, 12, 22, 31 years, and unmined (UM) samples ($n =$ groups of 3). Total $n = 15$. | <u>Functions</u> (SUPER-FOCUS): $n = 30,125$ ($20,328 \pm 767$). Functional % all sum to 100%. <u>Compounds</u> (CPP profiling): $n = 8,370$ ($7,777 \pm 92$). CPP (%): ($67 \pm 0.4$ %) | MG-RAST project mgp16379<sup>31</sup>;<br>Illumina HiSeq | United States |
| Disturbed vs. natural soils | Disturbed soils ($n = 29$) from agricultural land uses (cereals, cotton, horticulture, irrigated pasture) vs. natural soils ($n = 55$) from conservation and national park land uses. Total $n = 84$. From the Australian Microbiome Initiative (AMI). | <u>Functions</u> (SUPER-FOCUS): $n = 36,828$ ($22,158 \pm 1156$). Functional % all sum to 100%. <u>Compounds</u> (CPP profiling): $n = 8,529$ ($7,904 \pm 152$). CPP (%): ($66.4 \pm 1.2$ %) | AMI Data Portal [Data accessed Sep 2022]<sup>32</sup> <sup>b</sup>; Illumina HiSeq | Australia |
| Prairie restoration | Samples span unrestored agricultural fields (0 yr, $n = 6$), prairie grassland restoration (4, 5, 9, 17, 25, 30 yr, $n =$ groups of 3) and intact reference sites ($n = 6$). Total $n = 30$ | <u>Functions</u> (SUPER-FOCUS): $n = 35,926$ ($26,105 \pm 740$). Functional % all sum to 100%. <u>Compounds</u> (CPP profiling): $n = 8,527$ ($8,225 \pm 60$). CPP (%): ($67.0 \pm 0.3$ %) | JGI IMG Study ID Gs0144357<sup>33</sup>;<br>Illumina NovaSeq | United States |
| Plantation succession | From northeastern China, samples follow a gradient of *Pinus koraiensis* plantation succession from young, to half-mature, near-mature plantations, and reference old-growth mixed forest ($n =$ groups of 15). Total $n = 60$. | <u>Functions</u> (SUPER-FOCUS): $n = 36,406$ ($23,563 \pm 775$). Functional % all sum to 100%. <u>Compounds</u> (CPP profiling): $n = 8,493$ ($8,023 \pm 103$). CPP (%): ($66.0 \pm 0.5$ %) | SRA accessions<sup>34</sup><br>PRJNA1215775, PRJNA1215778,<br>PRJNA1215780, PRJNA1215781;<br>Illumina NovaSeq | China |
| Vegetation succession | From southwestern China, samples span successional stages of ecosystem recovery from abandoned farmland (1 yr), grassland ($\sim 5$ yr), shrubland ($\sim 20$ yr), shrub-arboreal mixed stage ($\sim 35$ yr), and arboreal stage ($\sim 50$ yr) ($n =$ groups of 3). Total $n = 15$. | <u>Functions</u> (SUPER-FOCUS): $n = 30,996$ ($20,564 \pm 1067$). Functional % all sum to 100%. <u>Compounds</u> (CPP profiling): $n = 8,349$ ($7,829 \pm 102$). CPP (%): ($66.3 \pm 0.4$ %) | SRA accession PRJNA1080685<sup>35</sup>;<br>Illumina HiSeq | China |

**Extended Data Table 2.** List of 30 example compounds relevant to soil and/or gut health and summary of trends in compound processing potential (CPP %), also visualized in Supplementary Figs. 2-31. Trends with ecosystem degradation are the inverse of Kendall’s tau correlation with vegetation age, or difference (Wilcoxon) due to disturbance. Shared trends across three or more soil-ecosystem datasets, or both type 2 diabetes (T2D) datasets are highlighted with red or blue shading. CHN = Chinese cohort. SWE = Swedish cohort. Trend symbols: **↑ = Increased**; **↓. = Decreased;** (↑) Marginally increased; (↓.) = Marginally decreased; - = No trend. Significance: *** = *P*-value < 0.001, ** = 0.001 ≤ *P*-value < 0.01, * = 0.01 ≤ *P*-value ≤ 0.05. Marginal trend indications have 0.5 < *P*-value < 0.1.

| Compound name | Trend with degradation |  |  |  |  | Trend with T2D |  | Compound description | Ref. |
| --- | --- | --- | --- | --- | --- | --- | --- | --- | --- |
|  | Post-mining restoration | Disturbed vs. natural | Prairie restoration | Plantation succession | Vegetation succession | T2D-CHN | T2D-SWE |  |  |
| D-Fructose | ↑ *** | ↑ *** | - | ↓ * | ↑ ** | ↑ ** | ↑ * | Dietary monosaccharide. Simple fruit sugar. | 78 |
| Sucrose | ↑ *** | ↑ *** | - | ↓ *** | ↑ ** | - | ↑ * | Disaccharide sugar, comprised of glucose and fructose | 78 |
| L-Arabinose | ↑ *** | ↑ *** | ↑ ** | - | ↑ *** | ↑ *** | ↑ * | L-arabinose sugar is a common component of plant biopolymers such as hemicellulose and pectin. | 79 |
| D-Glucose | ↑ *** | ↑ *** | $(\uparrow)$ | - | ↑ *** | ↑ ** | - | Dietary monosaccharide. Most abundant simple sugar found in blood, produced by plants. | 78 |
| Galactose | ↑ ** | ↑ *** | ↑ * | ↑ *** | ↑ ** | ↑ ** | - | Dietary monosaccharide. Found in lactose (combined with glucose), and in polymeric form in hemicellulose | 79 |
| Xylose | ↑ *** | ↑ *** | $(\uparrow)$ | $(\uparrow)$ | ↑ ** | ↑ ** | - | Monosaccharide sugar first isolated from wood. Major component of lignocellulose. | 79 |
| Mannose | ↑ ** | ↑ *** | ↑ * | ↑ *** | ↑ ** | ↑ *** | - | Monosaccharide sugar found in human metabolism made via digestion of polysaccharides & glycoproteins | 79 |
| Glucosamine | ↑ * | ↑ *** | ↑ *** | ↑ *** | ↑ * | ↓ ** | - | Abundant monosaccharide amino sugar, found in chitin | 79 |
| Chitin | ↑ *** | ↑ *** | - | ↑ ** | ↑ * | - | - | 2<sup>nd</sup> most abundant polysaccharide (after cellulose) main component of fungi cell walls, arthropod exoskeletons | 80 |
| Arabinoxylan (hemicellulose) | ↑ * | - | ↓ * | ↓ *** | - | - | - | Form of hemicellulose xylan found in plant cell walls. Main nonstarch polysaccharides in cereals and grasses | 80 |
| Hydrogen sulfide (H<sub>2</sub>S) | ↑ ** | ↑ *** | ↑ * | - | - | ↓ *** | - | Hydrogen sulfide is often produced from the microbial breakdown of organic matter in the absence of oxygen | 6 |
| Ammonia | $(\uparrow)$ | ↑ ** | - | ↑ *** | - | ↓ *** | ↑ ** | Compound of nitrogen and hydrogen | 81, 82 |
| Starch | - | $(\uparrow)$ | ↓ * | ↓ ** | - | - | ↑ * | Complex carbohydrate energy storage for plants, and major component in many human diets. Mainly comprised of amylopectin and amylose. | 78 |
| Amylopectin | - | ↓ *** | ↓ *** | ↓ *** | - | ↑ * | ↑ * | Highly branched polysaccharide polymer of α-glucose units found in plants. A major component of starch. | 78 |
| Amylose | - | ↑ * | ↓ ** | ↓ * | - | $(\uparrow)$ | ↑ * | Polysaccharide made of α-D-glucose units. Form of resistant starch due to tightly packed helical structure. | 78 |
| Glycogen | ↓ ** | ↓ *** | ↓ *** | ↓ *** | - | - | - | Multibranched polysaccharide of glucose that serves as a form of energy storage in animals, fungi, and bacteria | 83 |
| Acetate | ↓ *** | ↓ *** | ↓ ** | - | - | ↓ *** | (↑) | Short chain fatty acid produced by bacterial fermentation of carbohydrates | 6, 81 |
| Propionate | ↓ * | ↓ ** | ↓ *** | ↓ *** | ↓ * | ↓ *** | - | Short chain fatty acid produced by bacterial fermentation of carbohydrates | 6, 81 |
| Butyrate | ↓ ** | ↓ *** | ↓ *** | ↓ *** | ↓ *** | ↓ *** | - | Short chain fatty acid produced by bacterial fermentation of carbohydrates | 6, 81 |
| Methane | ↓ * | ↓ *** | ↓ *** | ↓ *** | ↑ ** | ↓ * | - | Formed via anaerobic respiration of carbohydrates. | 6 |
| Carbon dioxide | ↓ *** | ↓ *** | ↓ *** | ↓ *** | (↓) | ↓ *** | - | Primary product of microbial respiration when microbes break down organic matter to generate energy | 78 |
| Lignin | ↓ *** | ↓ *** | ↓ *** | ↓ *** | - | ↓ * | ↓ * | Class of complex organic polymers. Structural component in plants. Dietary fiber. | 80 |
| Trimethylamine N-oxide (TMAO) | ↓ ** | ↓ *** | ↓ ** | ↓ *** | ↓ ** | - | ↓ ** | Amine oxide that tends to increase with consumption of animal proteins. | 6, 81 |
| Trimethylamine (TMA) | ↓ ** | ↓ *** | ↓ ** | ↓ *** | ↓ ** | - | (↓) | Trimethylated derivative of ammonia synthesized by gut(-like) bacteria from dietary animal products | 6 |
| Pectin | (↑) | ↑ ** | - | - | - | (↓) | ↓ * | Soluble dietary fiber (polysaccharide) from plant cell walls, common in citrus fruits. | 80 |
| P-cresol | - | ↓ *** | ↓ *** | ↓ * | - | ↓ * | (↑) | p-Cresol is produced by bacterial fermentation of protein | 6 |
| Menaquinone (Vitamin K2) | (↓) | ↑ * | ↑ ** | - | - | ↓ ** | - | An electron carrier involved in anaerobic ATP-generating redox reactions. Also important in anaerobic synthesis of pyrimidines, porphyrins and succinyl CoA | 6 |
| Serotonin | (↑) | (↓) | ↓ * | - | - | ↓ * | (↓) | Key neurotransmitter produced by gut EEC cells | 84 |
| Cellulose | - | - | - | - | ↑ *** | - | - | Primary structural component in plants. Dietary fiber. | 78 |
| Indole | - | ↑ ** | ↑ ** | (↑) | - | ↓ *** | - | Aromatic amine, naturally in human feces. | 81 |

**Extended Data Table 3.**
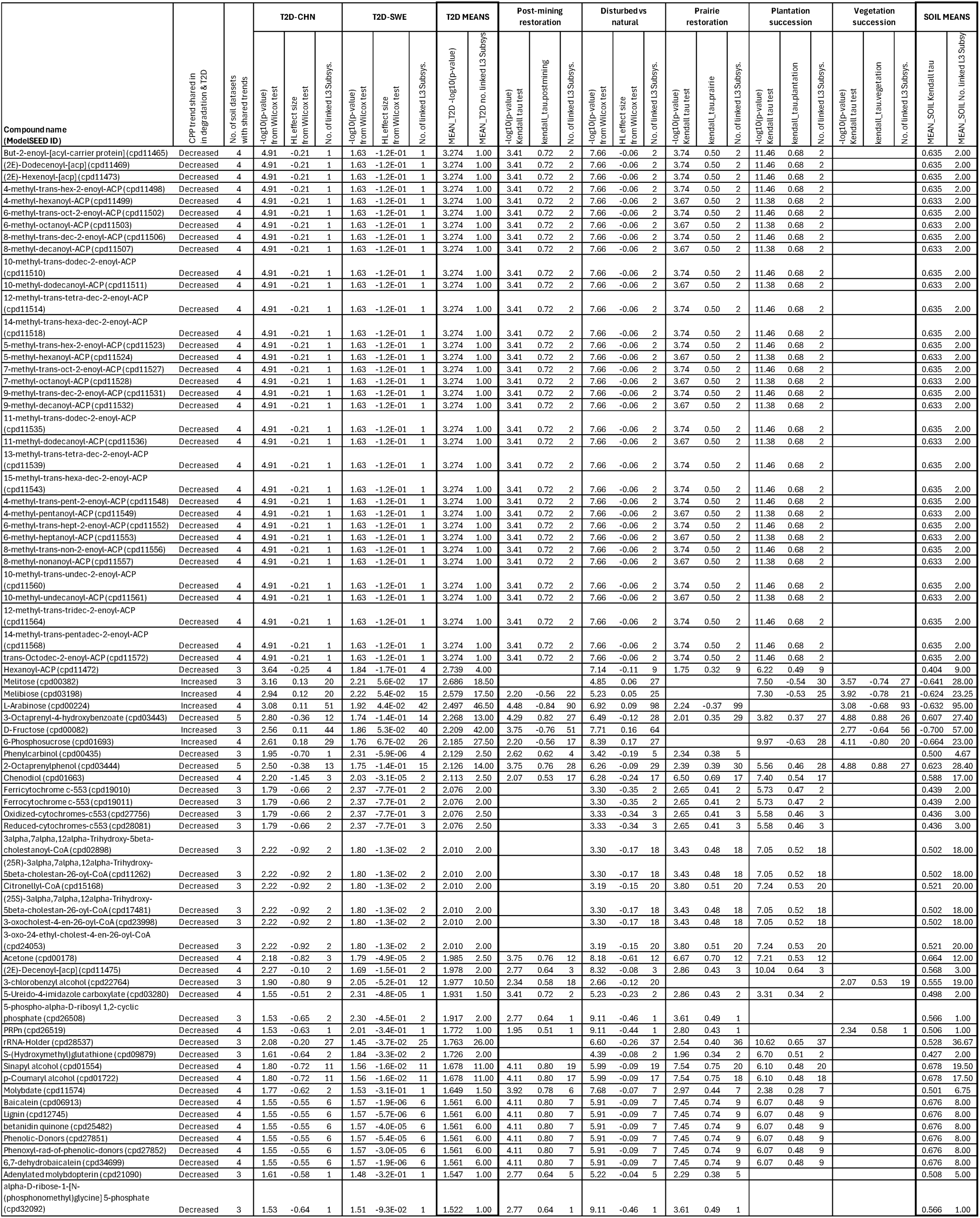
Compounds with shared *P*-adjusted directional CPP trends with degradation in at least 3 of 5 soil datasets, *P*-adjusted trends with T2D in Chinese (CHN), and unadjusted trends in Swedish (SWE) cohorts. Mean soil and T2D data, and numbers of soil datasets with shared trends are used in the main article Fig. 3.

**Extended Data Table 4.**
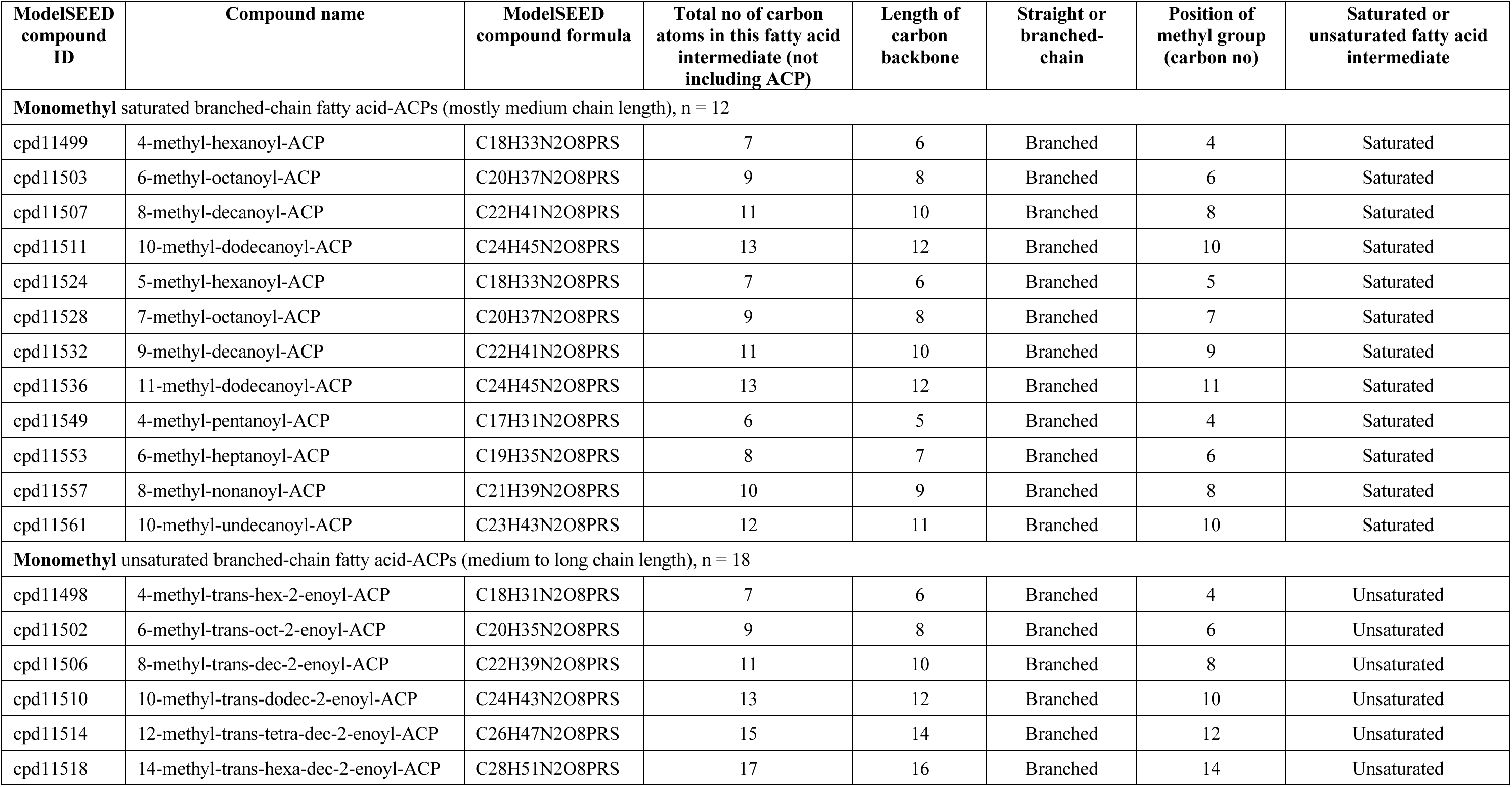
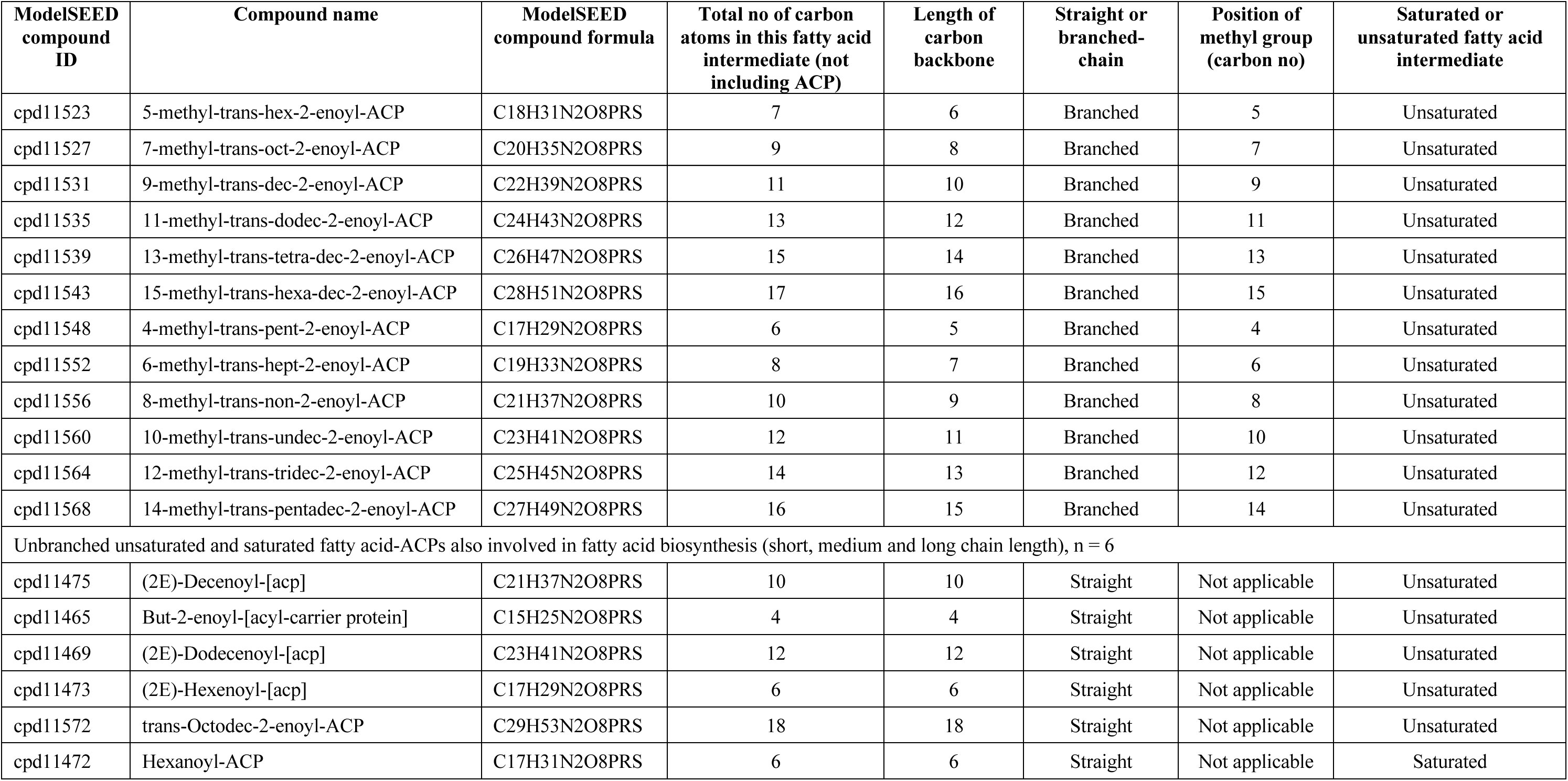
List of 36 fatty acid-acyl carrier protein (ACP) compounds that showed consistent trends of decreased potential metabolism in Chinese (*P*-adjusted) and Swedish (unadjusted) type 2 diabetes cohorts, and in 3 or more degraded soil-ecosystem datasets. These are dominated by monomethyl branched-chain fatty acid-acyl carrier proteins (BCFA-ACPs). Based on the length of the main carbon chain, BCFA are divided into short-chain branched fatty acids (< 6 carbon atoms), medium-chain branched fatty acids (6-12 carbon atoms), long-chain branched fatty acids (13-21 carbon atoms) ^47^. Note: the chemical formula for ACP compounds used in the ModelSEED database system, may vary from other systems (e.g., KEGG compound database) due to ModelSEED’s representation of the phosphopantetheine arm of the ACP, which contains 11 carbon atoms.

