## Supplementary Material for "Shared potential metabolism trends in degraded soils and type 2 diabetes gut microbiomes"

**This PDF file includes:**

Supplementary Information
Figures S1 to S57
Tables S1
Supplementary References

**Other supporting materials for this manuscript (available: https://github.com/liddic/cpp_t2d):**

Code used to support this study
Output files:
- ‘1-CPP-trend-results-Post-mining-restoration-soils-Jun-2026.xlsx’
- ‘2-CPP-trend-results-Disturbed-vs-natural-soils-Jun-2026.xlsx’
- ‘3-CPP-trend-results-Prairie-restoration-soils-Jun-2026.xlsx’
- ‘4-CPP-trend-results-Plantation-succession-soils-Jun-2026.xlsx’
- ‘5-CPP-trend-results-Vegetation-succession-soils-Jun-2026.xlsx’
- ‘6-CPP-trend-results-T2D-CHN-gut-Jun-2026.xlsx’
- ‘7-CPP-trend-results-T2D-SWE-gut-Jun-2026.xlsx’
- ‘8-Shared-CPP-trends-among-soil-datasets.xlxs’
- ‘9-Shared-CPP-trends-among-soil-and-T2D-datasets.xlsx’

### Supporting Information

***Examination of the BCFA-ACPs compound group***

The group of compounds named BCFA-ACPs (after branched-chain fatty acid – acyl-carrier proteins) in Fig. 2b was given this label as it is dominated by this type of compound ($n$ = 30), together with $n$ = 6 straight-chain fatty acid-ACPs (Extended Data Table 4). All 36 compounds in the BCFA-ACPs group are involved in fatty acid biosynthesis and they each occur on two straight lines in the O:C, H:C, N:C mapping space (line 1: H:C = -0.625 × O:C + 2, and line 2: H:C = -0.375 × O:C + 2, where N:C is in the range >0 to 0.2; Fig. S40). The BCFA-ACPs group comprised three categories of compounds: monomethyl unsaturated medium to long BCFA-ACPs ($n$ = 18), monomethyl saturated predominantly medium BCFA-ACPs ($n$ = 12), and unsaturated or saturated short, medium, and long straight-chain fatty acid-ACPs ($n$ = 6) (Extended Data Table 4). BCFA lengths are based on the main carbon chain (short: < 6, medium: 6-12, long: 13-21 carbon atoms; Ref.[1]).

As monomethyl BCFA-ACPs dominated this trend group, we were curious to understand whether they might feature there exclusively, or if similar compounds were also present throughout the data and not display trends. From a search across all soil and T2D datasets used in the study, we found $n$ = 295 other unique ACP-containing compounds, beyond the 36 compounds highlighted in the BCFA-ACPs group. However, when focusing only on the methyl-containing variants in the other 295 ACP-containing compounds they were different in character to the BCFA-ACPs we highlighted. Specifically, there were no other monomethyl unsaturated (-enoyl) BCFA in the datasets beyond those that featured in our BCFA-ACPs trend group. The other non-trending methyl-containing ACP compounds ($n$ = 56) either contained carbonyl (-oxo-) groups ($n$ = 18), or hydroxyl (-hydroxy-) groups ($n$ = 22), or ester groups ($n$ = 9), while the remaining ($n$ = 7) were saturated BCFA-ACPs (one short chain, and six long 13-16 chain length). These latter, non-trending saturated BCFA-ACPs also differed from the saturated BCFA-ACPs in our trend group, which contained mostly medium length chains (Extended Data Table 4).

***Implications for community connection to soils and ecosystems***

Our findings suggest that filling knowledge gaps in the pathogenesis of T2D may require more holistic consideration of linkages, especially at the systems level of health risk and promotion. For example, where landscape-scale ecosystem management may drive changes in functional capacities of health-supporting microbiomes and units of interest may span scales from microbial community-scale activity to constituent compounds of microbial bodies and necromass. Our findings suggest that maintaining balanced functioning healthy soil ecosystems in human habitats, especially those used for food production, may be increasingly important for supporting healthy human populations. Increasing urbanization, ecosystem degradation, the industrialization of food production and its distribution, and reducing time spent in nature have collectively reduced contact with soil microbiomes for a large proportion of people. When humans are exposed to soil in modern environments, it is increasingly within urbanized or highly modified landscapes (for example, degraded ecosystems with low natural biodiversity, highly modified land and disturbed soils, monocultural agricultural fields) where microbiomes and their functional capacities have been greatly altered.

From a societal viewpoint, our findings suggest that addressing the global T2D pandemic may benefit from restoring urban biodiversity, developing effective approaches for regenerative agriculture, increasing soil biological health and diversity as key parts of a restorative continuum, along with finding a better balance between anthropogenic land use and nature. At the heart of this challenge lies the need for a deeper understanding of soil health—recognizing soil as a complex living ecosystem in which the microbiome plays a pivotal role. Understanding the critical links between soil health, food production, and human health is fundamental for strengthening the diminished connections between people, land and soil[2] and provides a foundation for developing strategies and actions that might lead to improved gut health and resilience to T2D. This critical understanding of links between soil-ecosystem health and human health also draws on parallels of thinking about nature from an Indigenous peoples' worldview. Indigenous knowledge offers guidance and support to theories that place the health or condition of nature (land, biodiversity, water and air) as essential to maintaining human health and wellbeing. Central concepts also recognize the importance of reconnecting with nature by strengthening engagement with ecosystems through activities such as restoration and regeneration. For example, in Aotearoa-New Zealand, Indigenous Māori stress connection and inter-dependency with ecosystems through core principles such as ancestral links and association[3]. This includes concepts of whakapapa (genealogical descent of all living things) which holistically reaffirms the importance of genetic and chemical assemblages in soils and landscapes to support all life forms, including microbiomes[4]. Concepts of kaitiakitanga (guardianship of the environment) are also commonly used to reinforce actions and strategies to guide environmental restoration and promote health. Indigenous perspectives offer important guidance to all peoples for personally reconnecting and engaging with our ecosystems and their stewardship in order to support healthy landscapes, soils, microbiomes and people.

***Study limitations***

There is well documented regional geographic variation in soil and gut microbiomes[5, 6, 7, 8] which may affect the interpretation of our results. We have examined a limited number of case studies to demonstrate a proof-of-concept, because relevant case study metagenome datasets (with publicly available metadata that span gradients of ecosystem quality, or provide consistent comparisons between untreated T2D and healthy control subjects with relevant medication histories, while limiting uncontrolled sources of variation) are rare. Although, we note that land uses and management that degrade ecosystems may be driving homogenization and hence greater predictability of soil microbiome structure and functions especially in urban areas[9, 10]. There are also limitations with assessing ecosystem degradation-restoration gradients from chronosequence (space for time proxy) study designs, and comparing T2D gut microbiomes versus healthy controls from cross-sectional (snap-shot in time) data. For soil-ecosystems, existing evidence suggests they may recover over decades from disturbed states towards reference community structures with restoration[11]. Ideally, longitudinal and experimentally controlled data are needed to study causal shifts in both soil and gut microbiomes. That said, our CPP method aims to bypass variation that is often inherent with taxonomically focused studies, via targeting the hypothetical potential metabolism of compounds, based on the idea that microbiomes will be universally shaped by the resources and the environmental conditions available to them. It remains to be tested whether the patterns we observed hold in other geographic regions and human populations, and whether they may be replicated under controlled experimental conditions. The SUPER-FOCUS and ModelSEED resources we used have not been actively updated in recent years and may be missing the latest functional annotations. There may be some inaccuracies in the mapping from SUPER-FOCUS functions to ModelSEED pathways and reactions, however we use rule-based scripts that can be refined, consistent with other software development. We recognize this CPP method represents a simplification of complex metabolic processes within microbiomes – for example, it does not discern anabolism, catabolism, reactants or products. Moving beyond this proof-of-concept, future work should use updated resources that enable comprehensive linkage from functional pathways to reaction and compound databases. Noting the conceptual difference in our approach (CPP reflects functional potential imprinted in metagenomic DNA), we suggest future work should compare CPP measures to quantitative measurement of compounds present (for example, via metabolomics). Control samples were not included in the case study datasets analysed, however all samples are considered 'high biomass' with low risk of contamination, and we removed human genome sequences (observed at low levels) from gut metagenome samples.

***Reporting of human genome sequence removal from T2D gut metagenome case study datasets***
The Swedish cohort data contained only low levels (median 0.03 %, interquartile range (IQR) 0.01 - 0.08 %) of trimmed reads that were classified as human sequences. After host-removal, median 7,820,878 (IQR: 5,572,690 - 12,868,662) read 1 sequences were used for SUPER-FOCUS functional profiling. The Chinese cohort data contained even less human sequence data (median 0.0008 %, IQR 0.0003 - 0.002 % of trimmed reads), leaving median 12,700,702 (IQR: 10,224,381 - 19,183,919) non-host read 1 sequences for SUPER-FOCUS functional profiling.

***Searching for compounds linked to gut health***
We searched for oligosaccharides (for example, fructan, inulin, galactan) but did not detect these compounds in our datasets. Also, the following gut hormones were sought but not found in the datasets: cholecystokinin (CCK), secretin, somatostatin, motilin, ghrelin, glucagon-like peptide 1 (GLP1), glucose-dependent insulinotropic peptide (GIP), insulin-like peptide 5 (INSL5), peptide YY (PYY), gastrin, neurotensin, growth differentiation factor 15 (GDF15), fibroblast growth factor 19 (FGF19), guanylin, uroguanylin and oxyntomodulin.

***Robustness to data processing***
To show that our CPP method is not biased towards detecting signals in more highly 'connected' compounds (i.e., that feature in many more reactions and functional pathways), we visualized the frequency distribution of compounds with trending CPP in terms of their number of linked level 3 subsystems in the post-mining restoration dataset. This distribution follows a similar pattern to all compounds in the dataset indicating that trends are drawn widely from less connected to more connected compounds (Fig. S48).

To address uneven sequencing effort, SUPER-FOCUS normalizes function counts by dividing by the total annotated reads per sample (https://github.com/metageni/SUPER-FOCUS#run). Moreover, we used relative abundance format functional potential outputs as the basis for CPP values. To check that our method is robust to samples with varying sequencing effort (which are routinely encountered in metagenomics pipelines) we checked for correlations between the main signals (BCFA-ACPs, sugars, lignin and precursors) and sequence counts across all datasets (Fig. S49). There were no correlations with sequence counts in the soil datasets. However, correlations were detected in T2D-CHN dataset with BCFA-ACPs and sugars, and in T2D-SWE with sugars only. Rarefying data is used in some bioinformatic pipelines especially with a focus on diversity comparisons[12], however this approach is criticized[13] for throwing out data and samples to meet rarefaction requirements. To provide reassurance that our CPP method would produce reliable results, without undue influence from variation in sequencing effort, we performed re-analyses from the stage of cleaned (host-removed) DNA sequences in both the T2D-CHN and T2D-SWE datasets with rarefying of sequences and associated exclusion of samples at the minimum library size, 5th, 10th, 15th and 20th percentile sequence counts. It should be noted that these scenarios effectively created different data subsets often with large amounts of data and some samples discarded. Reassuringly, key signals generally persisted when datasets were reanalyzed with the rarefaction to even read depths, yielding 25 out of 30 results (83%) with $P \leq 0.05$ (Figs. S50-54; results including samples and sequence counts are provided in Table S1).

***Supplementary data analyses for preliminary validation of the CPP method***

The CPP method is new, therefore supplementary example datasets and analyses are provided for preliminary validation purposes. <u>However, caution is recommended to avoid over-interpretation of any results, as we expect the CPP approach to be most useful for hypothesis-generation purposes.</u> Here, we demonstrate the ability of the CPP method to: (1) recapture previously published functional differentiation in gut metagenomes of mice raised in different soil environments; (2) show dynamic response of CPP for glucose in soil metagenomes following glucose amendment to soil; and (3) show differentiation and clustering of different gut bacteria from single strain culture metagenome standards.

In Figs. S55-57, we explore each dataset providing plots and statistical tests for apparent differences between groups (Kruskal-Wallis) or trends (Kendall tau correlations) for CPP values for selected example compounds (e.g., glucose, cellulose, $CO_2$, $O_2$, $H_2O$). We also provide beta diversity ordinations for samples (based on Bray-Curtis distances), and a heat map for scaled $\log_{10}$(CPP) showing compounds with above 50$^{th}$ percentile variation in CPP values. We also explore two metabolic indices based on CPP values: (a) adenylate energy charge index, AEC = (ATP + 0.5xADP)/(ATP + ADP + AMP) which has been used to indicate metabolic vitality of cells[14]; and (b) adenosine triphosphate/ adenosine diphosphate (ATP/ADP) ratio, which can indicate the balance in capacity for anabolism or catabolism (cells need to maintain an adequate ATP/ADP ratio to drive biological reactions)[15].

<u>Supplementary analysis #1: Mice gut metagenomes raised in different soil environments</u>

Liu et al.[16] (their Fig. 2) show gut microbiomes from mice raised in desert, grassland (steppe) or forest soils develop markedly different functional beta diversity groupings. In our beta diversity ordination plot using CPP data (Fig. S55a), we recapture similarly prominent functional groupings based on soil exposures, including the position of the forest soil exposure group intermediate between grassland and desert groups. Our PCoA plot achieved a comparable level of variance explained (69.8 + 21.6 = 91.4 %), compared to Liu and colleagues' original analysis based on KEGG functions (80.7 + 11.7 = 92.4 %). Interestingly, CPP values for $H_2O$ in the desert soil-exposed group are significantly higher than both grassland and forest groups (Kruskal-Wallis, $P = 1.6 \times 10^{-15}$). CPP-based values of ATP/ADP ratio are also substantially lower in desert soil group compared to others (Kruskal-Wallis, $P = 6.1 \times 10^{-13}$), which may reflect microbial communities that are prepared for destructive metabolism and decomposition of available organic material. These signals for CPP-$H_2O$ and ATP/ADP in the desert soil group seem consistent with influence from desert biocrust associated microbial communities that are activated following rare and short periods of rain, and adaptation to limited productivity[17]. Raw metagenome data were obtained from NCBI Sequence read archive, accession: PRJNA542998.

Supplementary analysis #2: Soil metagenome response to glucose amendment

Chuckran et al.[18] published soil metagenomes that followed glucose amendment in an agricultural soil, with samples taken at time intervals of 0h, 8h, 24h and 48h ($n$ = groups of 3) after glucose addition. Our CPP-based analysis of this data shows distinct time-based groupings in the beta diversity ordination (Fig. S56a), and a dynamic rise and peak, with a subsequent levelling out of CPP for glucose (Fig. S56c) in the soil microbiome samples. (We note that Chuckran and colleagues discarded three metagenome samples due to heavy degradation, and we also excluded sample C0D3 due to outlying beta diversity.) Raw metagenome data were obtained from NCBI Sequence read archive, accessions[18]: PRJNA539715, PRJNA539712, PRJNA539720, PRJNA539713, PRJNA539717, PRJNA539718, PRJNA539719, PRJNA539721, PRJNA539722, PRJNA539723, PRJNA539714, PRJNA539711, PRJNA539716.

Supplementary analysis #3: Gut bacteria single-strain culture metagenome standards

Amos et al.[19] published metagenome standards for the National Institute for Biological Standards and Control (NIBSC) based on single-species cultures for a diverse range of gut bacteria. We visualized this data in CPP data format (Fig. S57) showing differentiation and clustering of the different gut bacteria, with examples that align with expected variation in traits of various species. For example, in the beta diversity ordination space (Fig. S57a), species such as *Escherichia coli* (fast-growing, opportunist, facultative anaerobe) and *Bacteroides thetaiotaomicron* (having wide metabolic flexibility) were distributed far apart from others, while more closely related *Bifidobacterium longum subsp. longum* and *Bifidobacterium longum subsp. infantis* had only a small level of separation. As expected, *E. Coli* (facultative anaerobe) featured with high levels of CPP for oxygen ($O_2$). *Clostridium butyricum* and *E. coli* featured with low levels of ATP/ADP ratio, and high levels of AEC index which may reflect their capacity for degrading substrates to release energy. Raw metagenome data were obtained from NCBI Sequence read archive, accession: PRJNA622674.

**(a)**

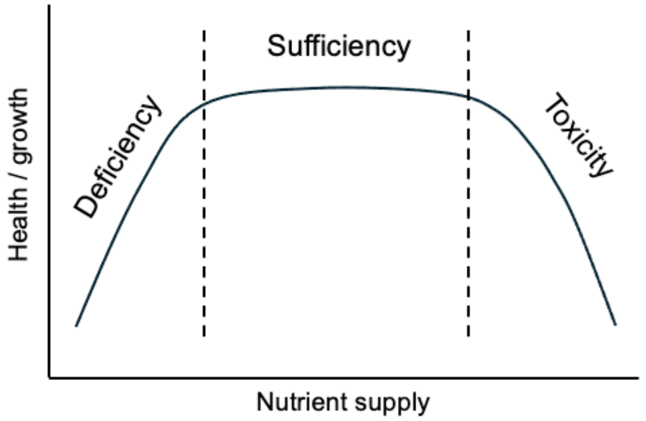


**(b)**

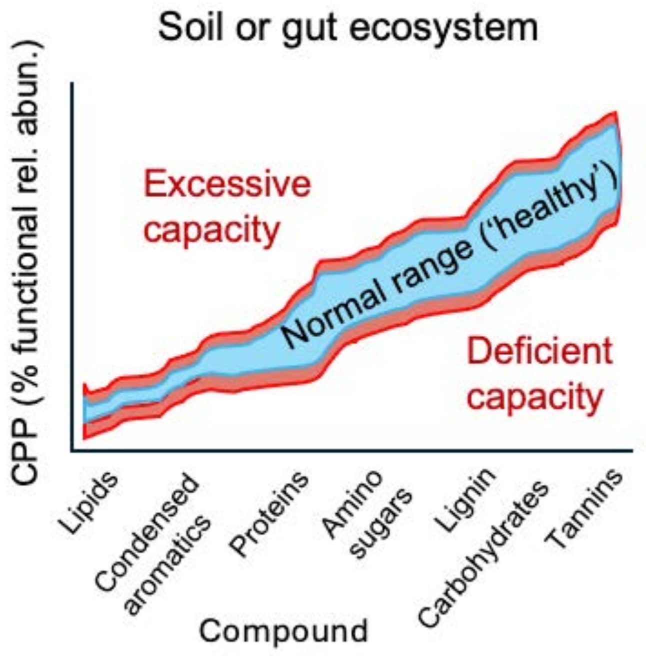


**Fig. S1.** Conceptual illustration of how compound processing potential (CPP) values might be used to quantify normal 'healthy' states with a normal range of functional capacity to process various compounds, versus 'unhealthy' states with deficient or excessive functional capacity. **(a)** Non-linear, u-shaped dose-health response relationships (e.g., deficiency-sufficiency-toxicity) are generalizable across biological systems[20]. **(b)** In the context of CPP, this conceptual approach suggests that, for any given compound, normal health might be associated with a moderate range of CPP values, while poor/abnormal health might be associated with deficient or excessive CPP values. In soils, such a quantitative concept of soil health – assessed via distributions of CPP values – could align with recognised concepts of 'soil capability' (i.e., potential upper range of functionality of a reference soil-ecosystem type, estimated from many samples) and 'soil condition' (current state of one particular sample)[21]. CPP measures reflect the integrated response of dynamic bioindicator microbiomes and variation is expected across different distinctive ecosystem types.

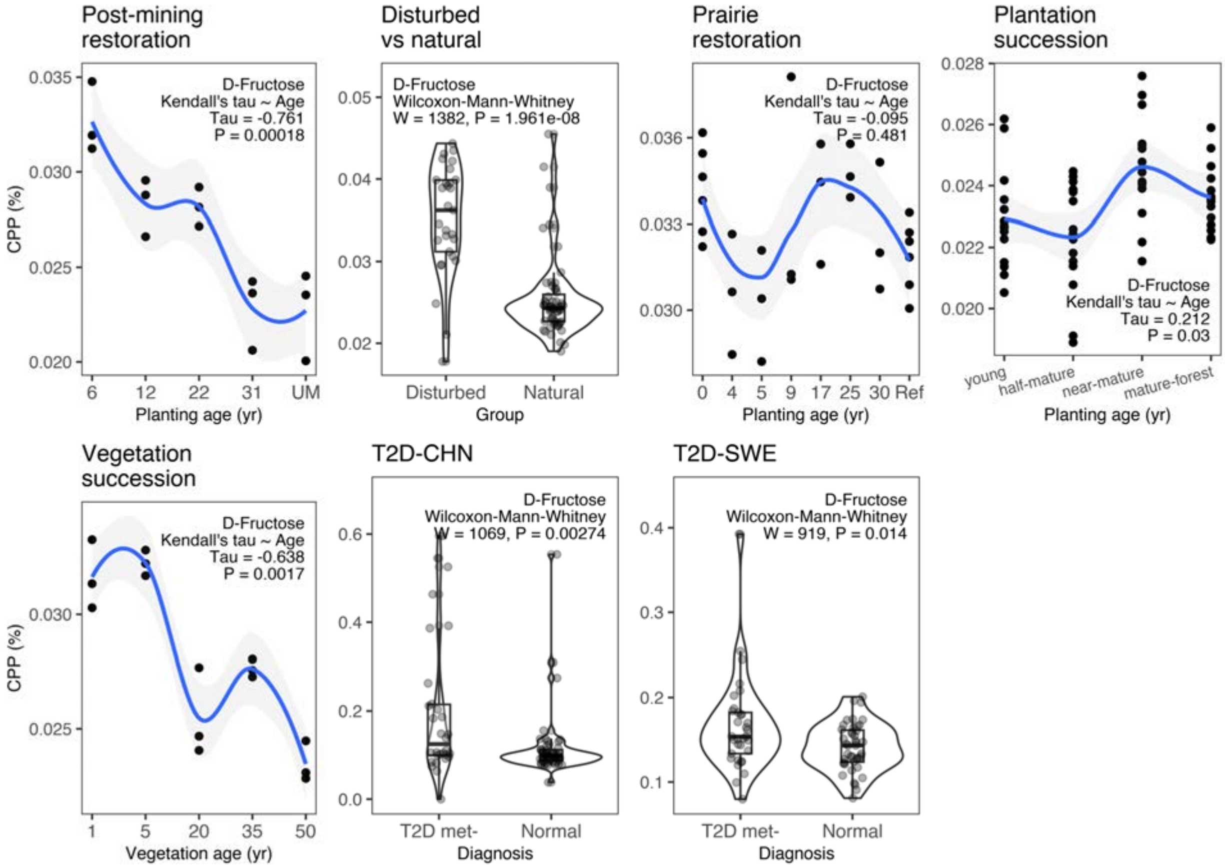


**Fig. S2.** Comparison of CPP values for **fructose** (dataset details in Extended Data Table 1).

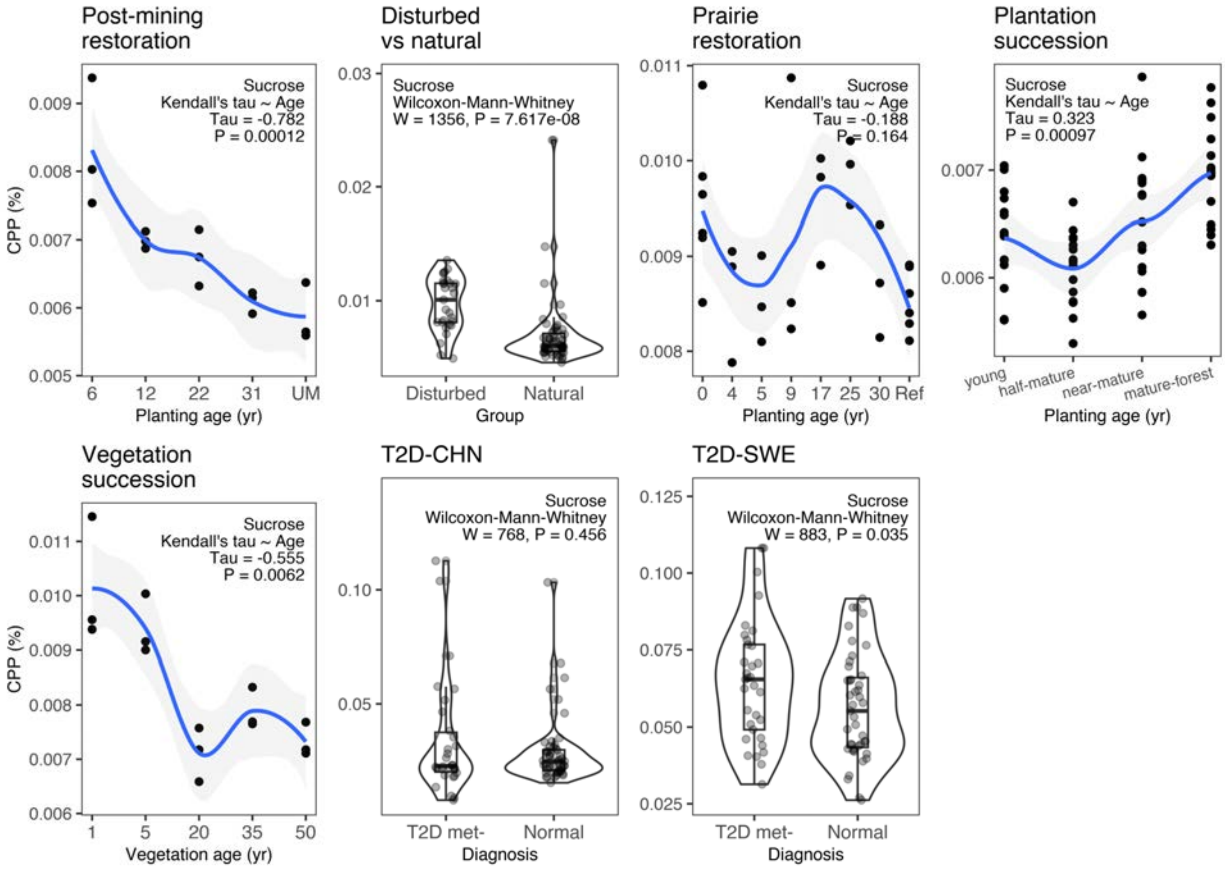


**Fig. S3.** Comparison of CPP values for **sucrose** (dataset details in Extended Data Table 1).

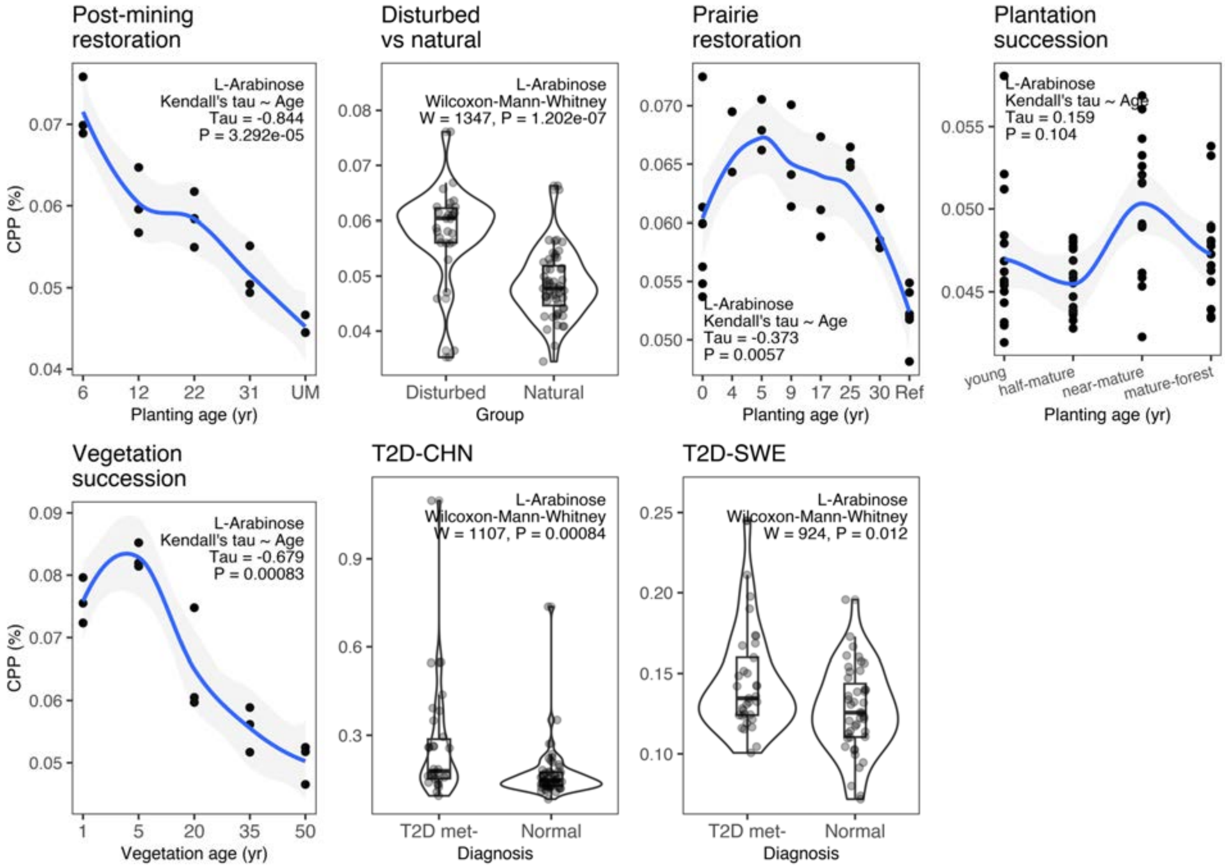


**Fig. S4.** Comparison of CPP values for **arabinose** (dataset details in Extended Data Table 1).

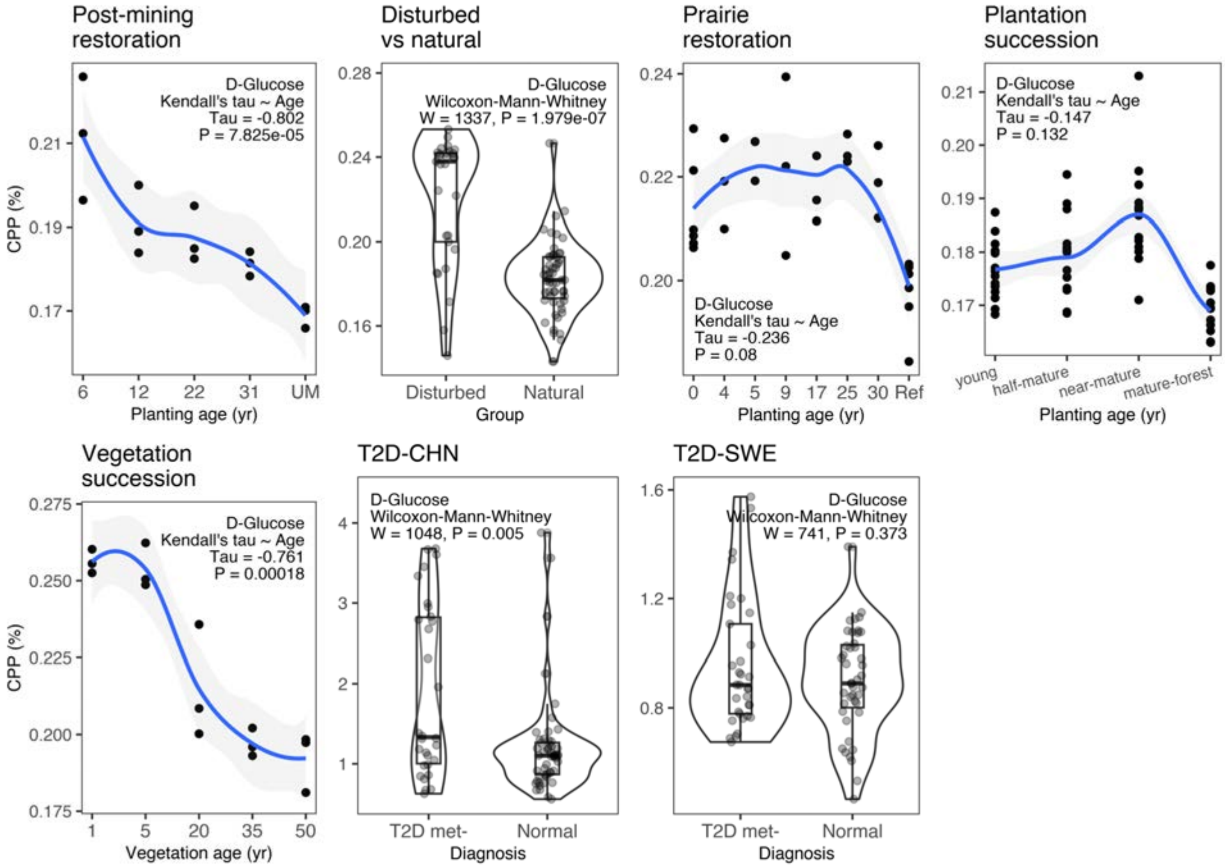


**Fig. S5.** Comparison of CPP values for **glucose** (dataset details in Extended Data Table 1).

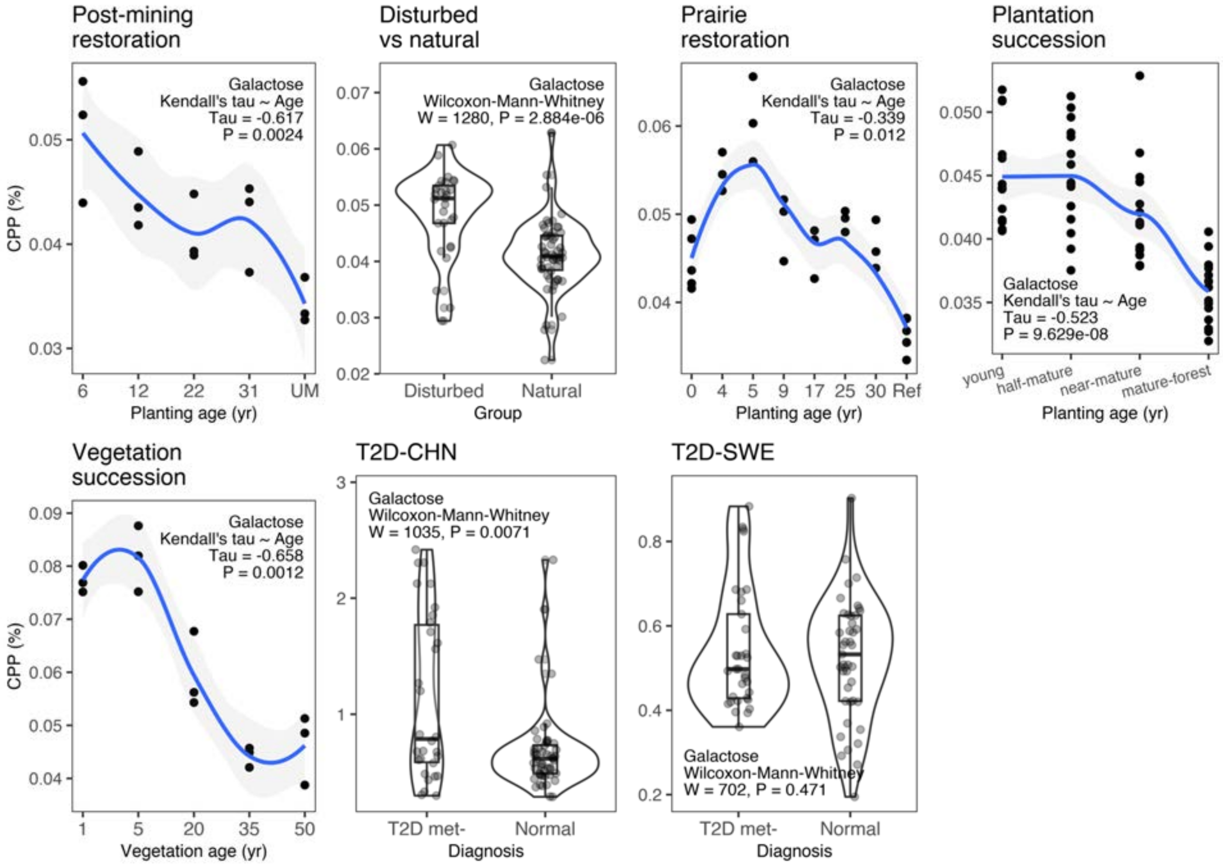


**Fig. S6.** Comparison of CPP values for **galactose**. (dataset details in Extended Data Table 1).

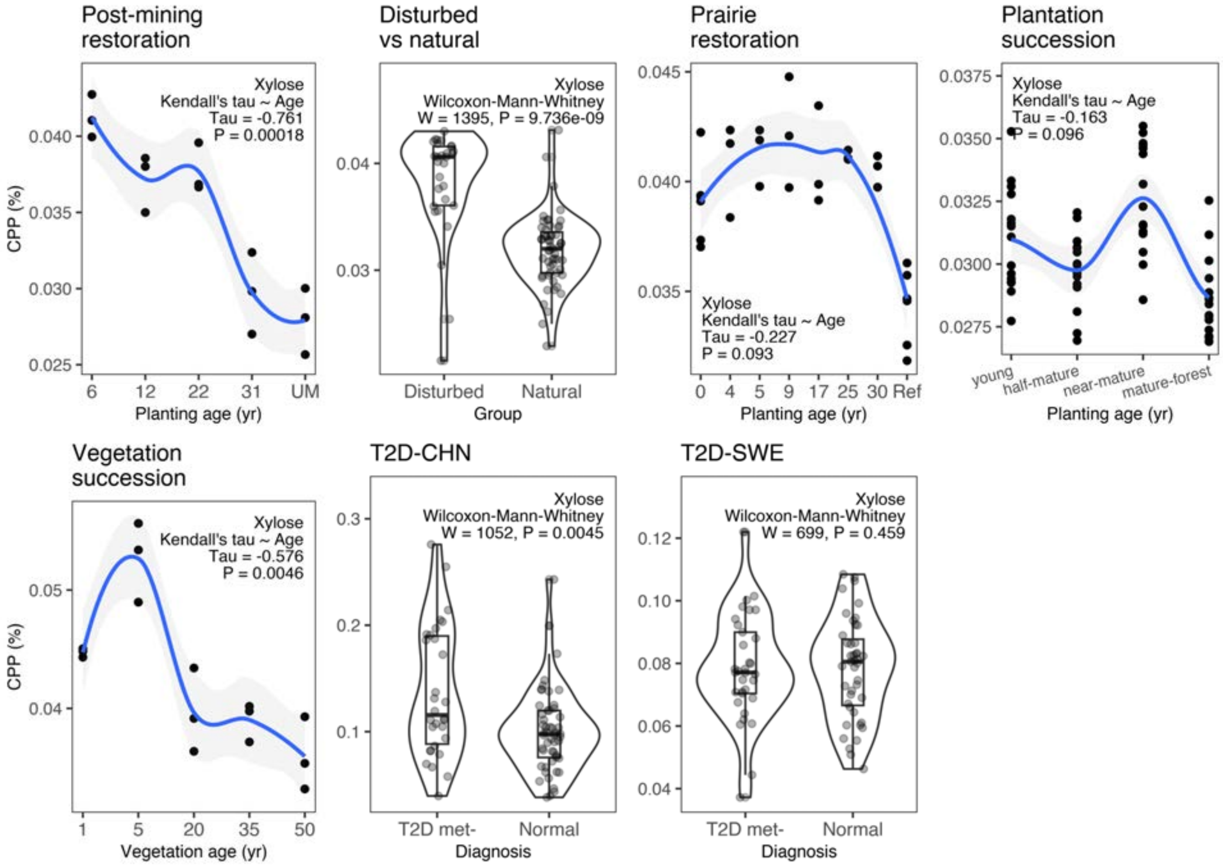


**Fig. S7.** Comparison of CPP values for **xylose** (dataset details in Extended Data Table 1).

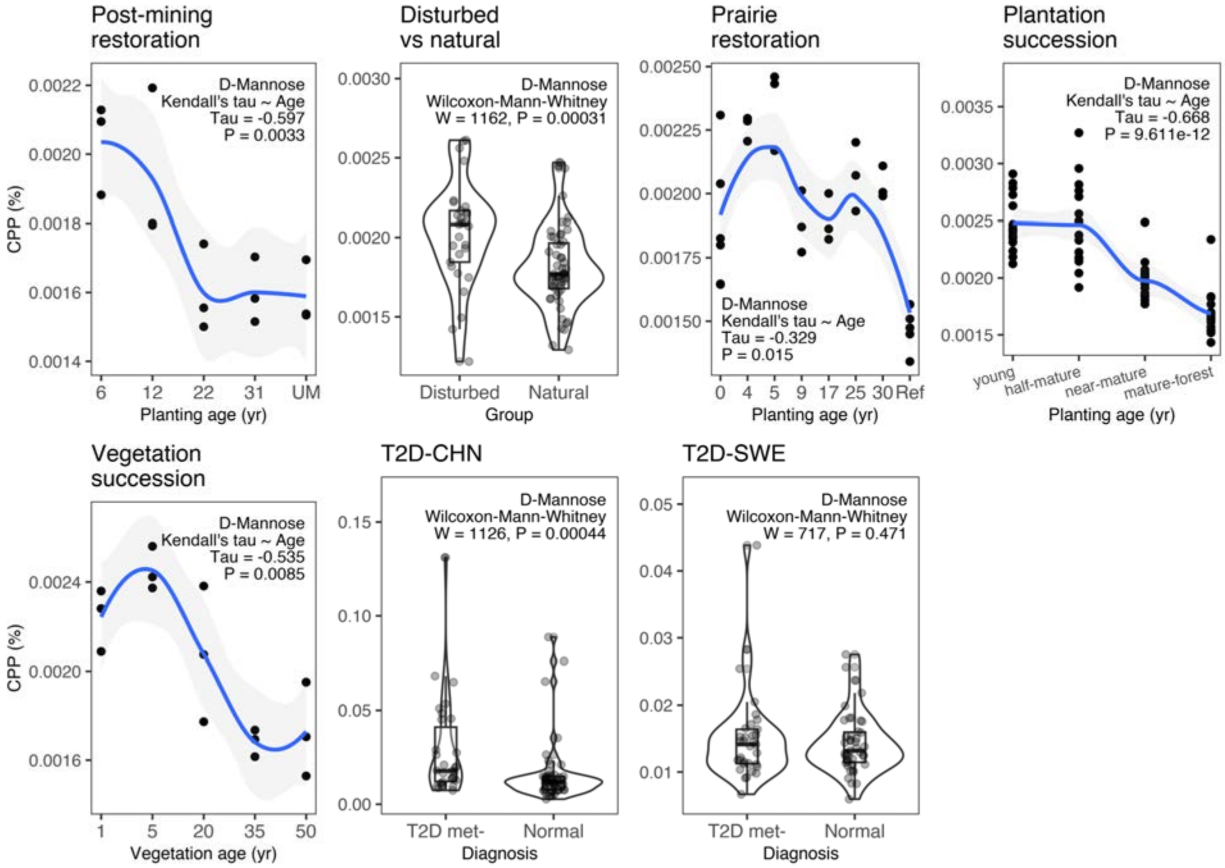


**Fig. S8.** Comparison of CPP values for **mannose** (dataset details in Extended Data Table 1).

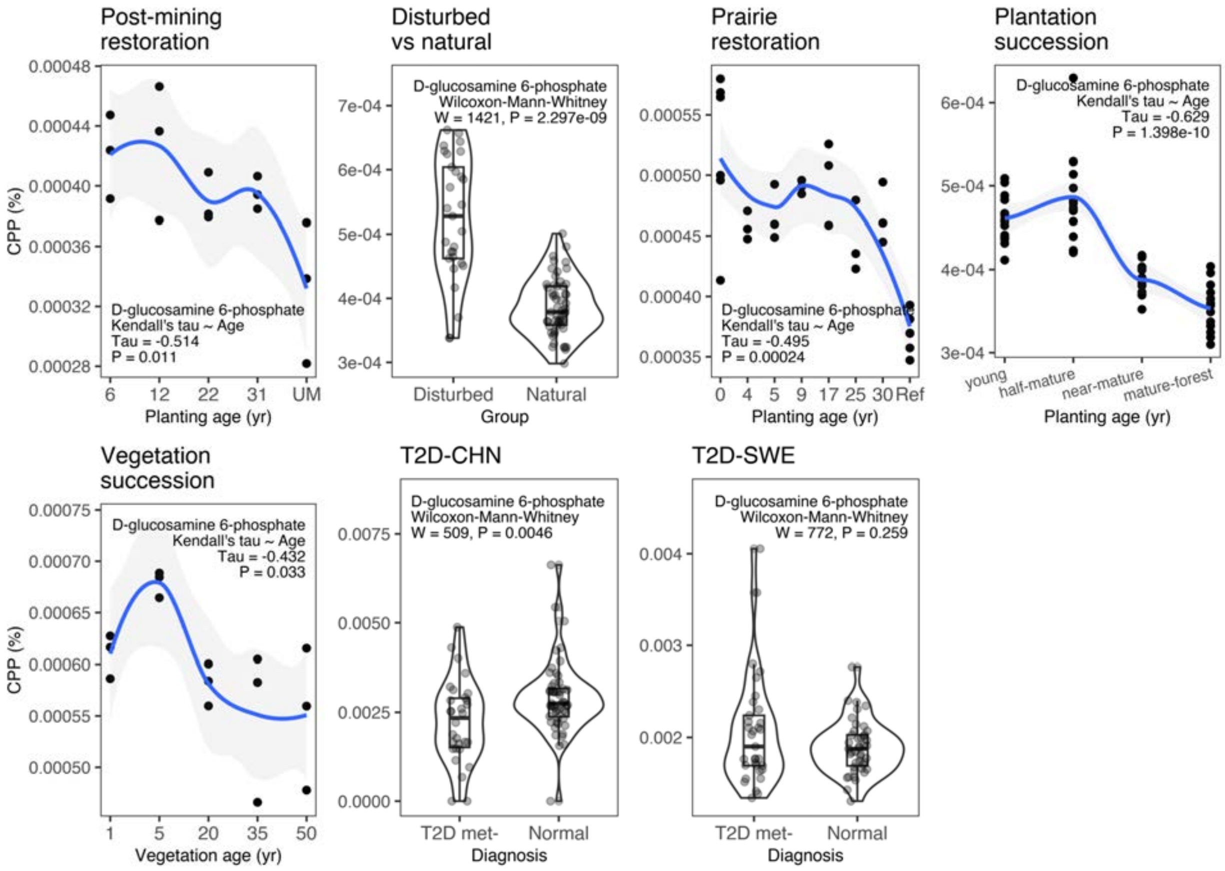


**Fig. S9.** Comparison of CPP values for **glucosamine** as D-glucosamine 6-phosphate (dataset details in Extended Data Table 1).

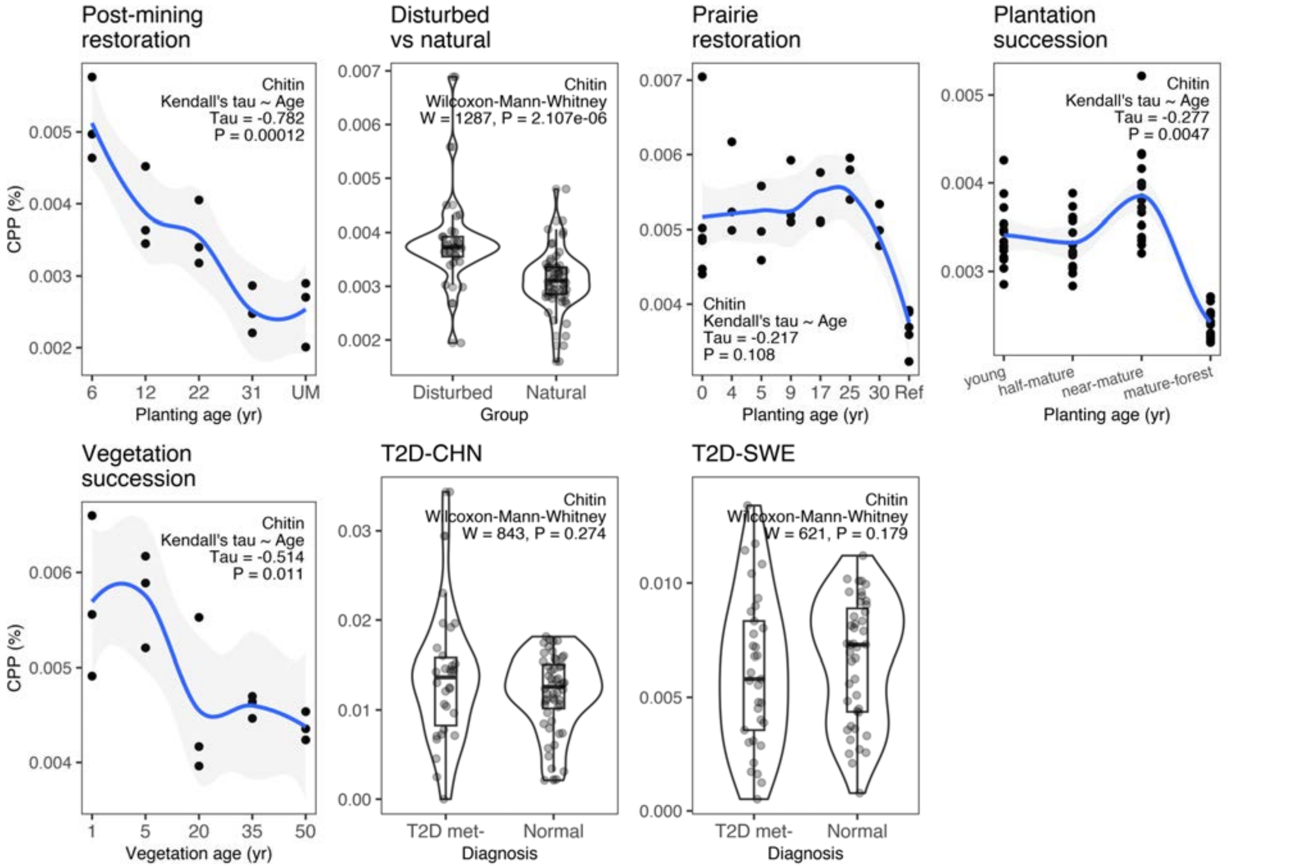


**Fig. S10.** Comparison of CPP values for **chitin** (dataset details in Extended Data Table 1).

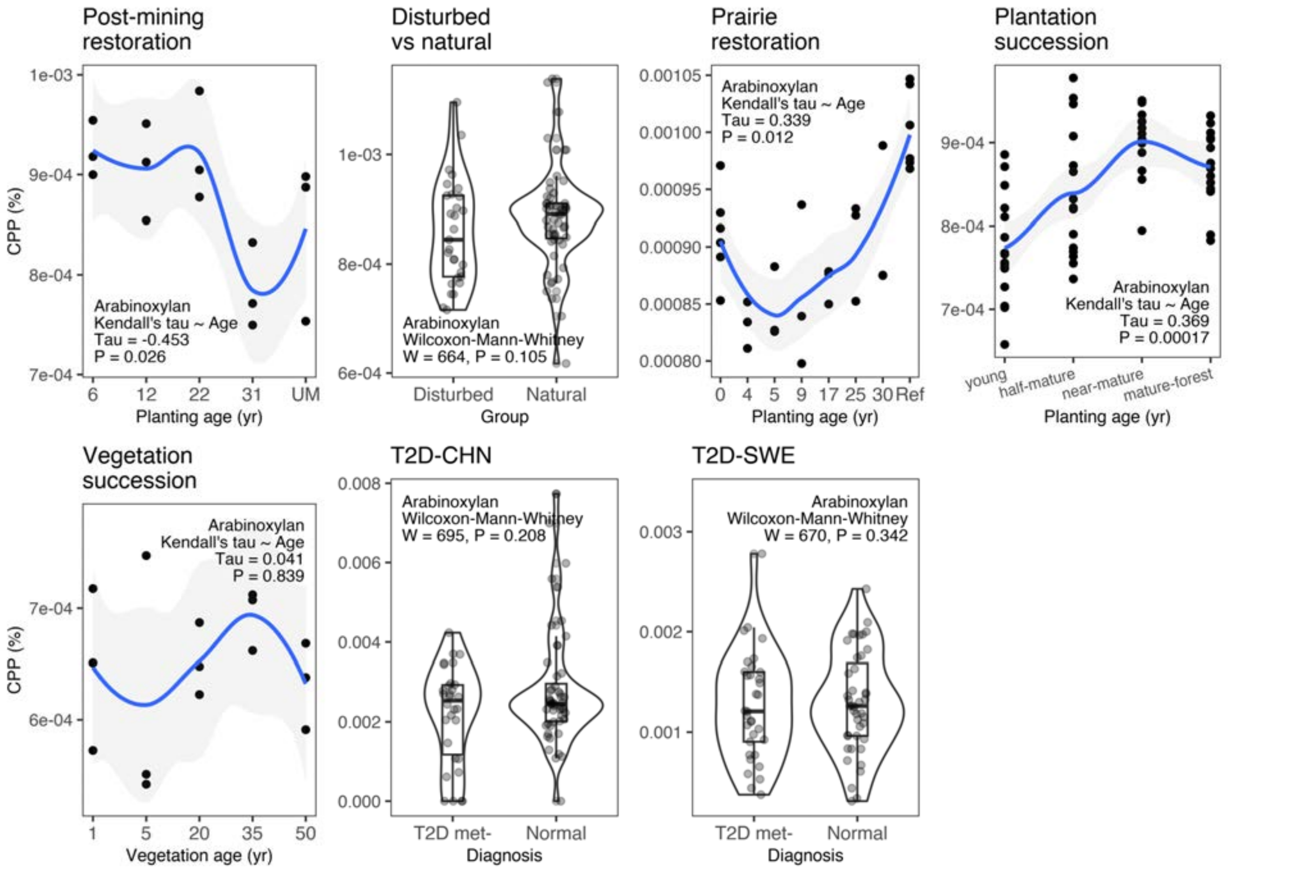


**Fig. S11.** Comparison of CPP values for **arabinoxylan,** a form of hemicellulose (dataset details in Extended Data Table 1).

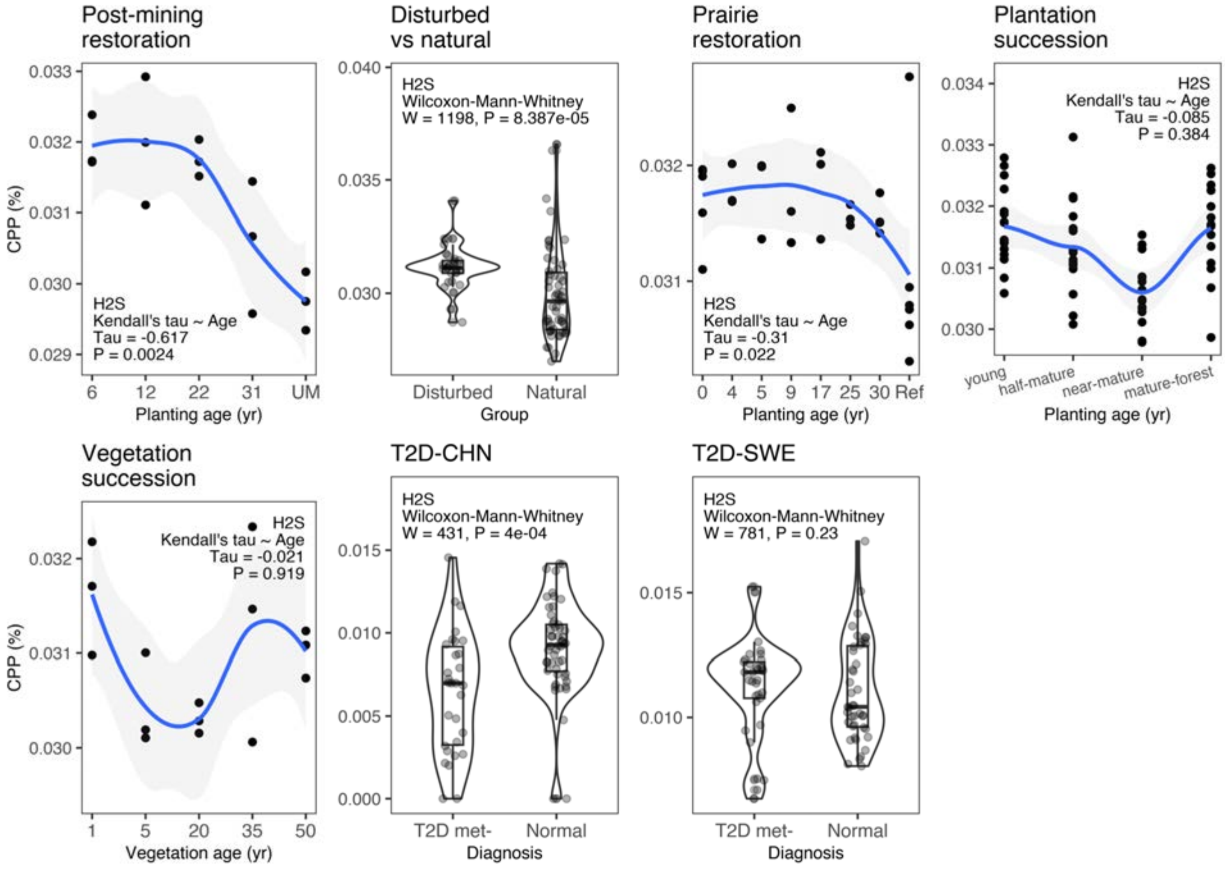


**Fig. S12.** Comparison of CPP values for **hydrogen sulfide** ($H_2S$) (dataset details in Extended Data Table 1).

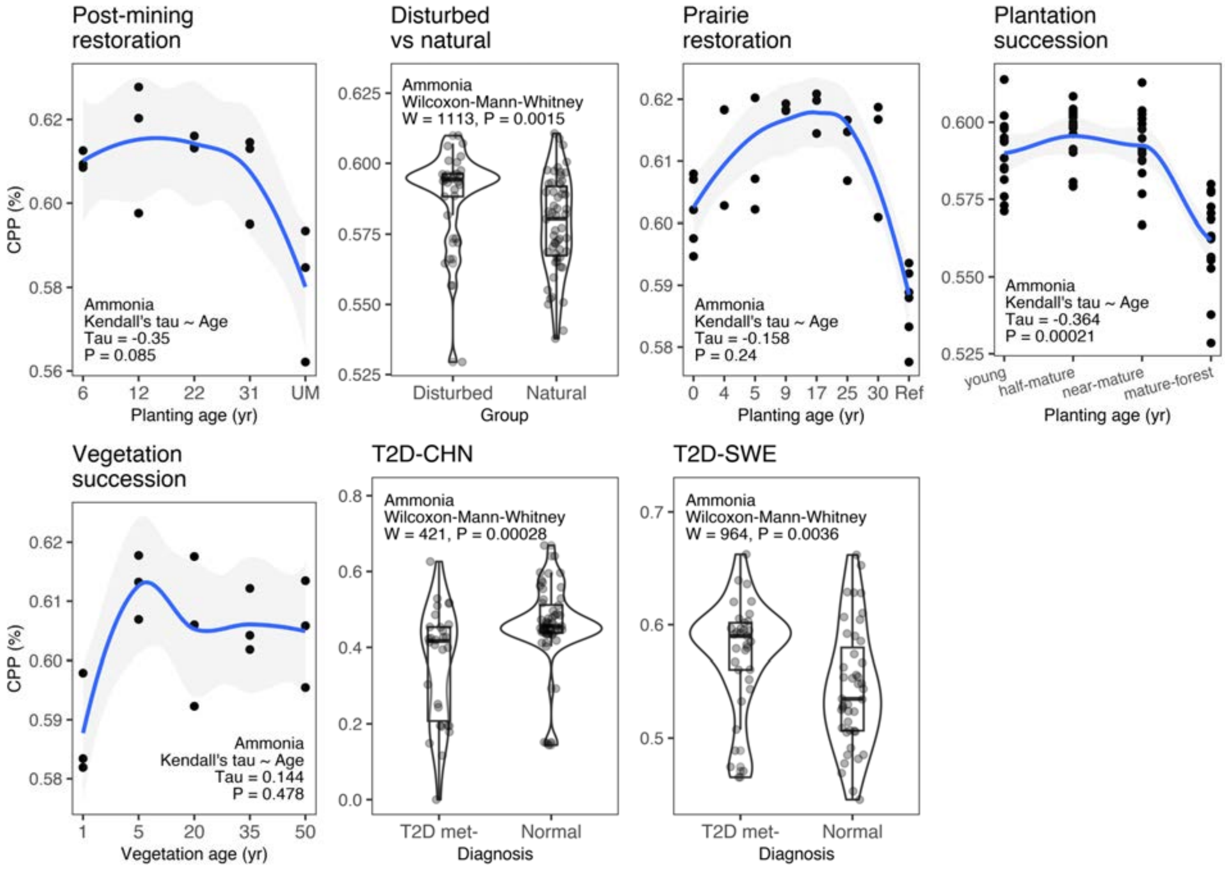


**Fig. S13.** Comparison of CPP values for **ammonia** (dataset details in Extended Data Table 1).

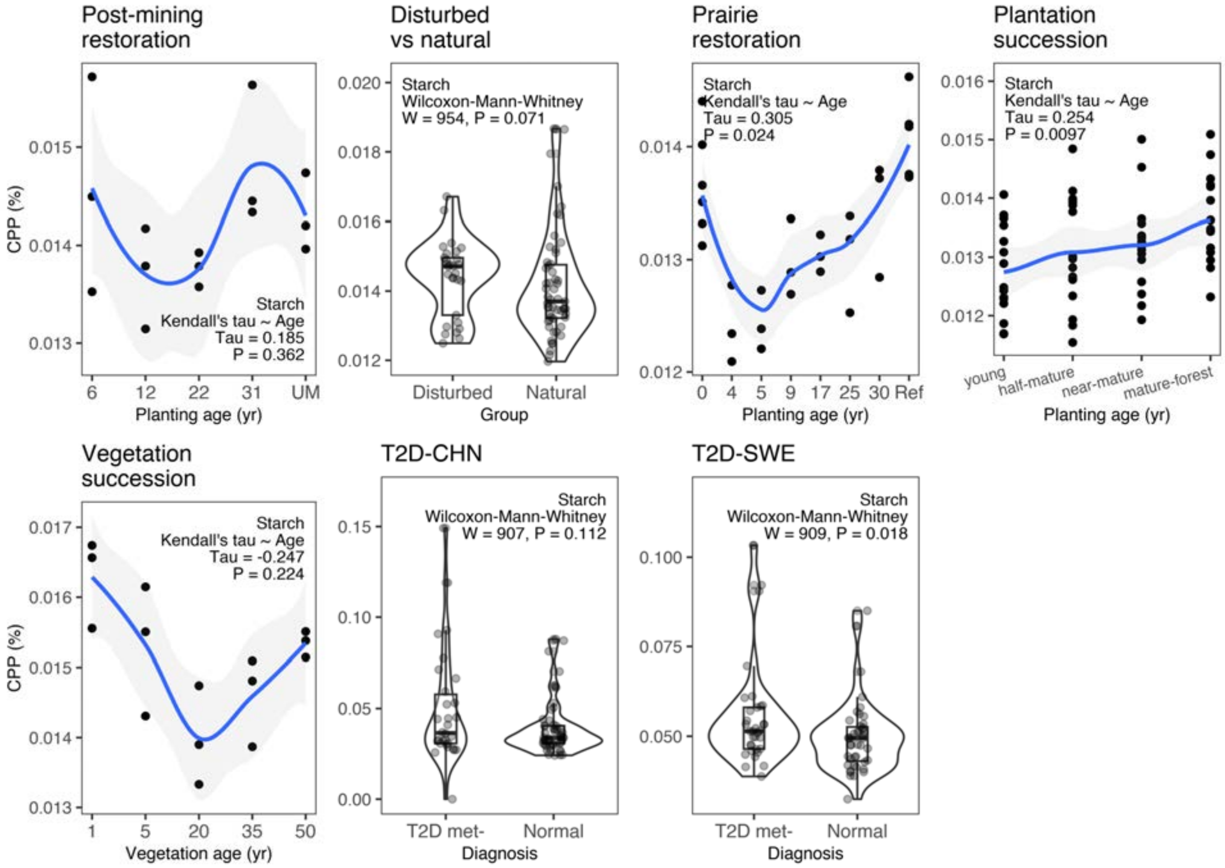


**Fig. S14.** Comparison of CPP values for **starch** (dataset details in Extended Data Table 1).

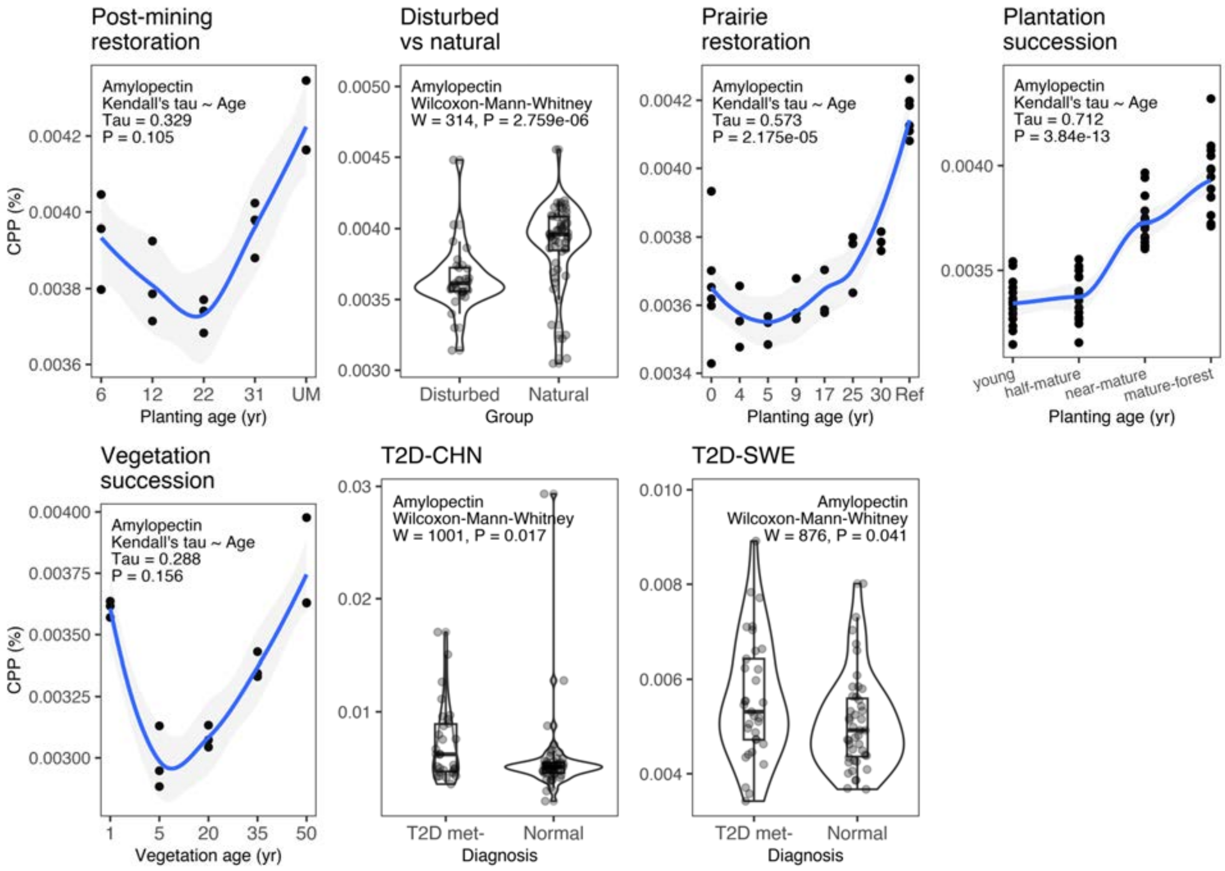


**Fig. S15.** Comparison of CPP values for **amylopectin**, a component of starch (dataset details in Extended Data Table 1).

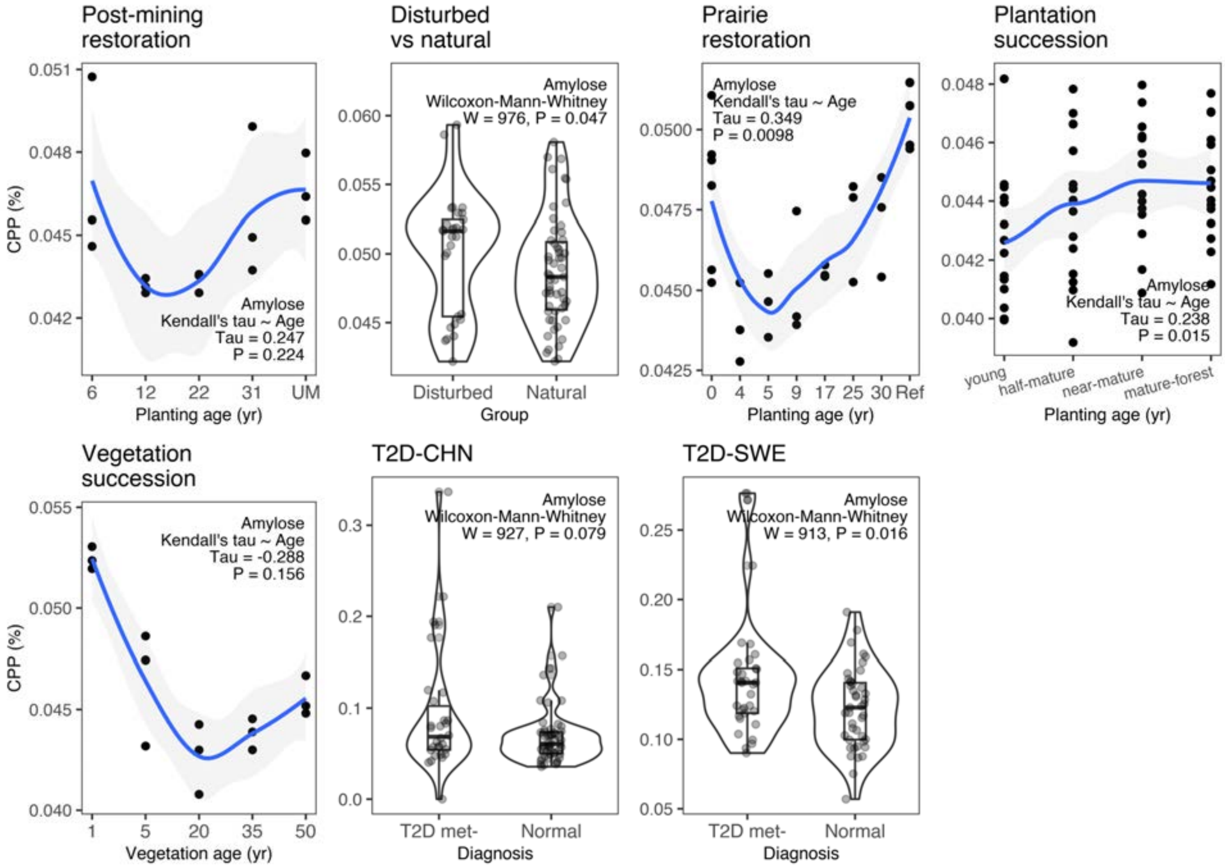


**Fig. S16.** Comparison of CPP values for **amylose**, a component of starch (dataset details in Extended Data Table 1).

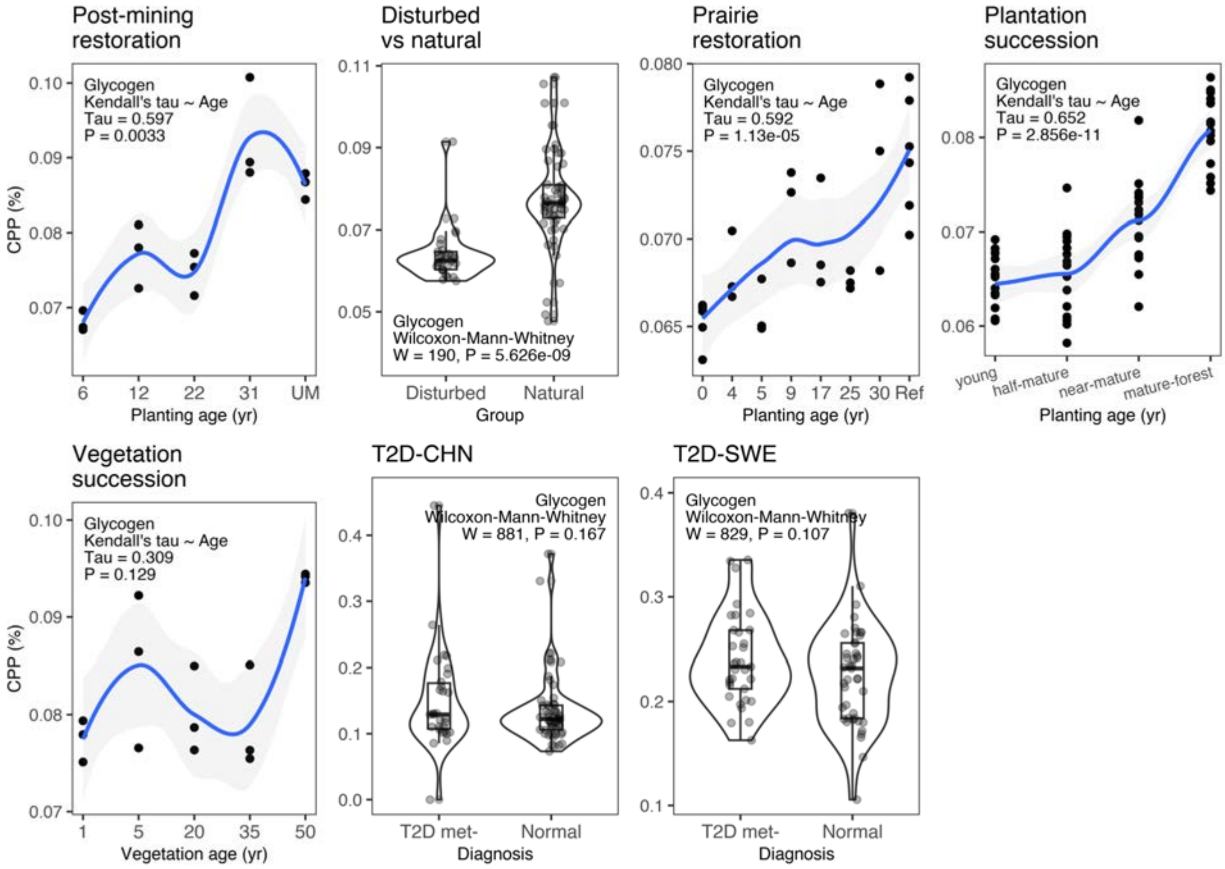


**Fig. S17.** Comparison of CPP values for **glycogen** (dataset details in Extended Data Table 1).

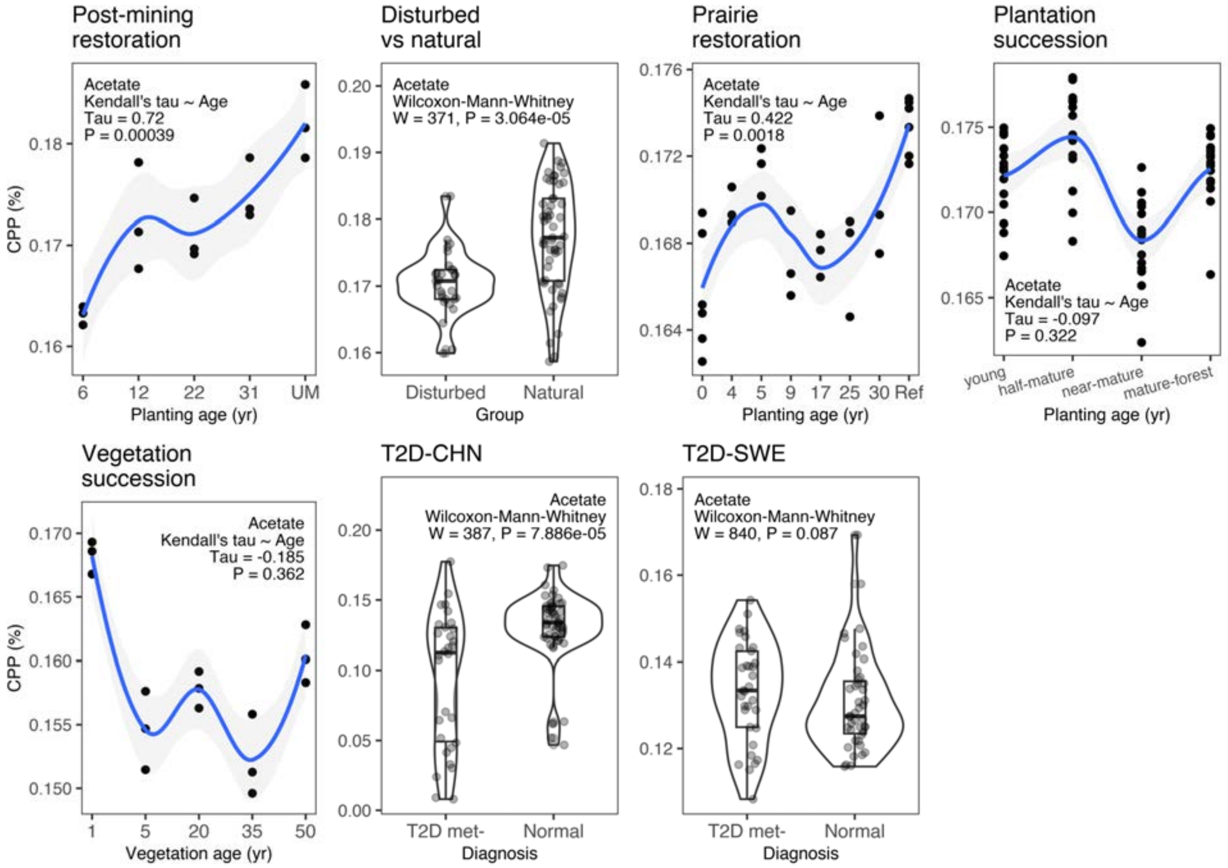


**Fig. S18.** Comparison of CPP values for **acetate** (dataset details in Extended Data Table 1).

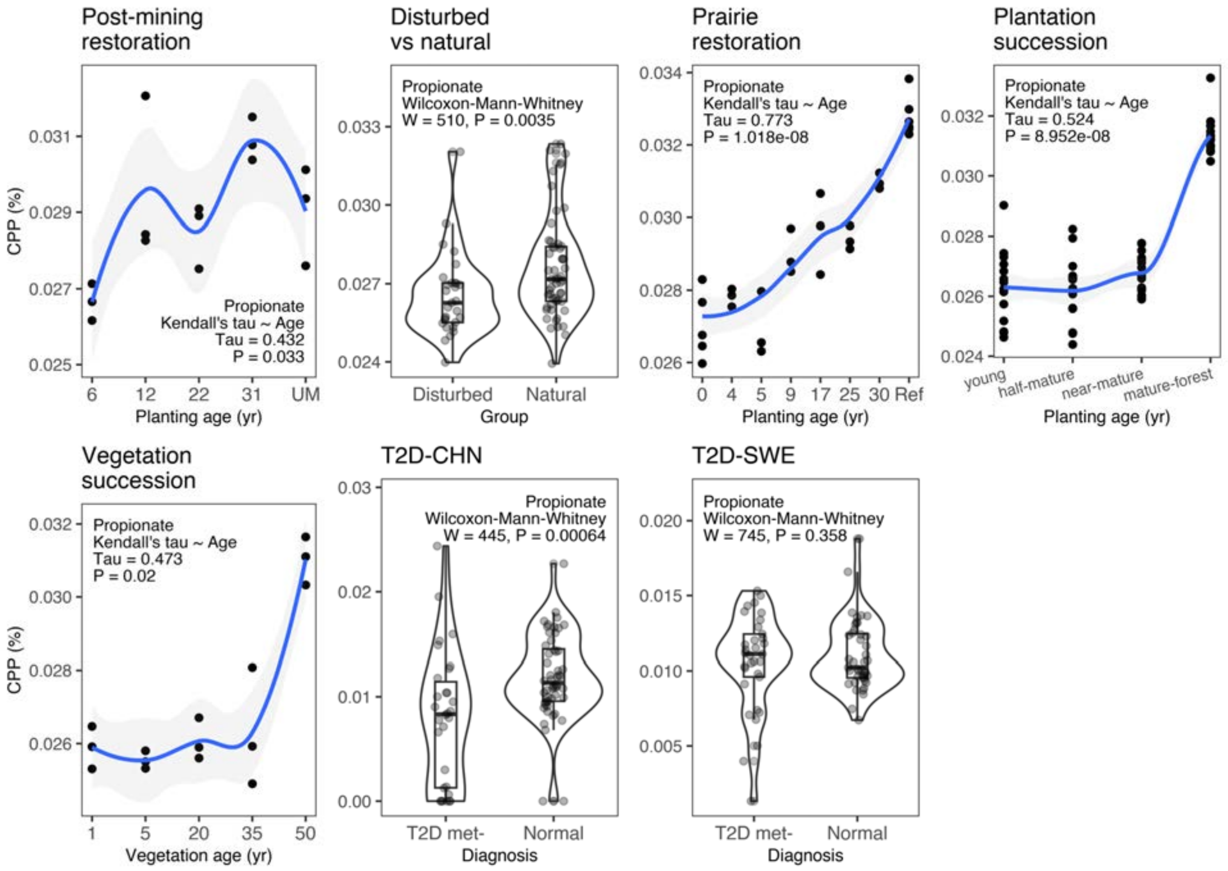


**Fig. S19.** Comparison of CPP values for **propionate** (dataset details in Extended Data Table 1).

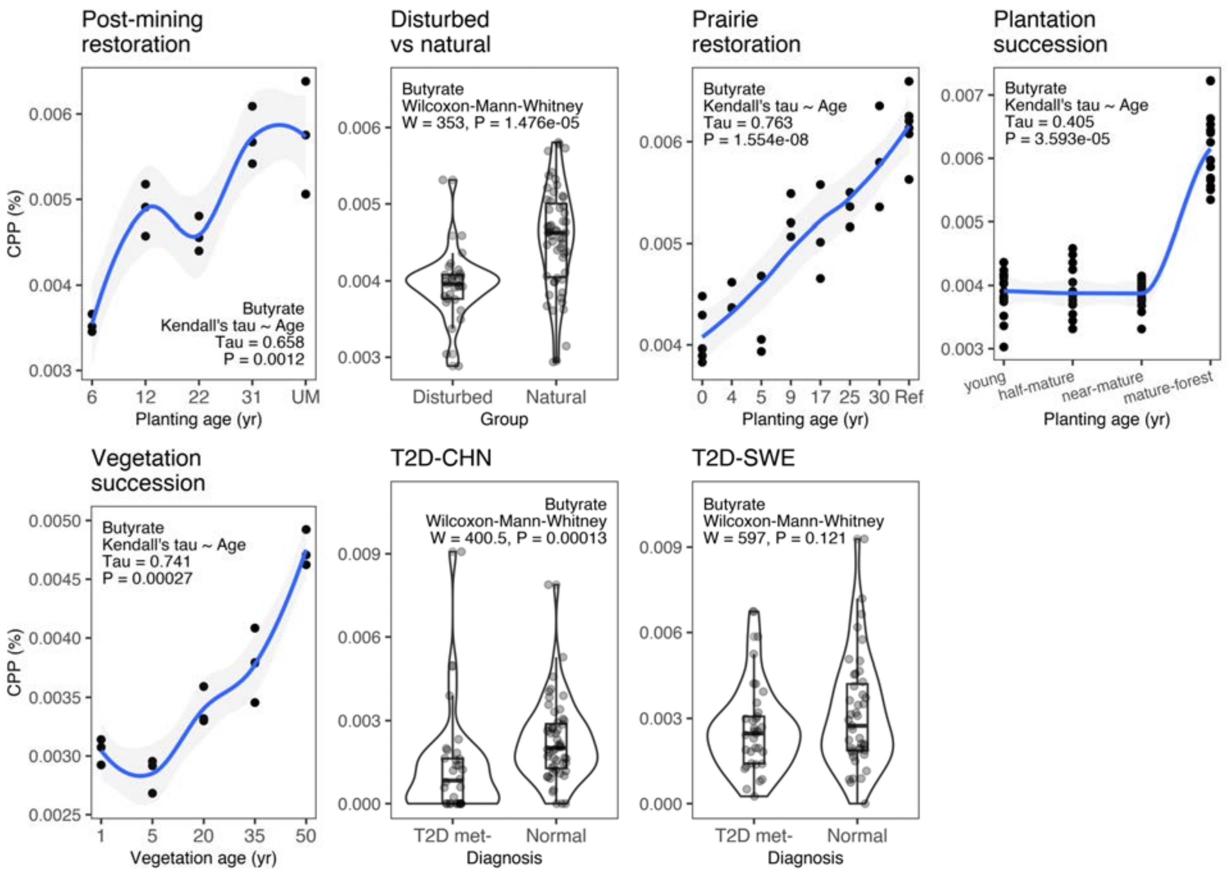


**Fig. S20.** Comparison of CPP values for **butyrate** (dataset details in Extended Data Table 1).

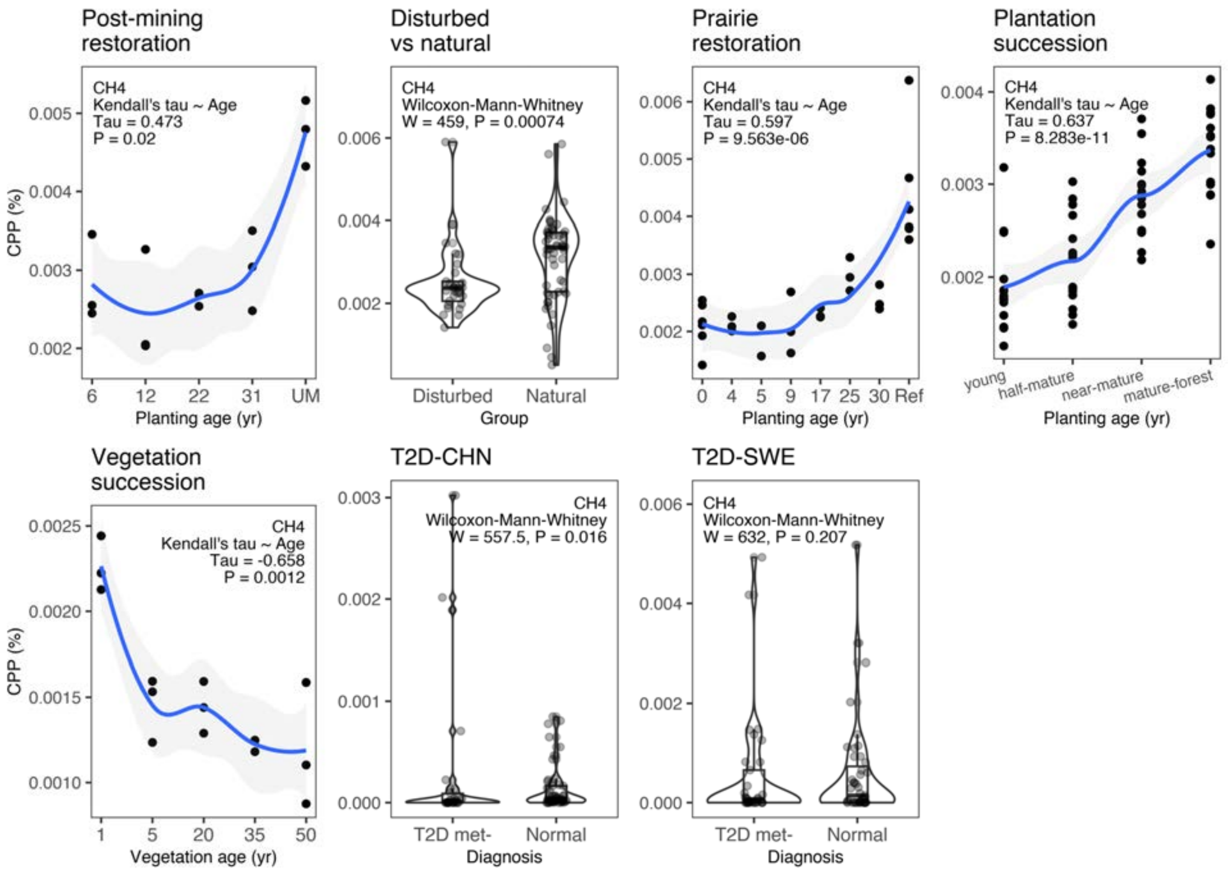


**Fig. S21.** Comparison of CPP values for **methane** ($CH_4$) (dataset details in Extended Data Table 1).

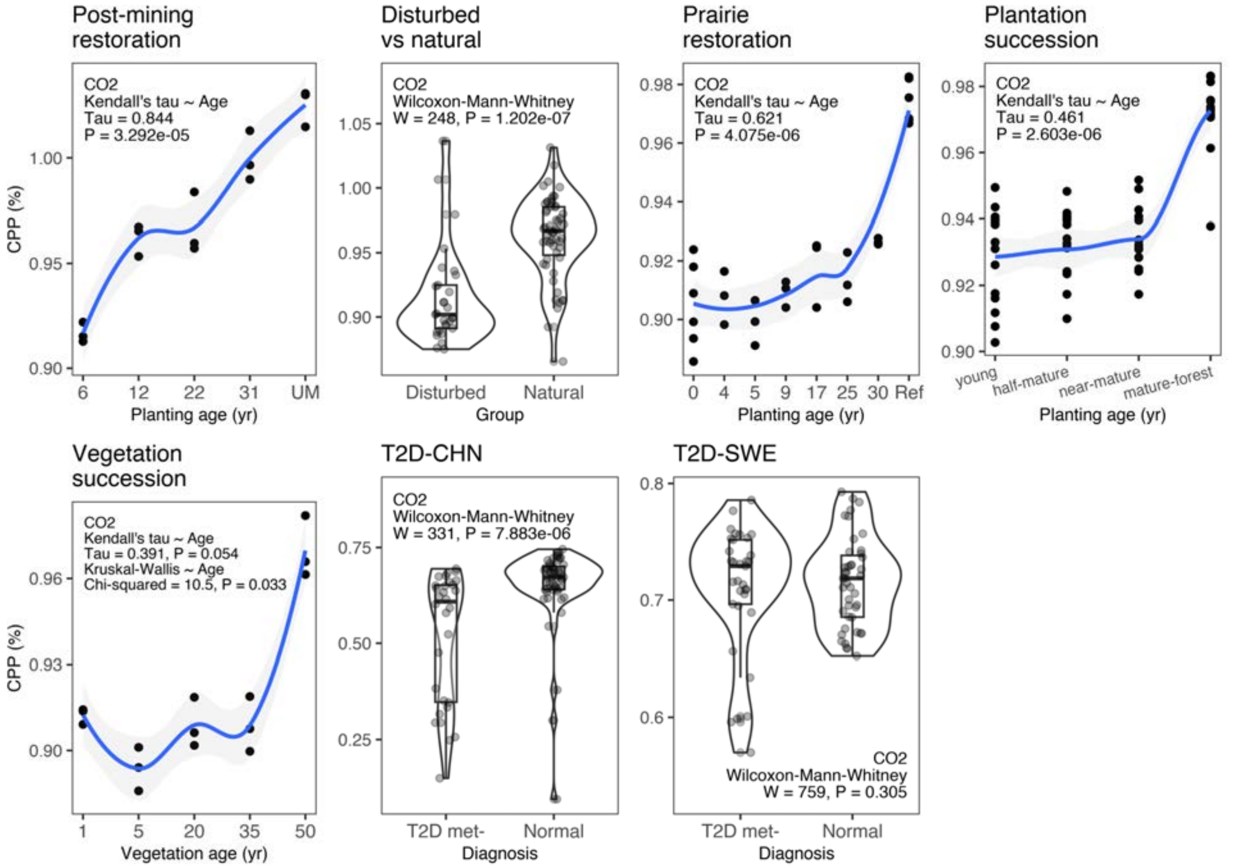


**Fig. S22.** Comparison of CPP values for **carbon dioxide** ($CO_2$) (dataset details in Extended Data Table 1).

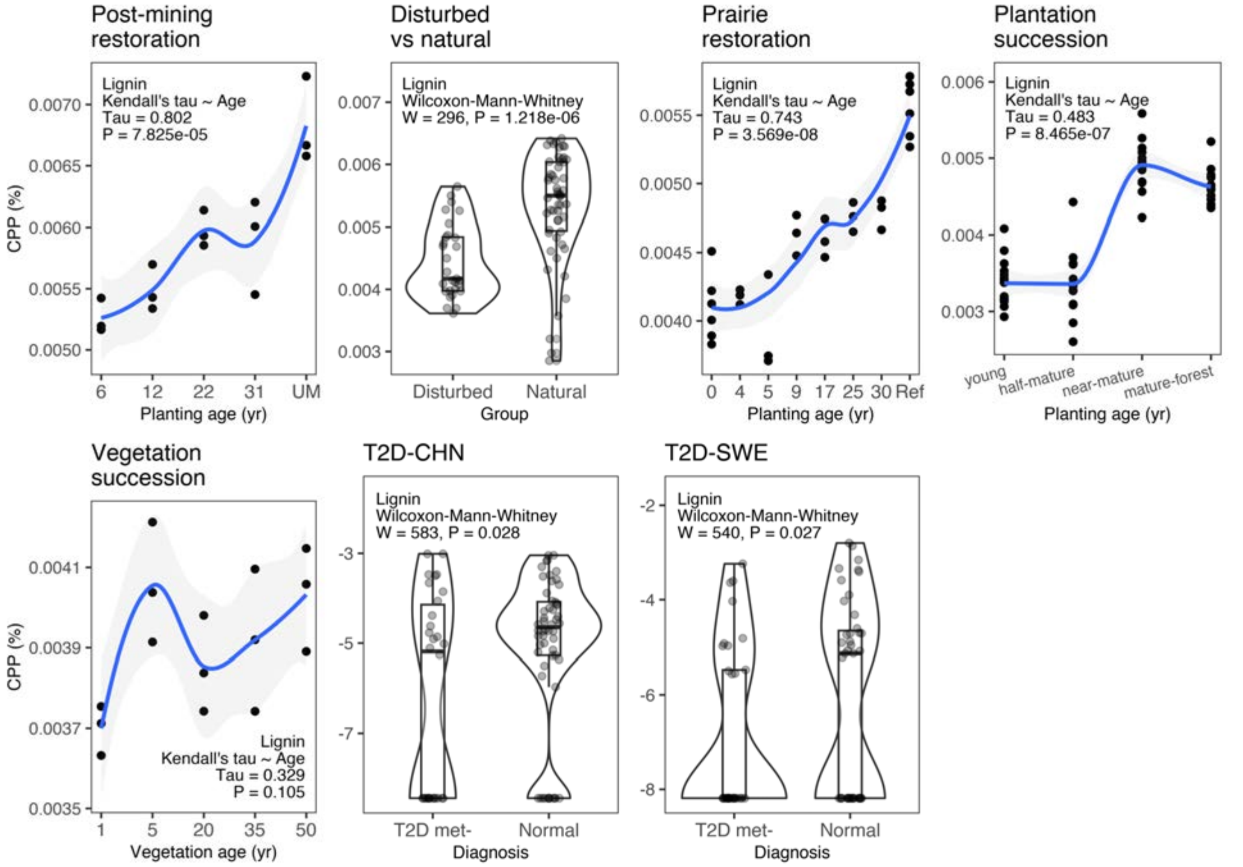


**Fig. S23.** Comparison of CPP values for **lignin** (dataset details in Extended Data Table 1).

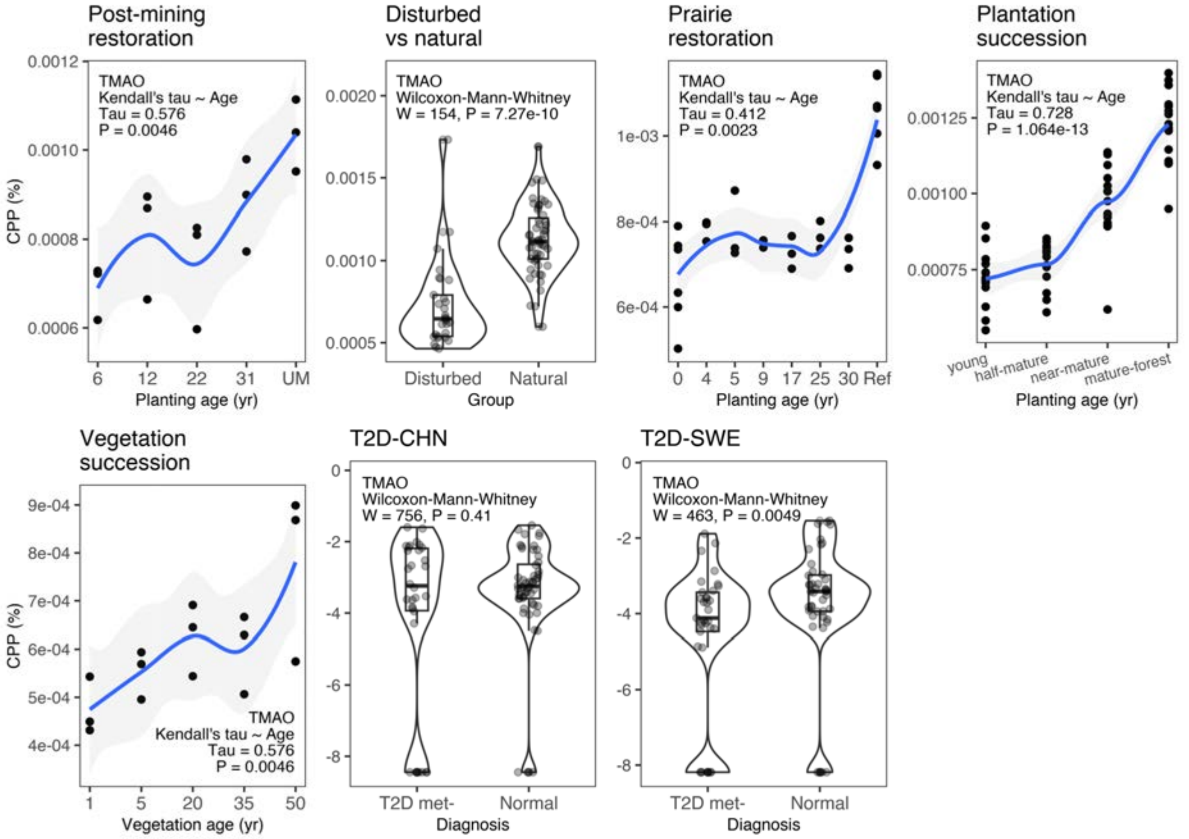


**Fig. S24.** Comparison of CPP values for **Trimethylamine N-oxide** (TMAO) (dataset details in Extended Data Table 1).

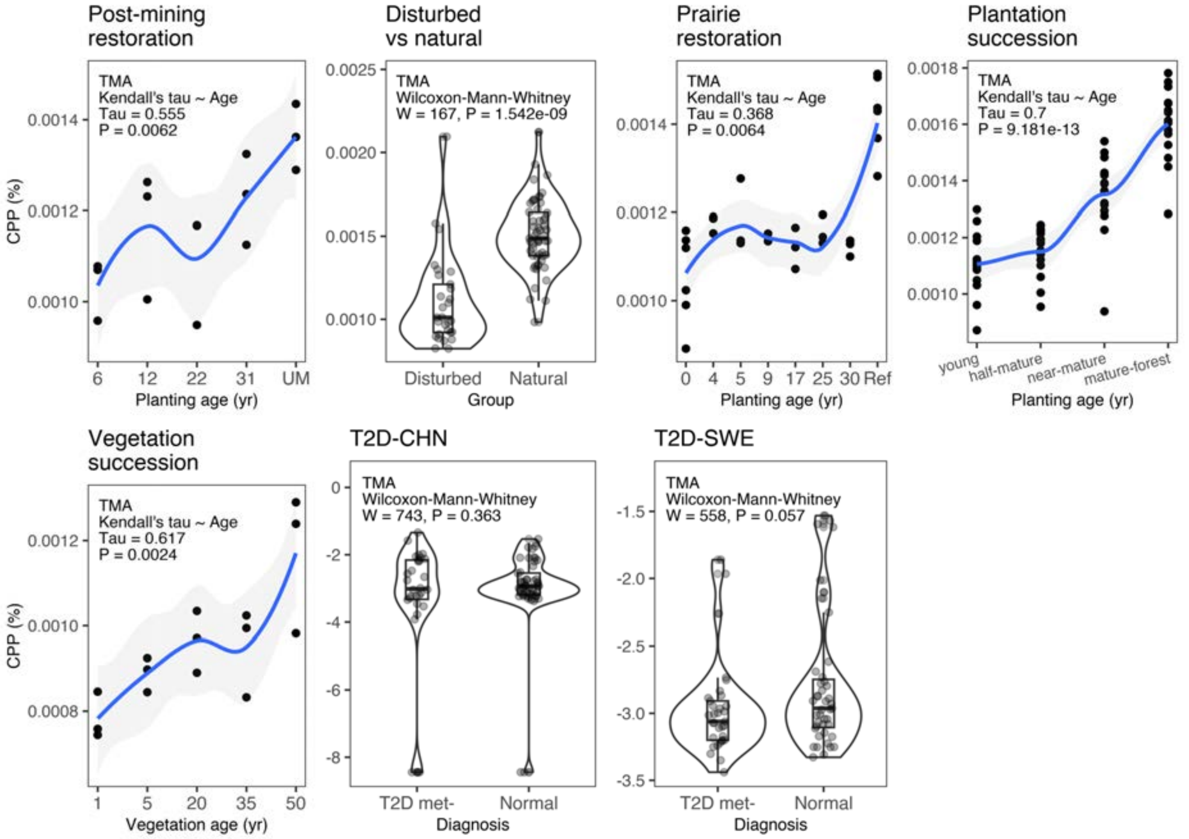


**Fig. S25.** Comparison of CPP values for **Trimethylamine** (TMA) (dataset details in Extended Data Table 1).

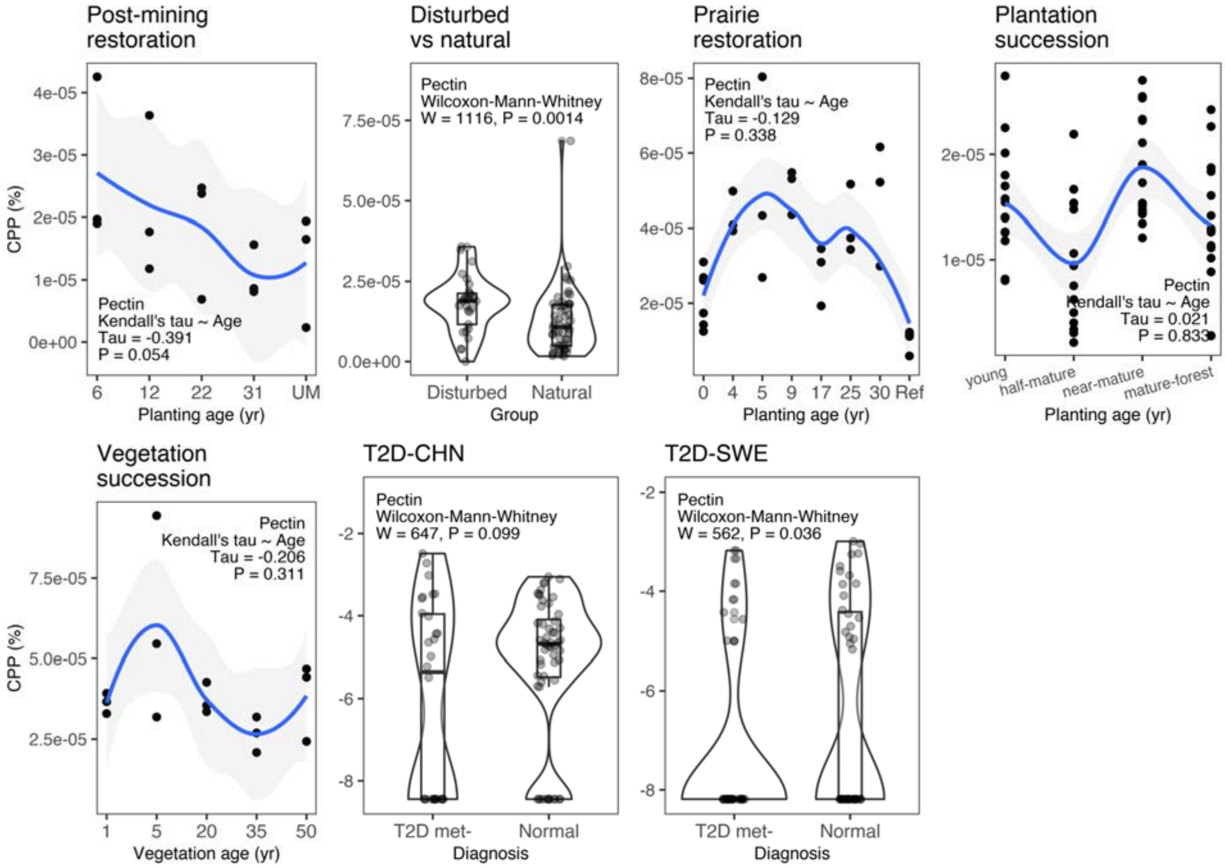


**Fig. S26.** Comparison of CPP values for **Pectin** (dataset details in Extended Data Table 1).

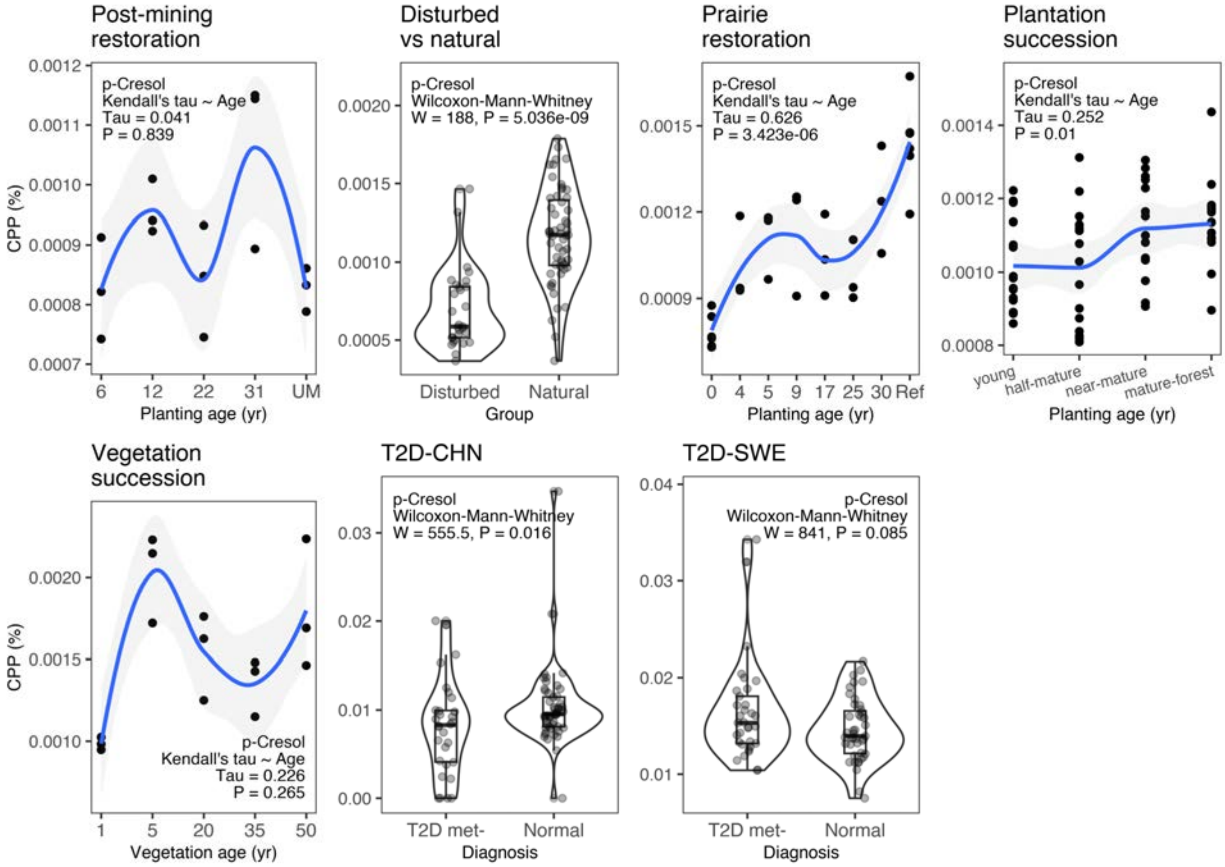


**Fig. S27.** Comparison of CPP values for **p-Cresol** (dataset details in Extended Data Table 1).

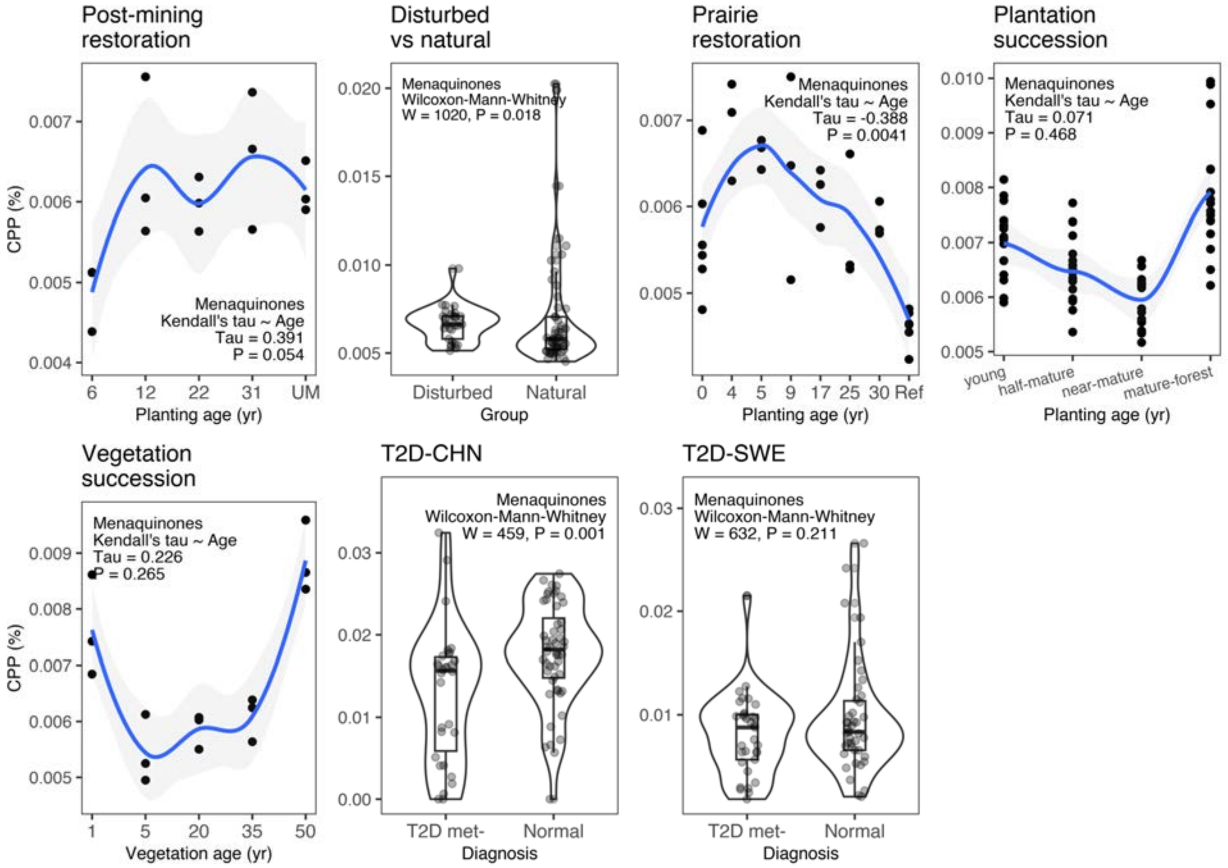


**Fig. S28.** Comparison of CPP values for **Menaquinones** (vitamin K2) (dataset details in Extended Data Table 1).

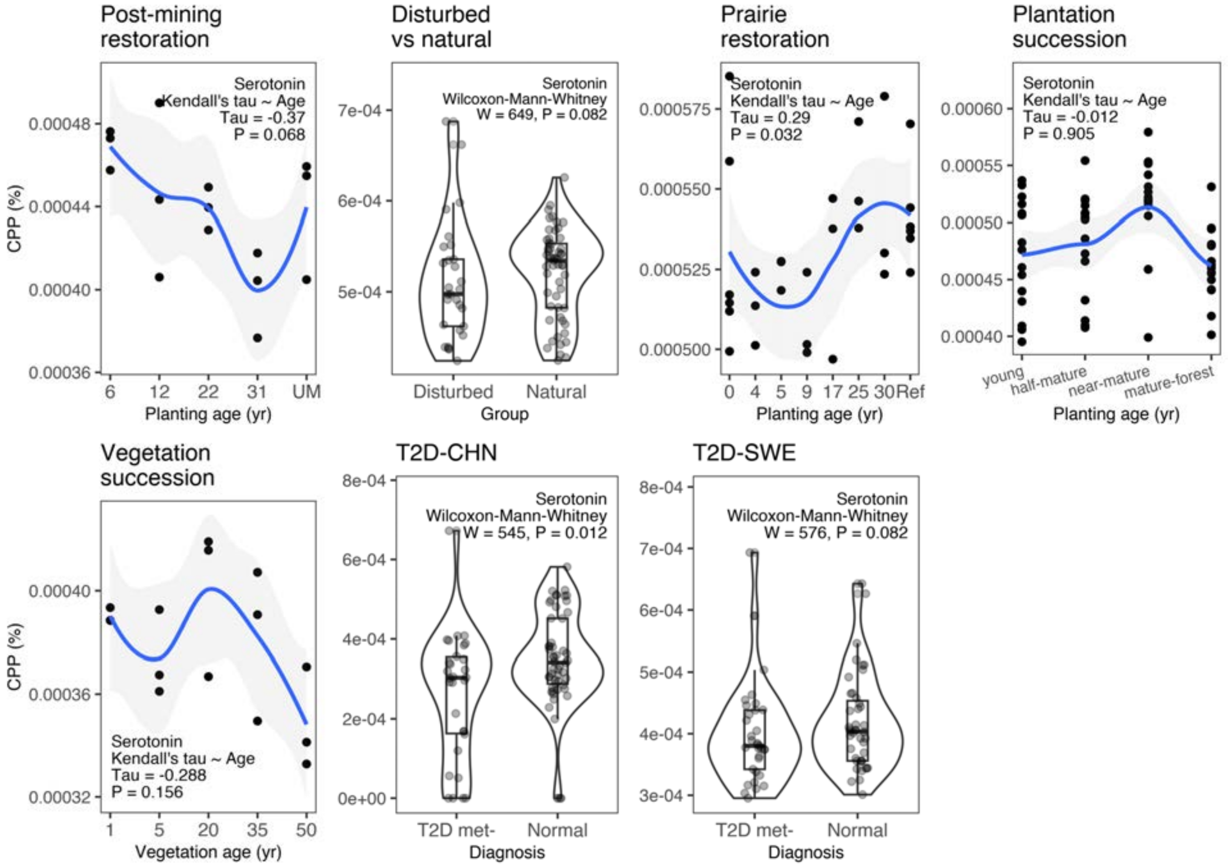


**Fig. S29.** Comparison of CPP values for **serotonin** (dataset details in Extended Data Table 1).

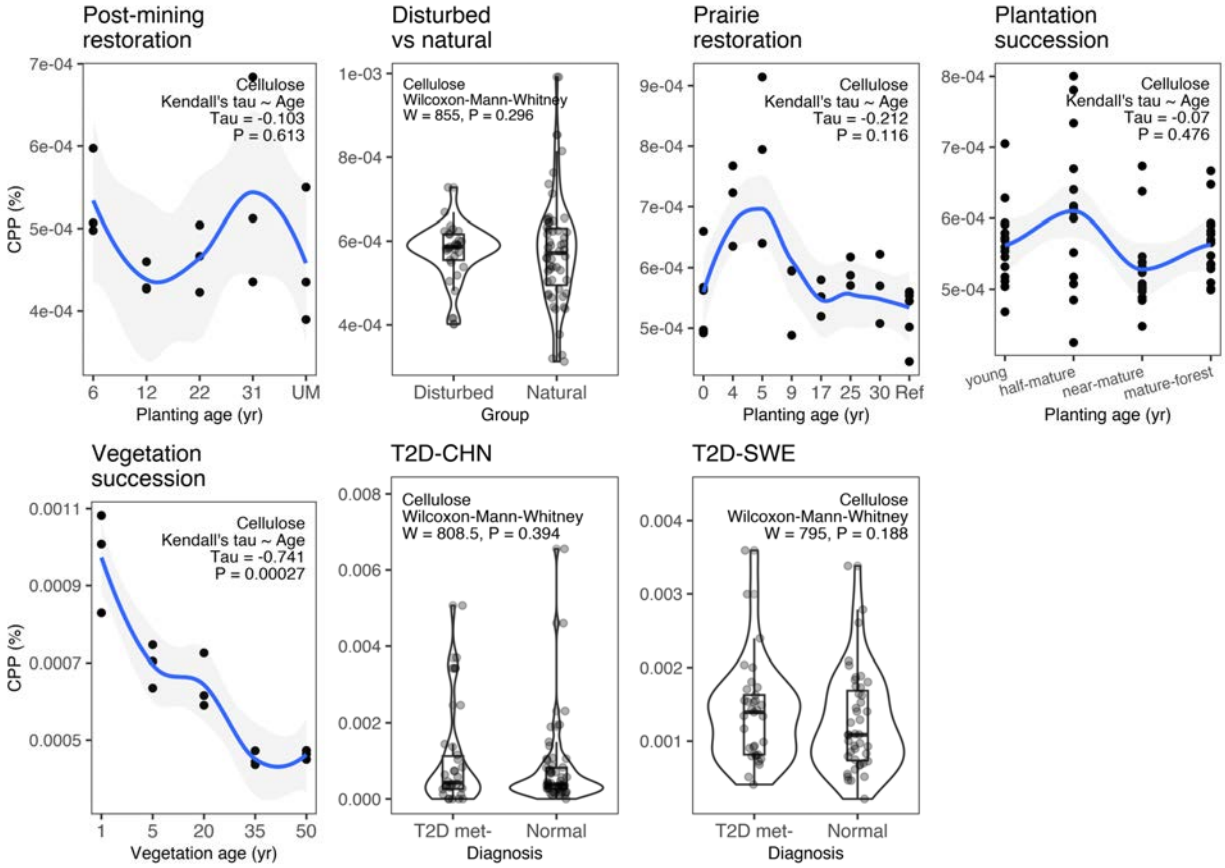


**Fig. S30.** Comparison of CPP values for **cellulose** (dataset details in Extended Data Table 1).

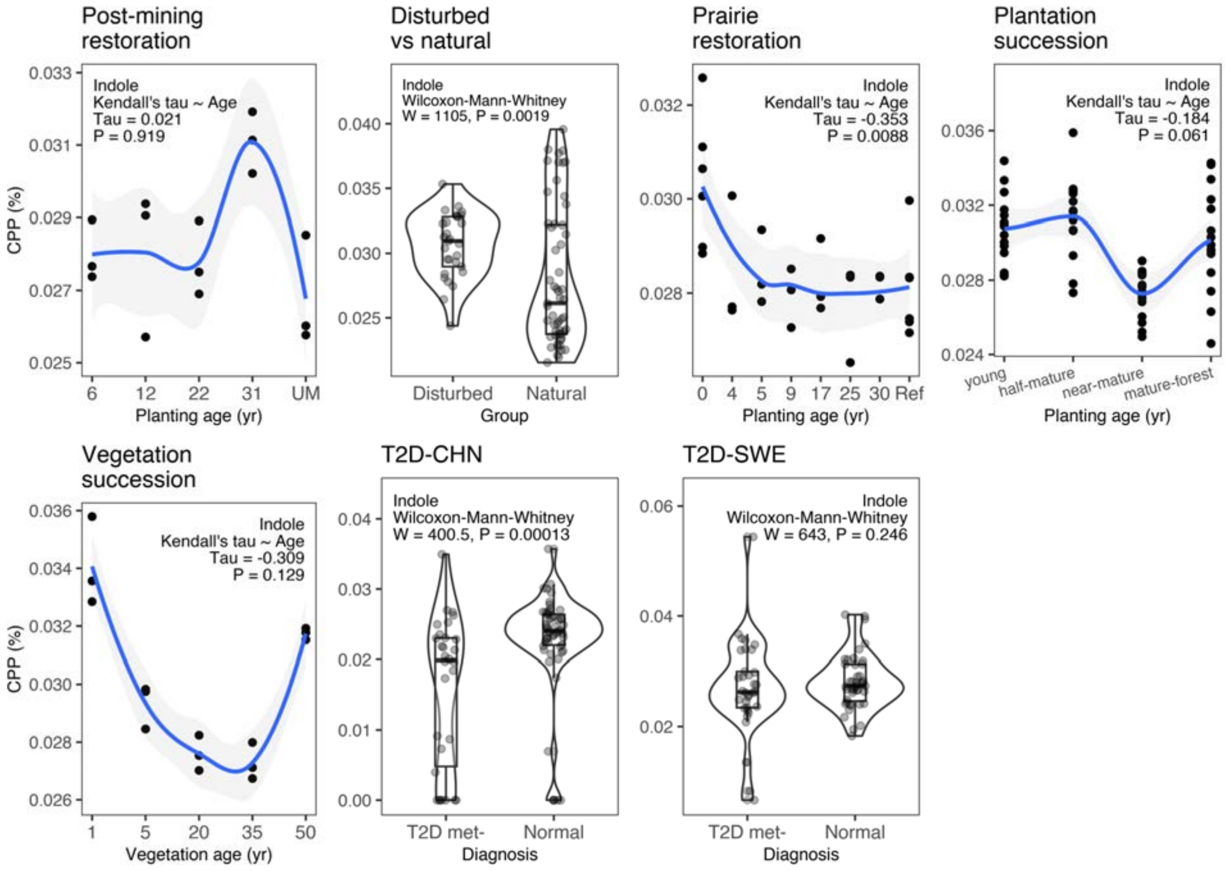


**Fig. S31.** Comparison of CPP values for **indole** (dataset details in Extended Data Table 1).

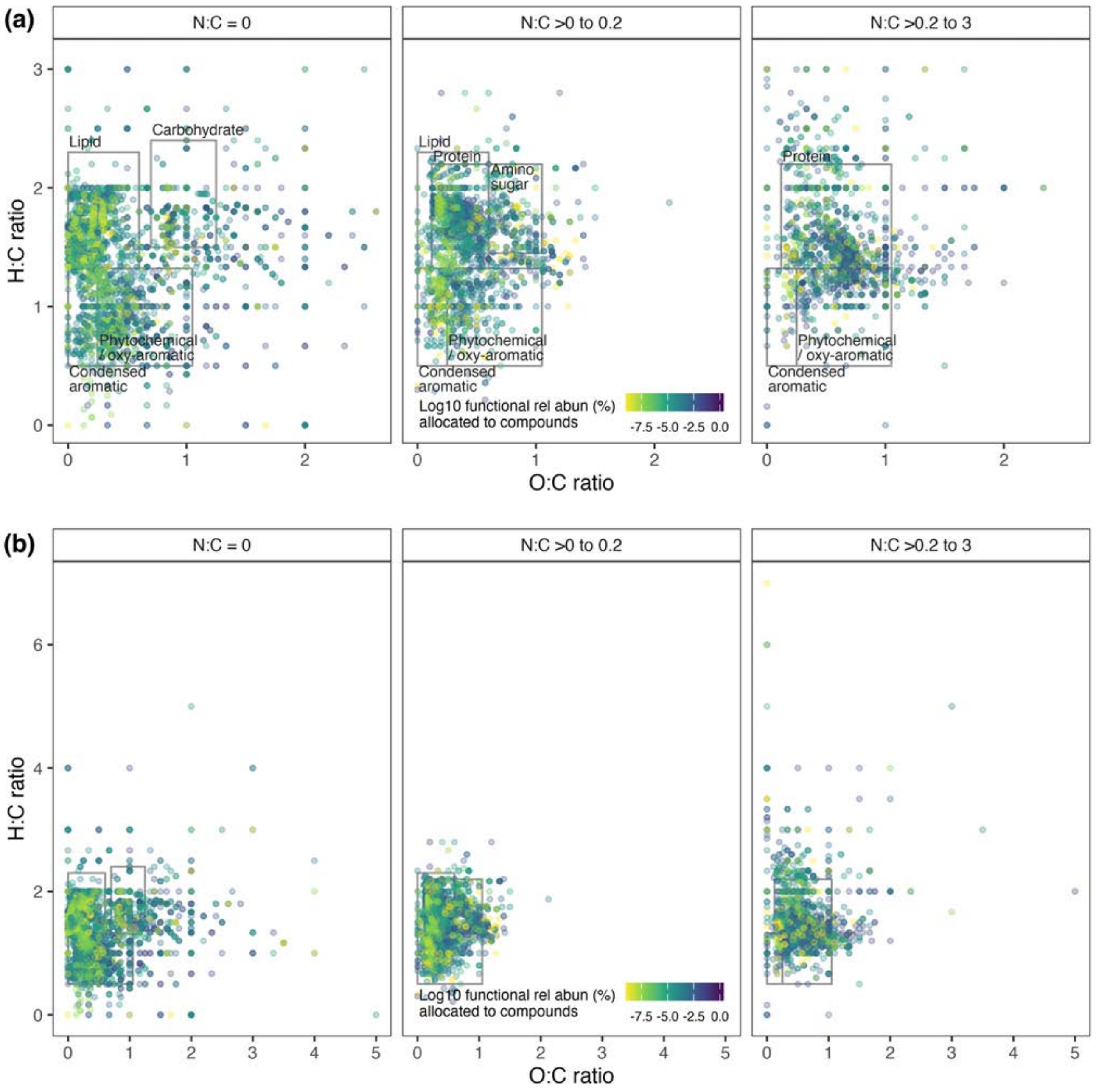


**Fig. S32.** Example visualization of carbon-containing compounds for a single post-mining restoration sample (from unmined site "UMA"). From a total of 8370 compounds observed across the entire restoration chronosequence dataset (total n = 15 samples), there are 7826 compounds with non-zero relative abundance in this particular sample. From this number, there are 7736 carbon-containing compounds that can be mapped using their chemical formula elemental O:C, H:C and N:C ratios. N:C ratios are calculated as continuous numeric values but displayed as three depth-slices of varying thickness to represent the third dimension. In panel (a) the O:C versus H:C panels are zoomed in from their full extent (b) to improve visualization of data-rich regions.

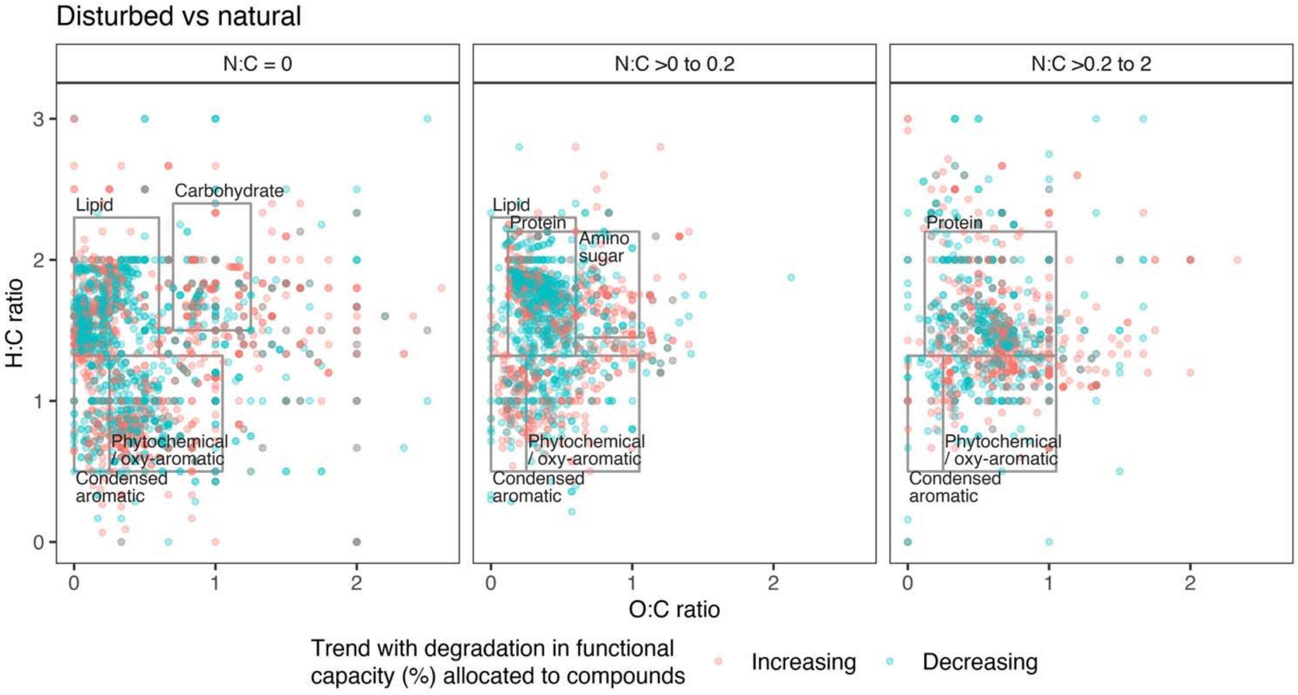


**Fig. S33.** Compound-wide trend analysis in **disturbed vs natural** soils (with *P*-adjustment for multiple testing) found *n* = 5,582 compounds (*n* = 5,181 with carbon) with CPP trends. With degradation, *n* = 2,618 compounds (*n* = 2,380 with carbon) had increased CPP values, while *n* = 2,964 compounds (*n* = 2,801 with carbon) had decreased CPP values.

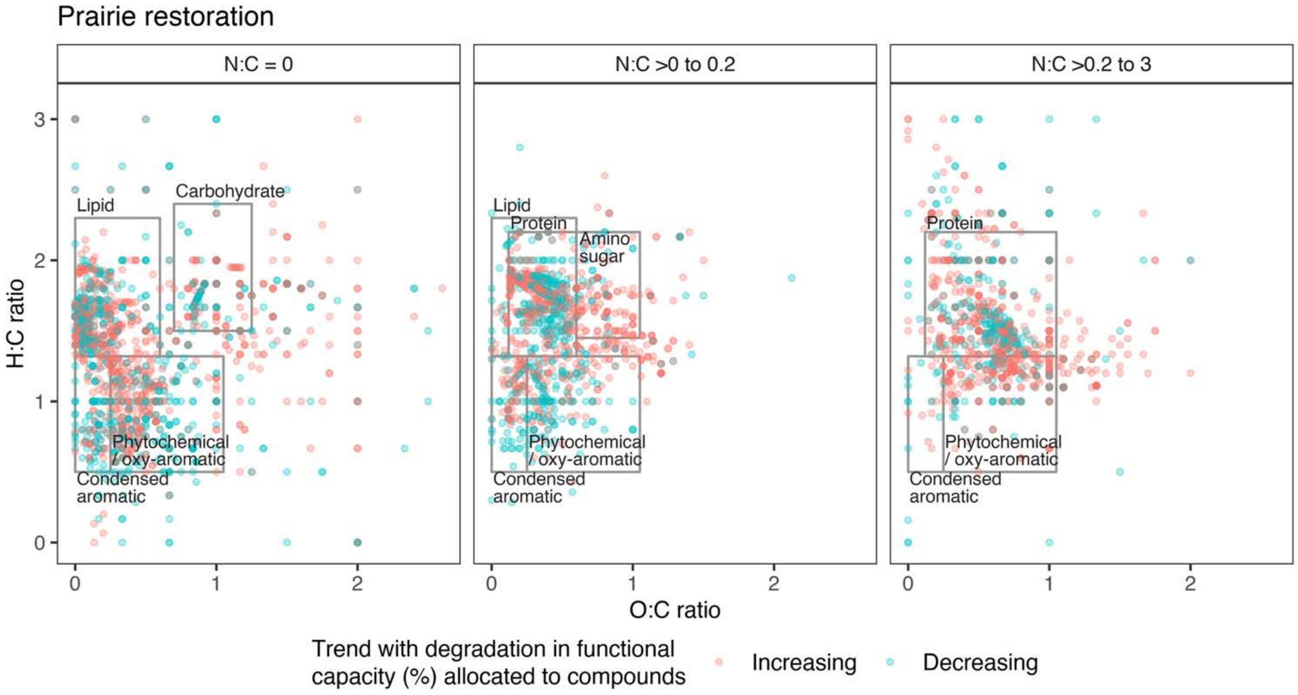


**Fig. S34.** Compound-wide trend analysis in **prairie restoration** soils (with *P*-adjustment for multiple testing) found *n* = 4,177 compounds (*n* = 3,880 with carbon) with CPP trends. With degradation, *n* = 2,422 compounds (*n* = 2,215 with carbon) had increased CPP values, while *n* = 1,755 compounds (*n* = 1,665 with carbon) had decreased CPP values.

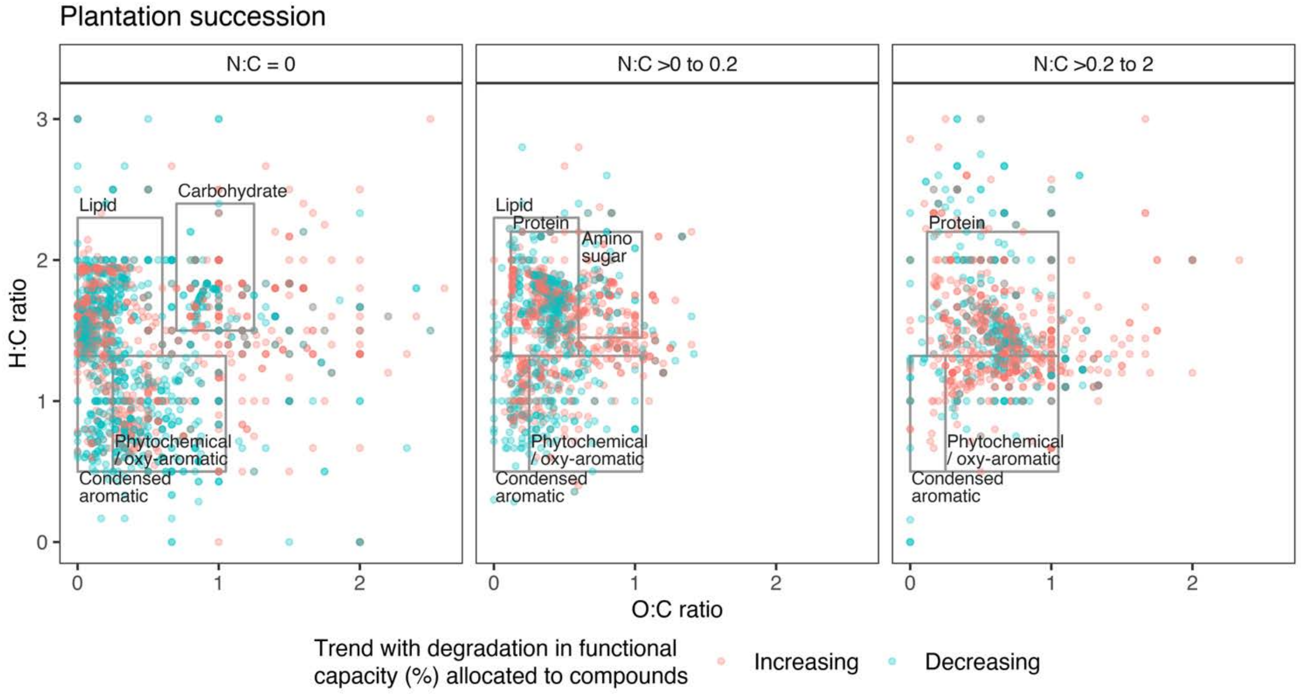


**Fig. 35.** Compound-wide trend analysis in **plantation succession** soils (with *P*-adjustment for multiple testing) found *n* = 4,779 compounds (*n* = 4,403 with carbon) with CPP trends. With degradation, *n* = 2,649 compounds (*n* = 2,402 with carbon) had increased CPP values, while *n* = 2,130 compounds (*n* = 2,001 with carbon) had decreased CPP values.

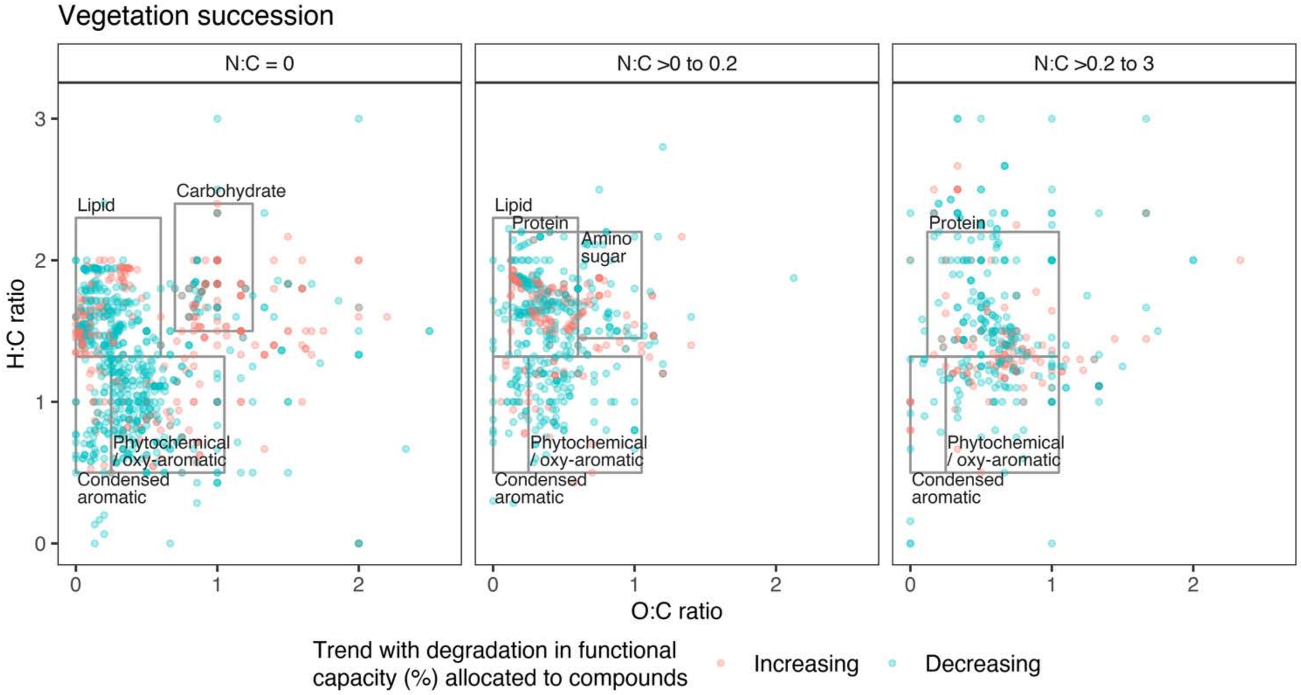


**Fig. 36.** Compound-wide trend analysis in **vegetation succession** soils (with *P*-adjustment for multiple testing) found *n* = 2,120 compounds (*n* = 1,958 with carbon) with CPP trends. With degradation, *n* = 716 compounds (*n* = 631 with carbon) had increased CPP values, while *n* = 1,404 compounds (*n* = 1,327 with carbon) had decreased CPP values.

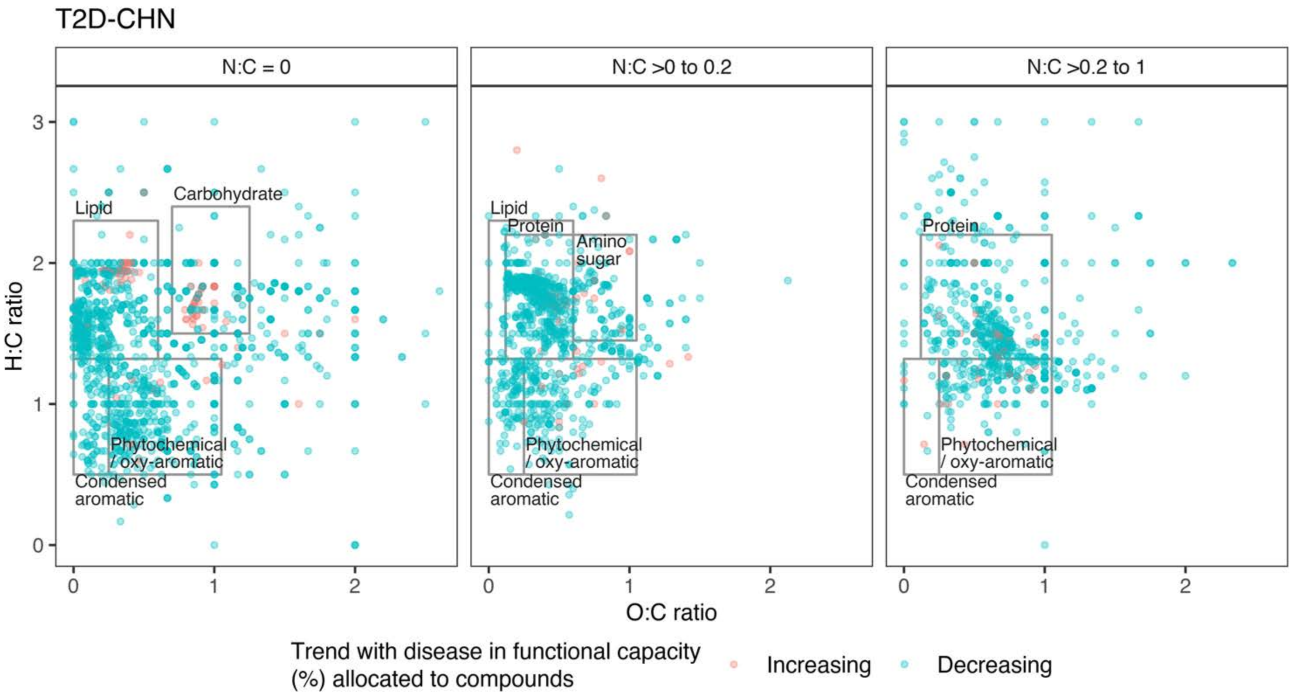


**Fig. 37.** Compound-wide trend analysis in the **Chinese (T2D-CHN)** T2D versus normal healthy cohort gut metagenomes (with *P*-adjustment for multiple testing) found *n* = 3,741 compounds (*n* = 3,451 with carbon) with CPP trends. With T2D, *n* = 255 compounds (*n* = 216 with carbon) had increased CPP values, while *n* = 3,486 compounds (*n* = 3,235 with carbon) had decreased CPP values

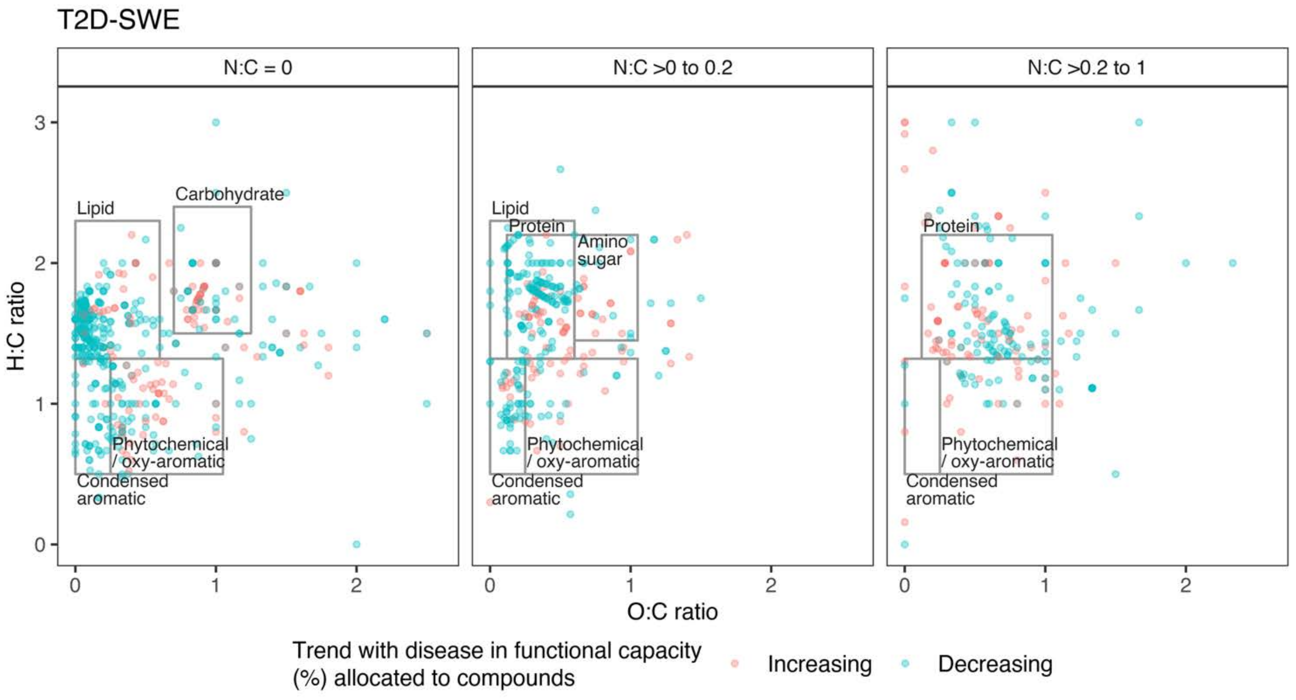


**Fig. 38.** Compound-wide trend analysis in the **Swedish (T2D-SWE)** T2D versus normal healthy cohort gut metagenomes (using unadjusted *P*-values) found *n* = 1,269 compounds (*n* = 1,181 with carbon) with CPP trends. With T2D, *n* = 358 compounds (*n* = 321 with carbon) had increased CPP values, while *n* = 911 compounds (*n* = 860 with carbon) had decreased CPP values.

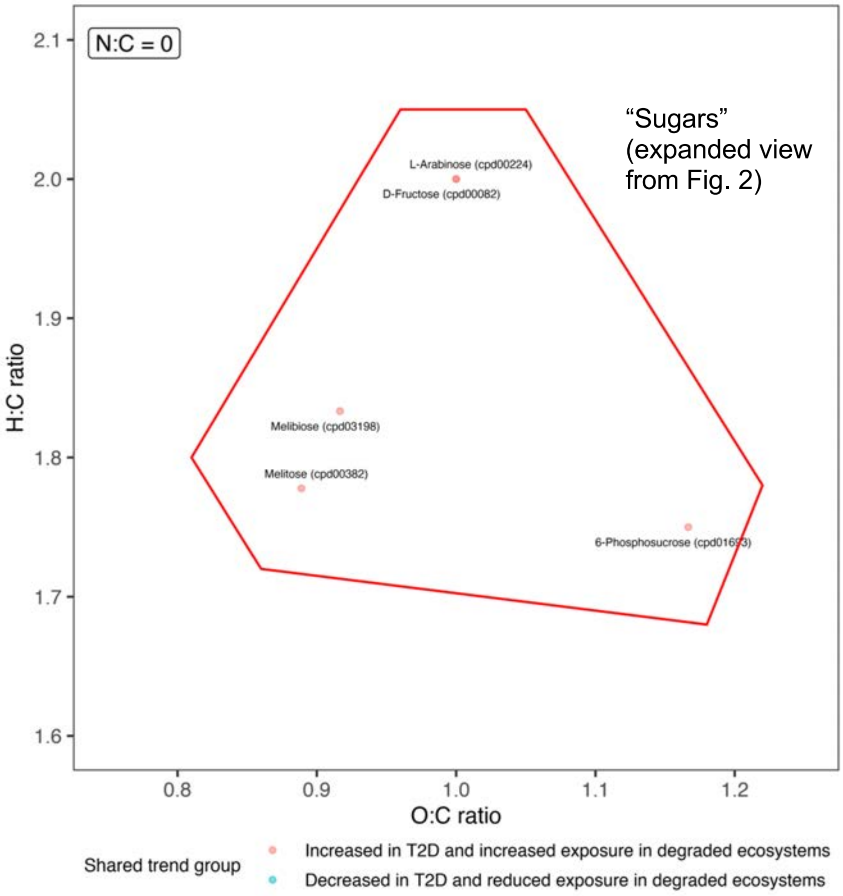


**Fig. S39.** The highlighted group (red points) comprise 5 sugars (L-arabinose, D-fructose, melibiose, melitose and 6-phosphosucrose) which all have increased potential metabolism with degradation and T2D based on shared directional CPP trends in 3 or more soil datasets, *P*-adjusted trends in T2D-CHN, and unadjusted trends in T2D-SWE datasets.

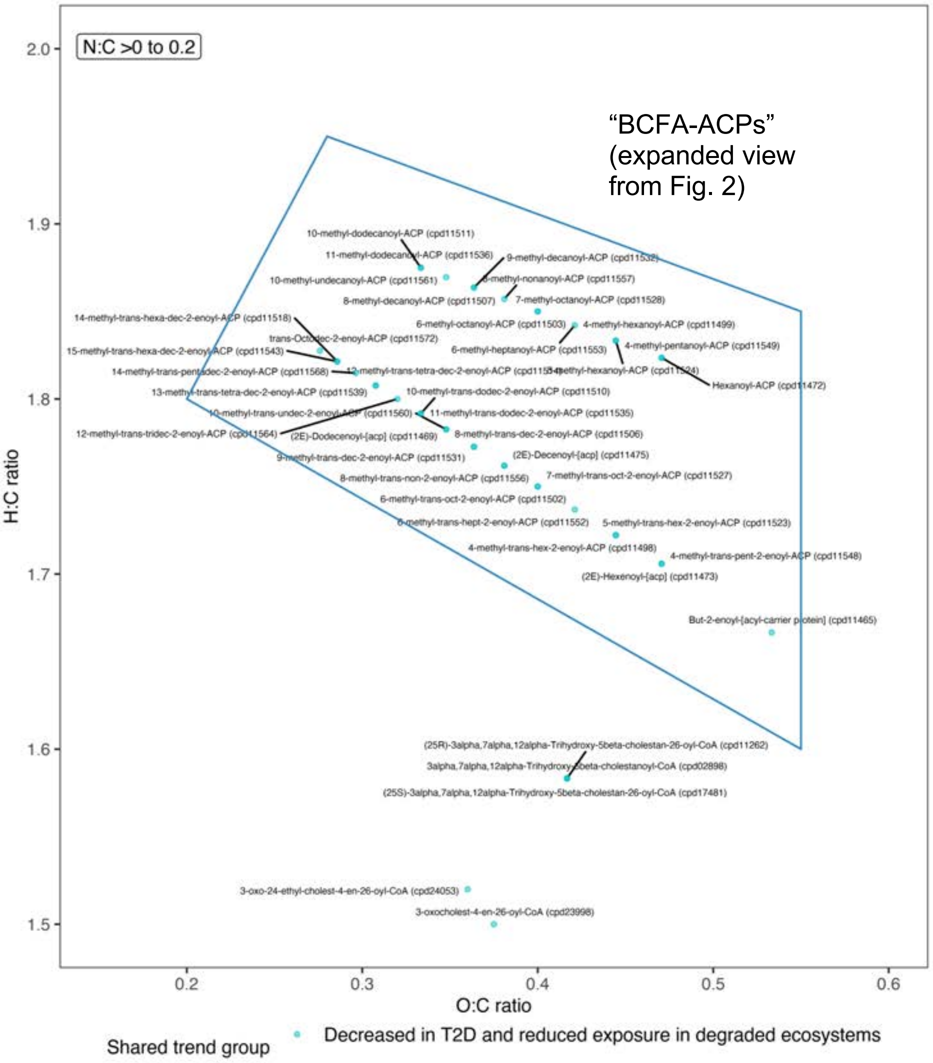


**Fig. S40.** The highlighted group (blue points) comprise 36 compounds dominated by branched-chain fatty acid – acyl-carrier proteins (BCFA-ACPs; Extended Data Table 4) involved in fatty acid biosynthesis, which all have decreased potential metabolism with degradation and T2D based on shared directional CPP trends in 3 or more soil datasets, *P*-adjusted trends in T2D-CHN, and unadjusted trends in T2D-SWE datasets.

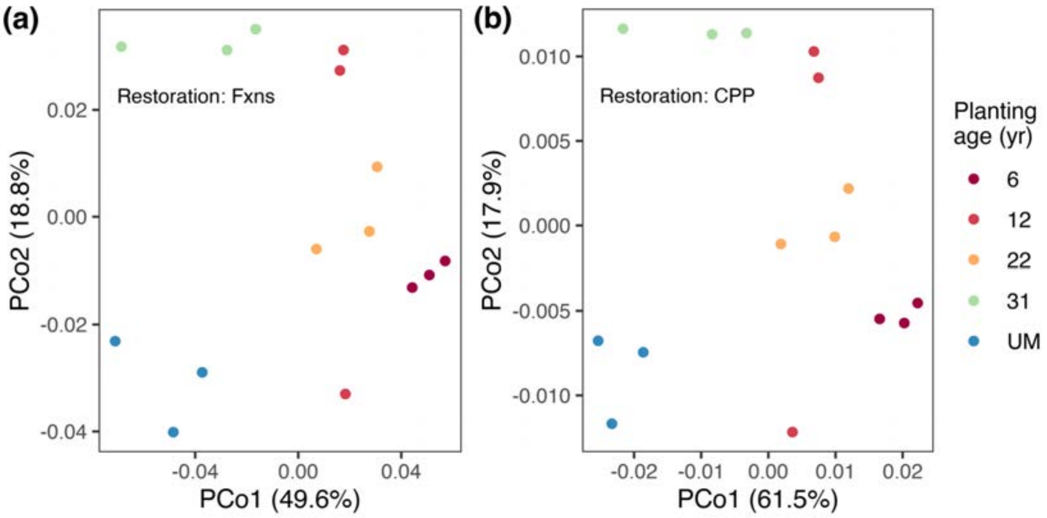


**Fig. S41.** PCoA ordinations of **post-mining restoration** soil metagenomes (raw data from[22]), using (a) SUPER-FOCUS functional potential (%) data (PERMANOVA|Soil-fxns: Df = 4, $R^2$= 0.67, F= 5.08, P= 0.002; Beta-disp|Soil-fxns: Df = 4, F = 0.345, P = 0.838; PCo1 + PCo2 axes explain 49.6 + 18.8 = 68.4 %); and (b) Compound processing potential (CPP, %) data (PERMANOVA|Soil-CPP: Df = 4, $R^2$= 0.759, F= 7.87, P= 0.002; Beta-disp|Soil-CPP: Df= 4, F= 0.364, P = 0.842; PCo1 + PCo2 axes explain 61.5 + 17.9 = 79.4 %).

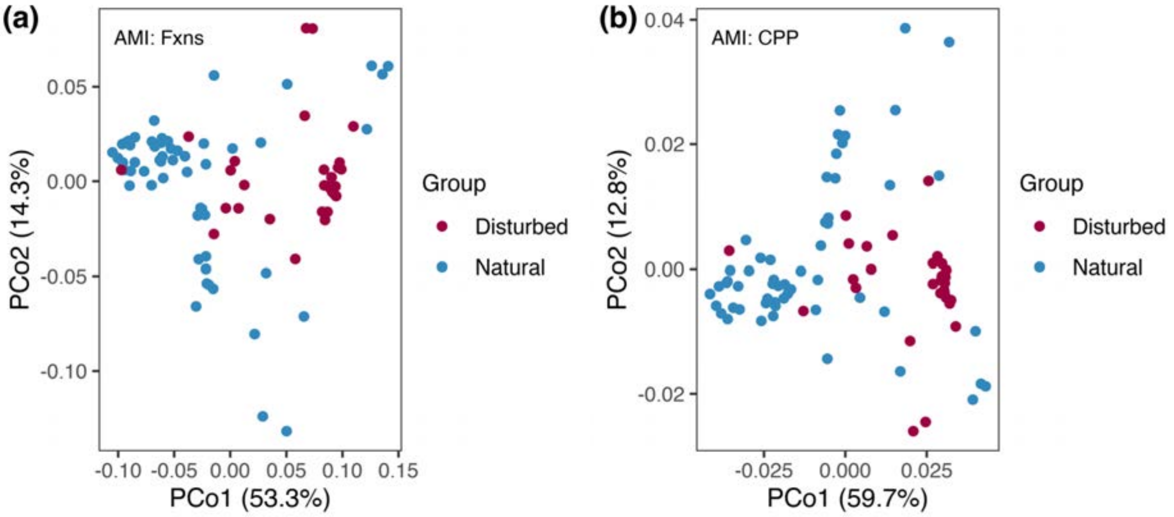


**Fig. S42.** PCoA ordinations of **disturbed vs natural soils** (raw data from the Australian Microbiome Initiative[23], described in[24]), using (a) SUPER-FOCUS functional potential (%) data (PERMANOVA|Soil-fxns: Df = 1, $R^2$= 0.193, F= 19.646, P= 0.001; Beta-disp|Soil-fxns: Df = 1, F = 0.553, P = 0.466; PCo1 + PCo2 axes explain 53.3 + 14.3 = 67.6 %); and (b) Compound processing potential (CPP, %) data (PERMANOVA|Soil-CPP: Df = 1, $R^2$= 0.202, F= 20.785, P= 0.001; Beta-disp|Soil-CPP: Df= 1, F= 1.843, P = 0.184; PCo1 + PCo2 axes explain 59.7 + 12.8 = 72.5 %).

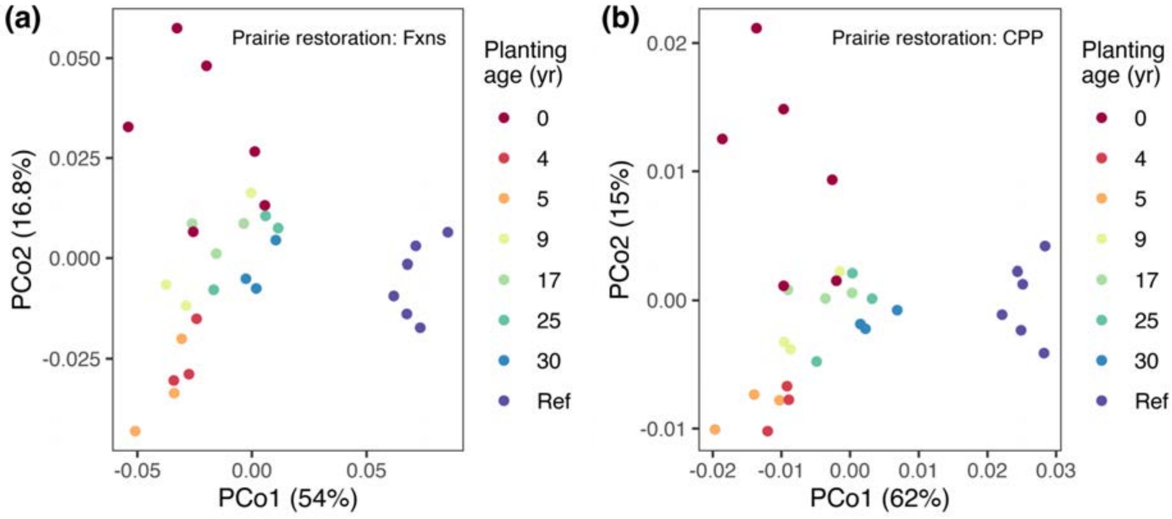


**Fig. S43.** PCoA ordinations of **prairie restoration** soils (raw data from[25]), using (a) SUPER-FOCUS functional potential (%) data (PERMANOVA|Soil-fxns: Df = 7, $R^2$= 0.711, F= 7.716, P= 0.001; Beta-disp|Soil-fxns: Df = 7, F = 1.62, P = 0.207; PCo1 + PCo2 axes explain 54 + 16.8 = 70.8 %); and (b) Compound processing potential (CPP, %) data (PERMANOVA|Soil-CPP: Df = 7, $R^2$= 0.762, F= 10.06, P= 0.001; Beta-disp|Soil-CPP: Df= 7, F= 1.8314, P = 0.145; PCo1 + PCo2 axes explain 62 + 15 = 77 %).

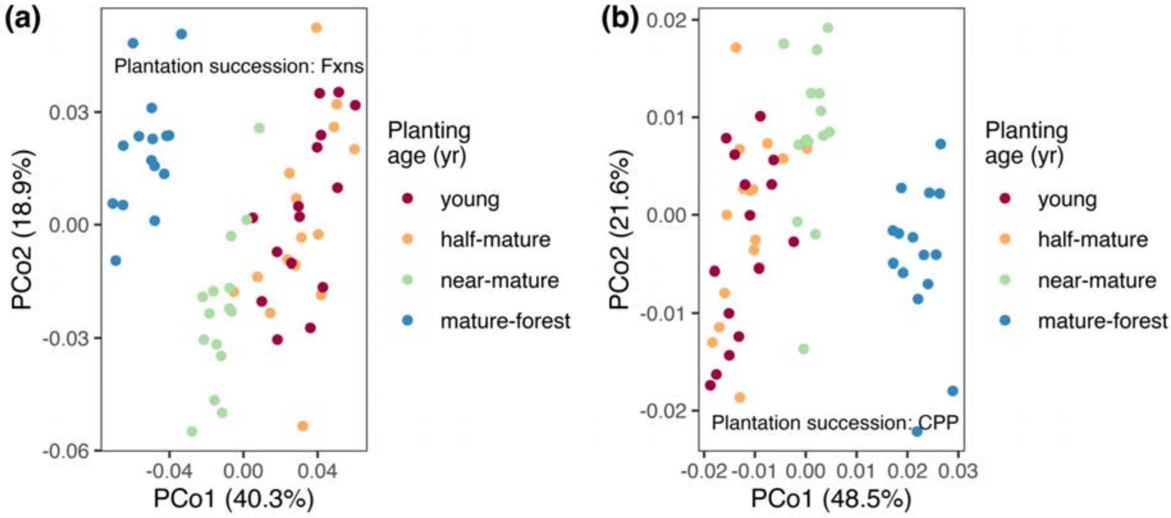


**Fig. S44.** PCoA ordinations of **plantation succession** soils (raw data from[26]), using (a) SUPER-FOCUS functional potential (%) data (PERMANOVA|Soil-fxns: Df = 3, $R^2$= 0.488, F= 17.785, P= 0.001; Beta-disp|Soil-fxns: Df = 3, F = 3.512, P = 0.023; PCo1 + PCo2 axes explain 40.3 + 18.9 = 59.2 %); and (b) Compound processing potential (CPP, %) data (PERMANOVA|Soil-CPP: Df = 3, $R^2$= 0.564, F= 24.10, P= 0.001; Beta-disp|Soil-CPP: Df= 3, F= 2.13, P = 0.09; PCo1 + PCo2 axes explain 48.5 + 21.6 = 70.1 %).

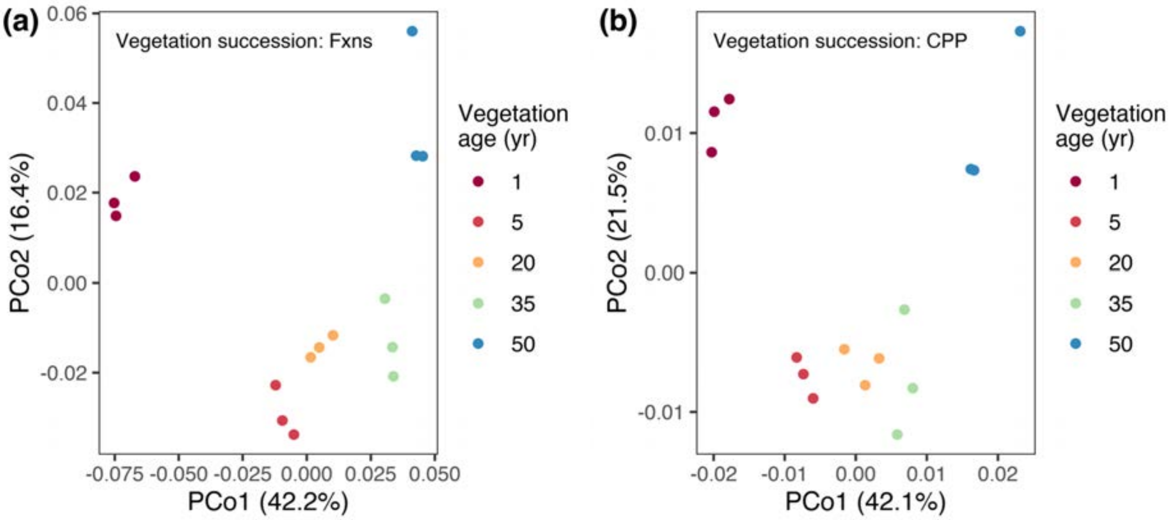


**Fig. S45.** PCoA ordinations of **vegetation succession** soils (raw data from[27]), using (a) SUPER-FOCUS functional potential (%) data (PERMANOVA|Soil-fxns: Df = 4, $R^2$= 0.758, F= 7.844, P= 0.001; Beta-disp|Soil-fxns: Df = 4, F = 0.1147, P = 0.964; PCo1 + PCo2 axes explain 42.2 + 16.4 = 58.6 %); and (b) Compound processing potential (CPP, %) data (PERMANOVA|Soil-CPP: Df = 4, $R^2$= 0.801, F= 10.065, P= 0.001; Beta-disp|Soil-CPP: Df= 4, F= 0.088, P = 0.981; PCo1 + PCo2 axes explain 42.1 + 21.5 = 63.6 %).

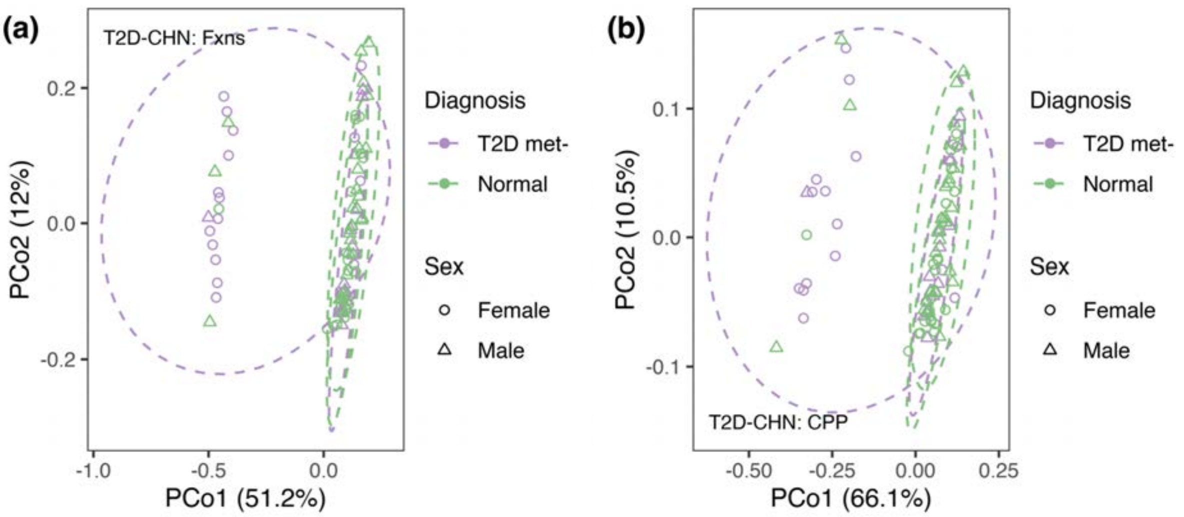


**Fig. S46.** PCoA ordinations of **T2D-CHN cohort** (raw data from[28, 29]), using (a) SUPER-FOCUS functional potential (%) data (PERMANOVA|Gut-fxns: ~Diagnosis: Df = 1, $R^2$= 0.095, F= 8.66, P= 0.001; Sex: Df = 1, $R^2$= 0.036, F= 3.31, P= 0.016 ; Beta-disp|Gut-CPP: ~Diagnosis: Df= 1, F= 17.537, P = 0.001; i.e., diagnosis groups have different beta-dispersions, which may invalidate Adonis-PERMANOVA result for different centroids; PCo1 + PCo2 axes explain 51.2 + 12= 63.2 %); and (b) Compound processing potential (CPP, %) data (PERMANOVA|Gut-CPP: ~Diagnosis: Df = 1, $R^2$= 0.1096, F= 10.216, P= 0.001; Sex: Df = 1, $R^2$= 0.043, F= 3.96, P= 0.016 ; Beta-disp|Gut-CPP: ~Diagnosis: Df= 1, F= 15.794, P = 0.001; i.e., diagnosis groups have different beta-dispersions, which may invalidate Adonis-PERMANOVA result for different centroids; PCo1 + PCo2 axes explain 66.1 + 10.5 = 76.6 %).

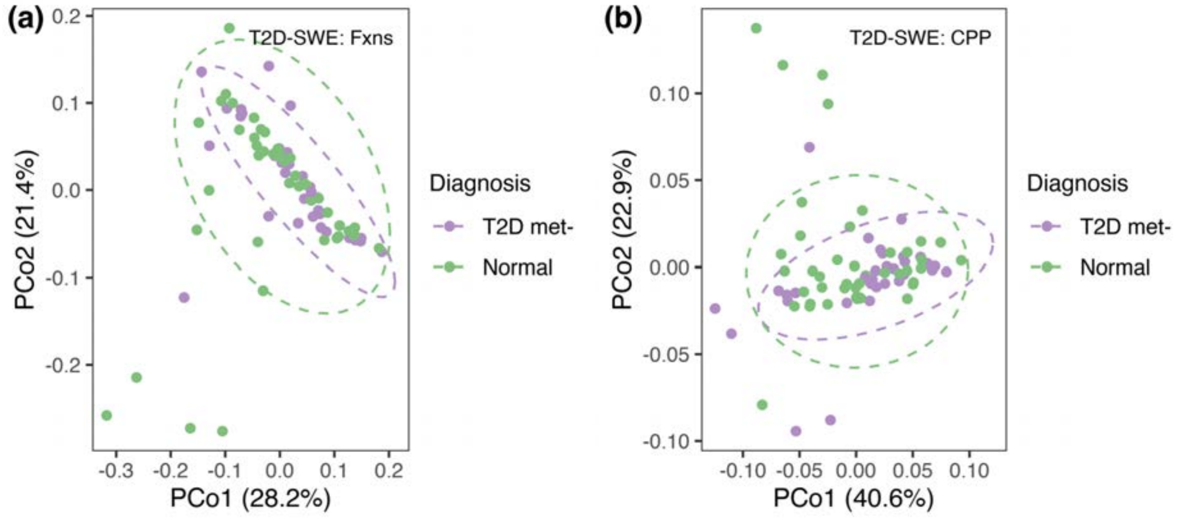


**Fig. S47.** PCoA ordinations of **T2D-SWE cohort** (raw data from[28, 30]), using (a) SUPER-FOCUS functional potential (%) data (PERMANOVA|Gut-fxns: Df = 1, $R^2$= 0.02, F= 1.549, P= 0.136; Beta-disp|Gut-fxns: Df = 1, F =0.158, P = 0.709; PCo1 + PCo2 axes explain 28.2 + 21.4= 49.6 %); and (b) Compound processing potential (CPP, %) data (PERMANOVA|Gut-CPP: Df = 1, $R^2$= 0.02, F= 1.478, P= 0.175; Beta-disp|Gut-CPP: Df= 1, F= 5e-04, P = 0.981; PCo1 + PCo2 axes explain 40.6 + 22.9 = 63.5 %).

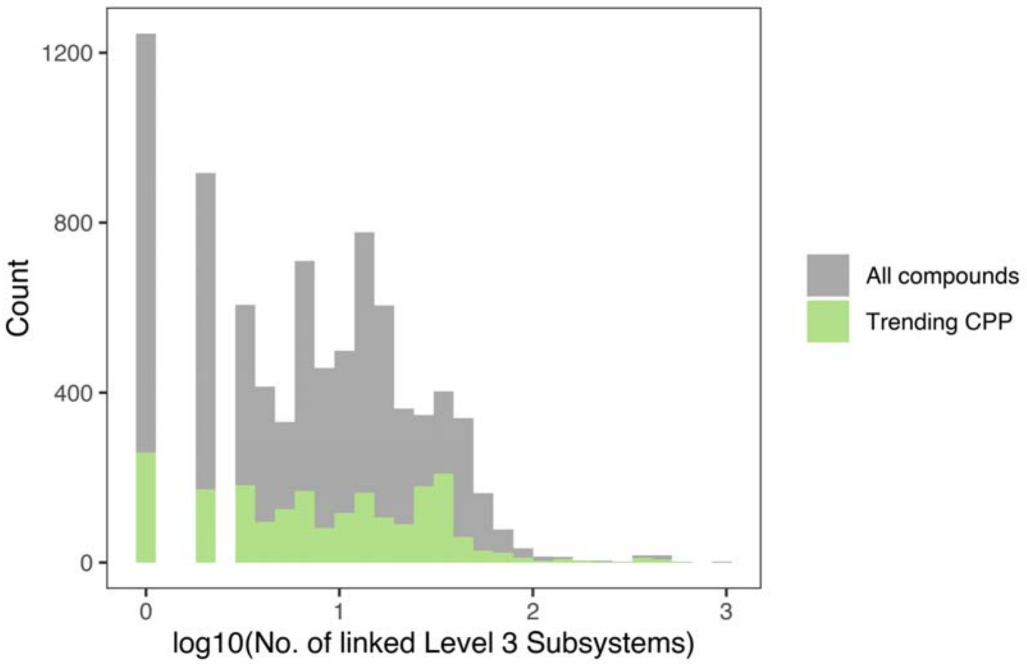


**Fig. S48.** Histogram distributions of numbers of linked level 3 subsystems for compounds with trending CPP ($n$ = 2,122), compared to all compounds ($n$ = 8,370), in the post-mining restoration dataset. The plot indicates that detection of significant trends draws widely from across the spectrum of 'more connected' to 'less connected' compounds.

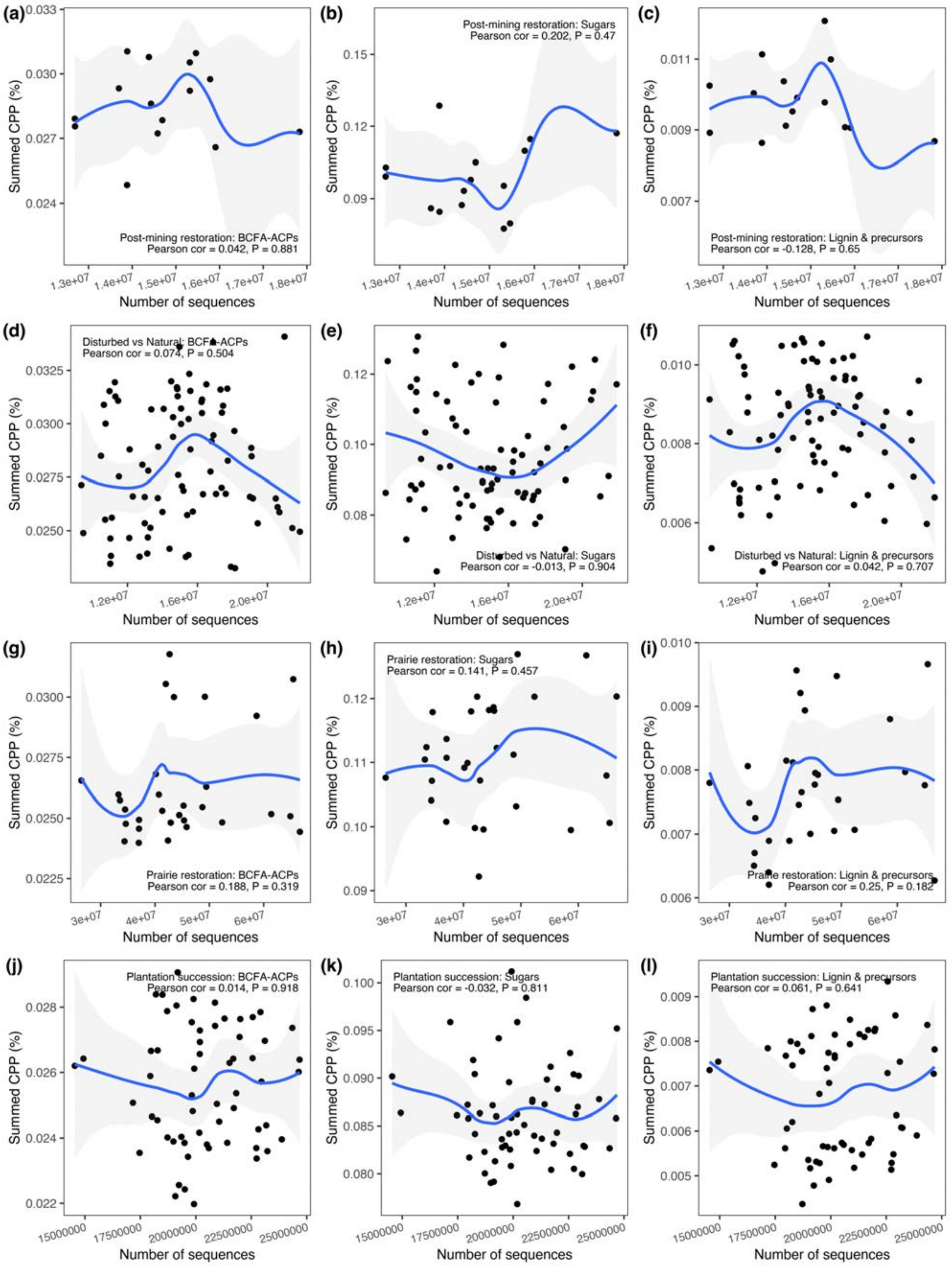


**Fig. S49.** Visualization and check for correlations between CPP trend groups (n = 36 BCFA-ACPs, n = 5 sugars, n = 3 lignin & precursors) and numbers of sequences in samples. Datasets are: post-mining restoration (a-c), disturbed vs. natural soils (d-f), prairie restoration (g-i), plantation succession (j-l), vegetation succession (m-o), T2D-CHN (p-r), and T2D-SWE (s-u). Data refer to numbers of cleaned sequences including host removal for T2D gut metagenomes. Samples are described in Extended Data Table 1.

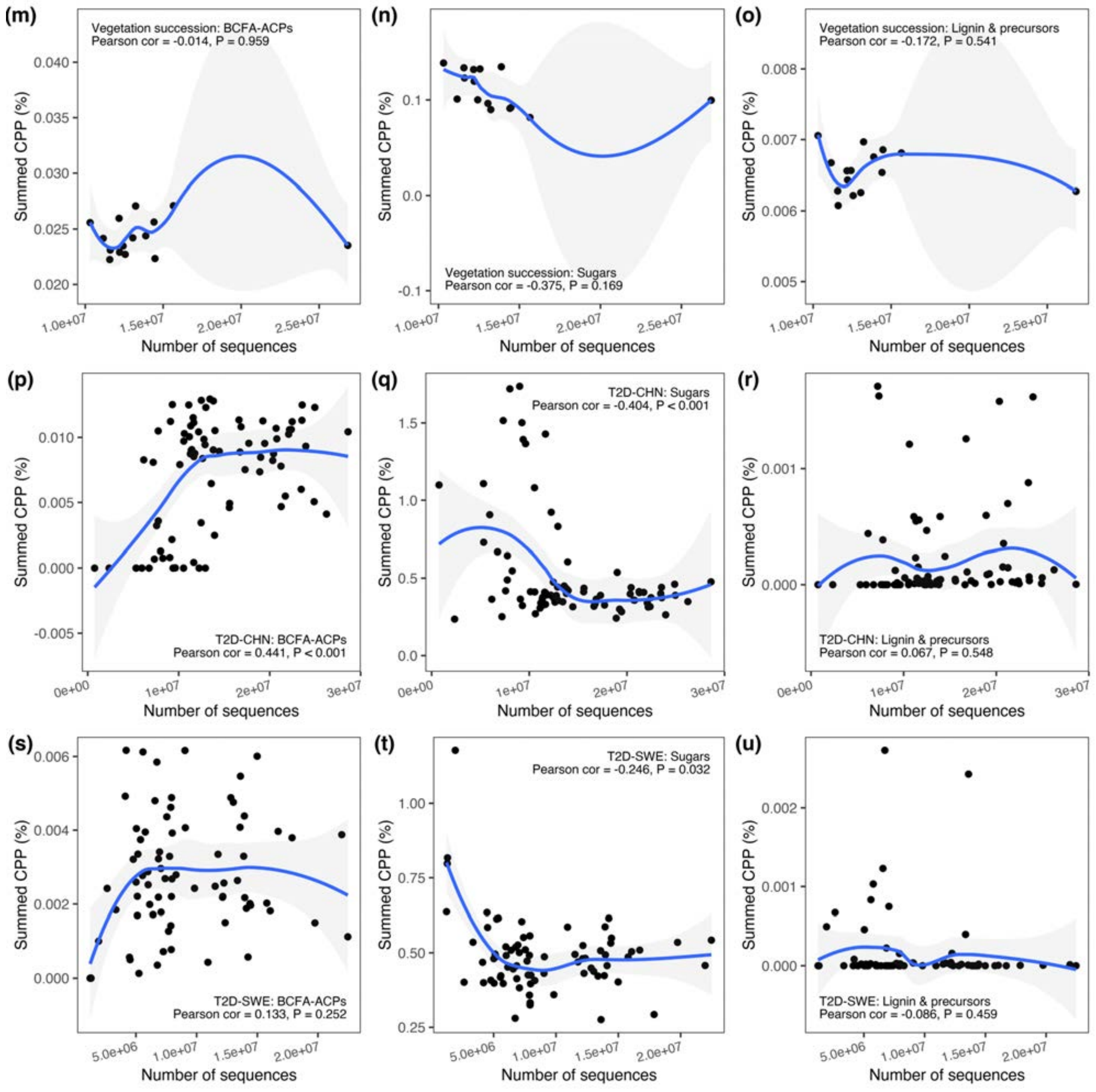


**Fig. S49. (continued)**

Rarefied to minimum library size

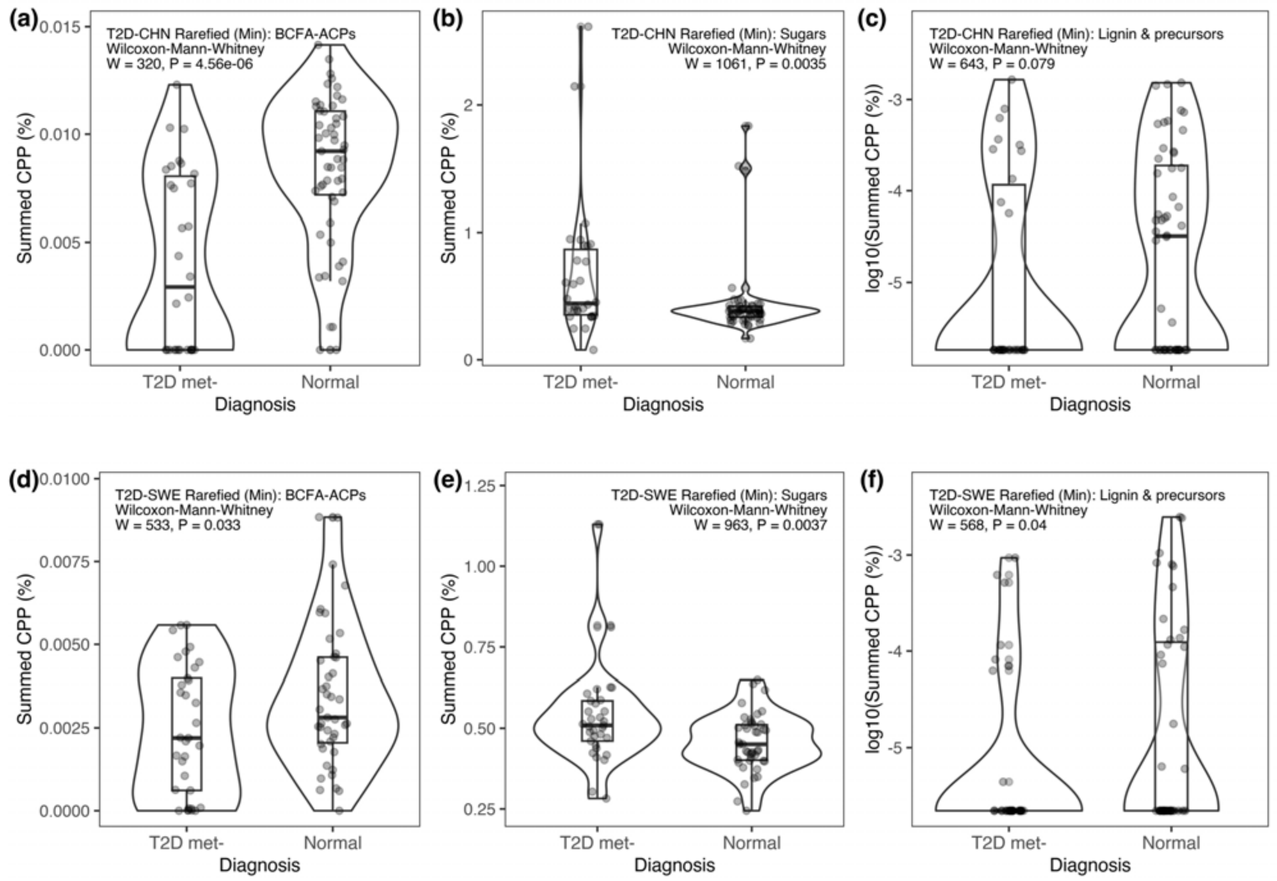


**Fig. S50.** Visualization and difference testing (Wilcoxon) between untreated T2D and normal healthy subjects for the CPP trend groups (n = 36 BCFA-ACPs, n = 5 sugars, n = 3 lignin & precursors) in samples that are rarefied to even sequencing depth at the **minimum library size** for respective datasets: (a-c) Chinese cohort (T2D-CHN; 758,416 sequences in each sample, total n = 82, n T2D = 30, n Normal = 52); and (d-f) Swedish cohort (T2D-SWE; 1,223,102 sequences in each sample, total n = 76, n T2D = 33, n Normal = 43).

Rarefied even sequences at 5th percentile

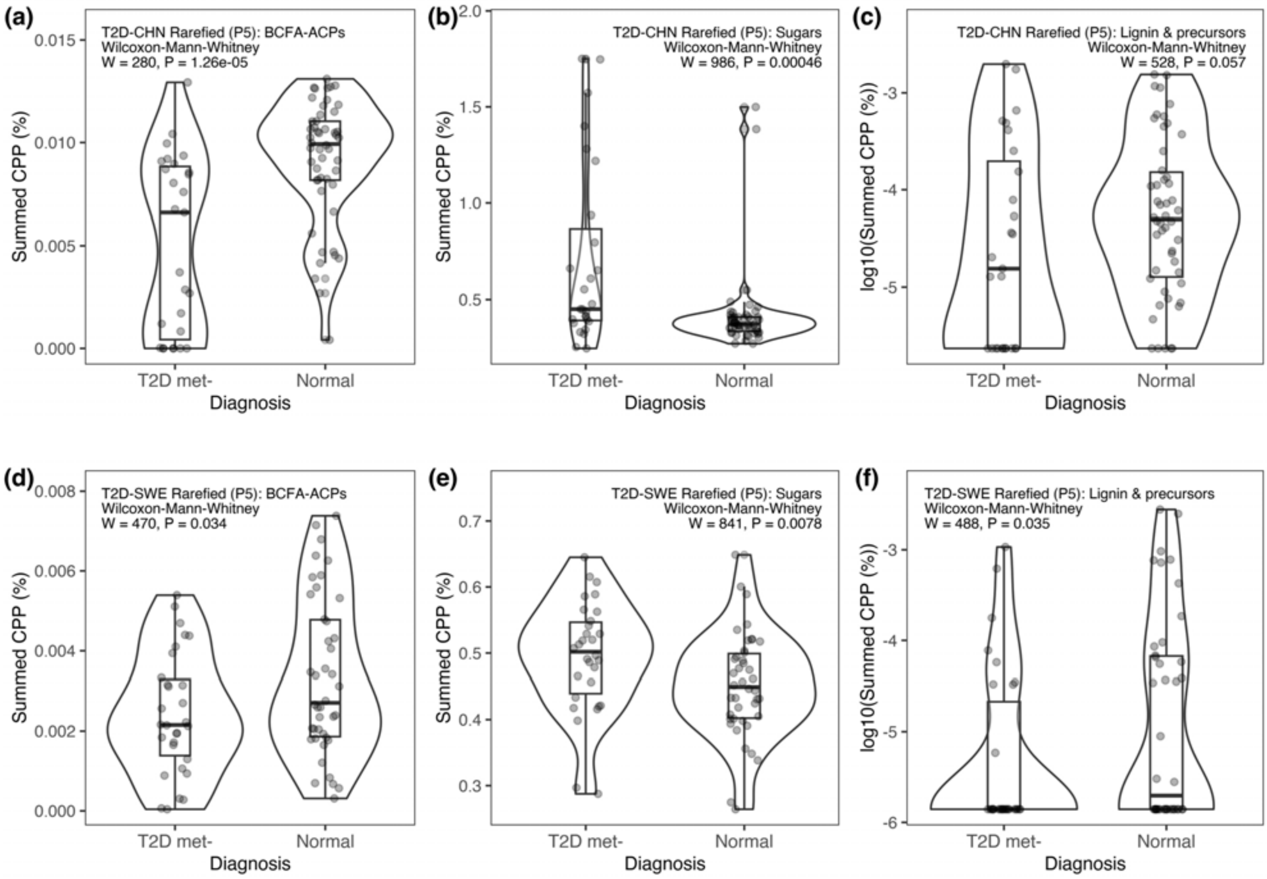


**Fig. S51.** Visualization and difference testing (Wilcoxon) between untreated T2D and normal healthy subjects for the CPP trend groups (n = 36 BCFA-ACPs, n = 5 sugars, n = 3 lignin & precursors) in samples that are rarefied to even sequencing depth at the **5th percentile of sequence counts** for respective datasets: (a-c) Chinese cohort (T2D-CHN; 5,962,972 sequences in each sample, total n = 77, n T2D = 27, n Normal = 50); and (d-f) Swedish cohort (T2D-SWE; 2,454,297 sequences in each sample, total n = 72, n T2D = 30, n Normal = 42).

Rarefied even sequences at 10th percentile

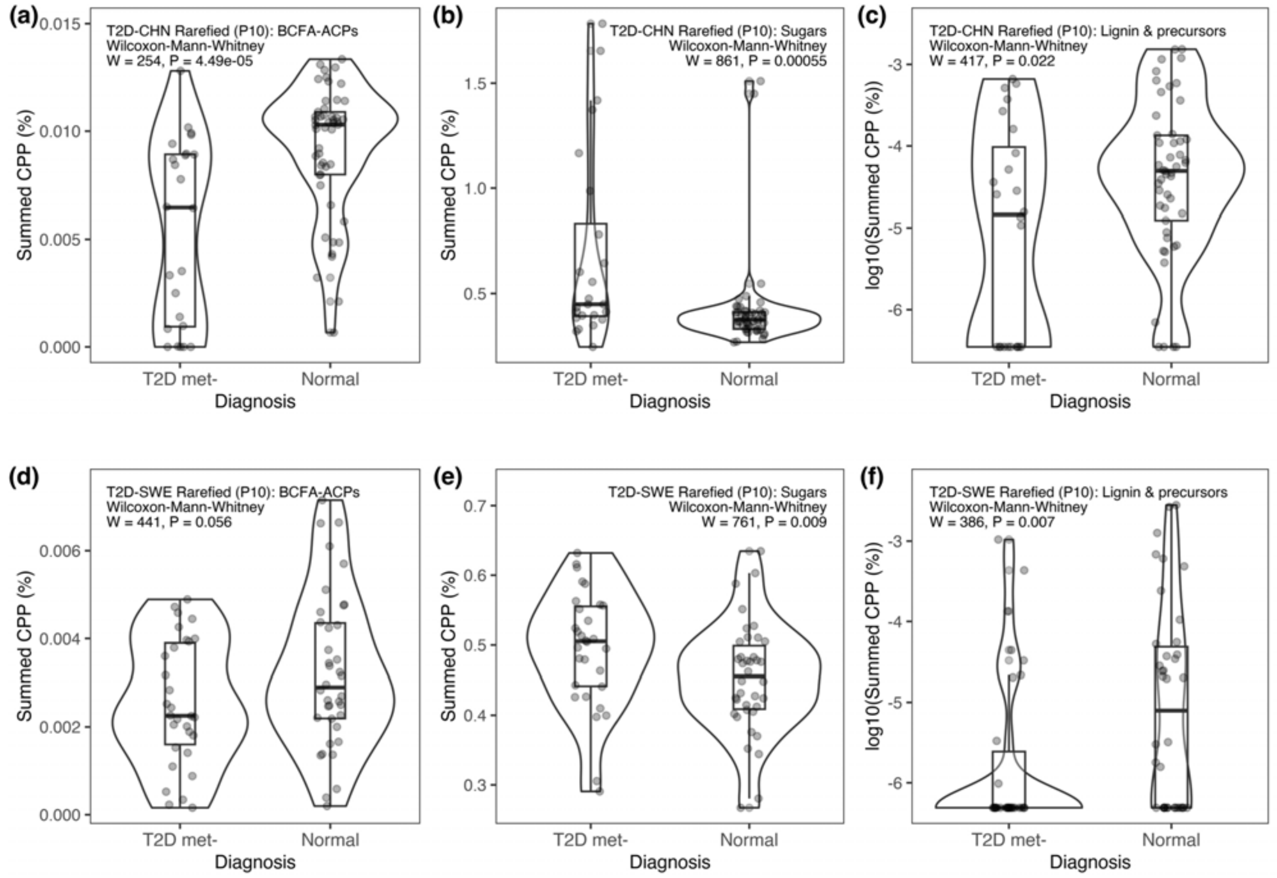


**Fig. S52.** Visualization and difference testing (Wilcoxon) between untreated T2D and normal healthy subjects for the CPP trend groups (n = 36 BCFA-ACPs, n = 5 sugars, n = 3 lignin & precursors) in samples that are rarefied to even sequencing depth at the **10th percentile of sequence counts** for respective datasets: (a-c) Chinese cohort (T2D-CHN; 7,356,437 sequences in each sample, total n = 73, n T2D = 24 , n Normal = 49); and (d-f) Swedish cohort (T2D-SWE; 4,330,969 sequences in each sample, total n = 68, nT2D = 30, n Normal = 38).

Rarefied even sequences at 15$^{th}$ percentile

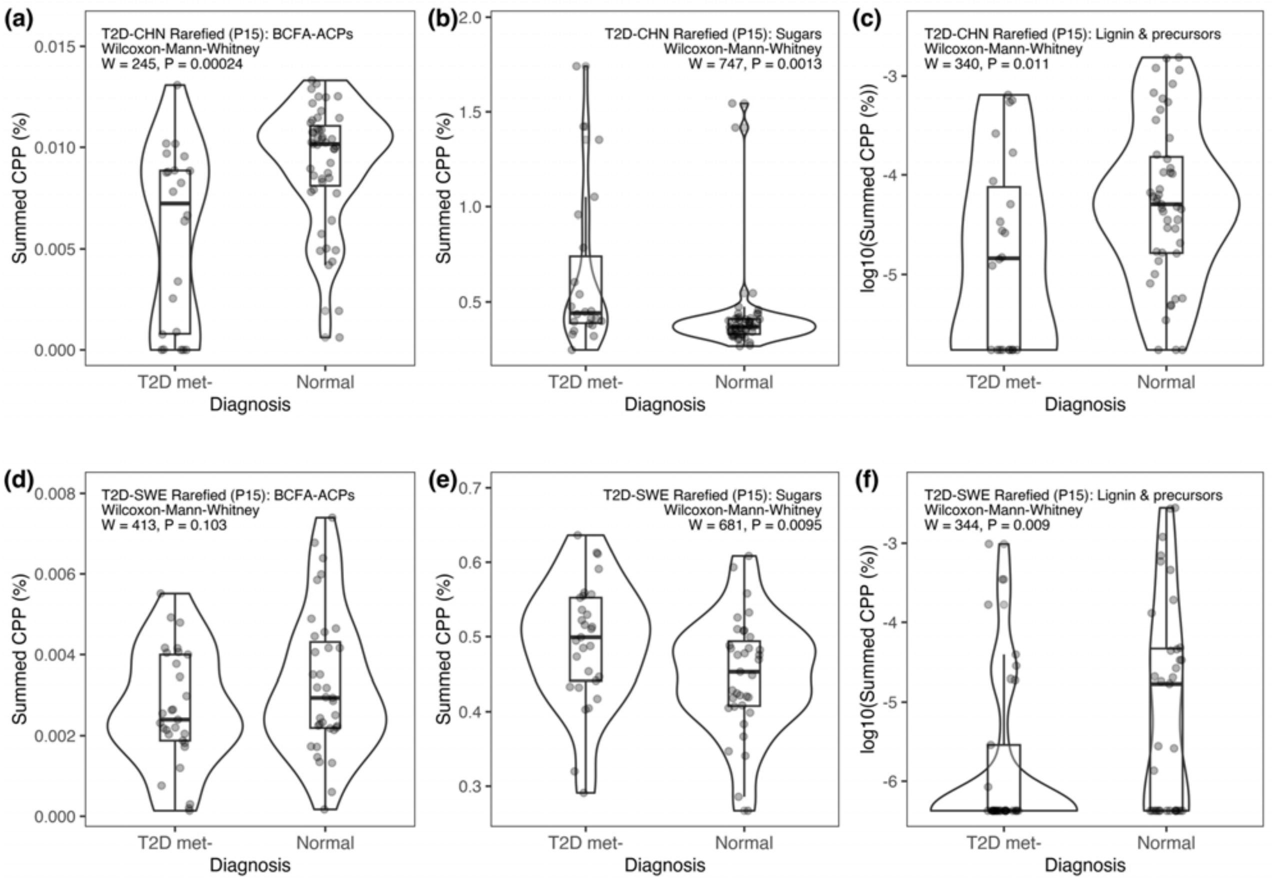


**Fig. S53.** Visualization and difference testing (Wilcoxon) between untreated T2D and normal healthy subjects for the CPP trend groups (n = 36 BCFA-ACPs, n = 5 sugars, n = 3 lignin & precursors) in samples that are rarefied to even sequencing depth at the **15$^{th}$ percentile of sequence counts** for respective datasets: (a-c) Chinese cohort (T2D-CHN; 8,050,681 sequences in each sample, total n = 69, n T2D = 22, n Normal = 47); and (d-f) Swedish cohort (T2D-SWE; 5,018,997 sequences in each sample, total n = 64, n T2D = 29, n Normal = 35).

Rarefied even sequences at 20th percentile

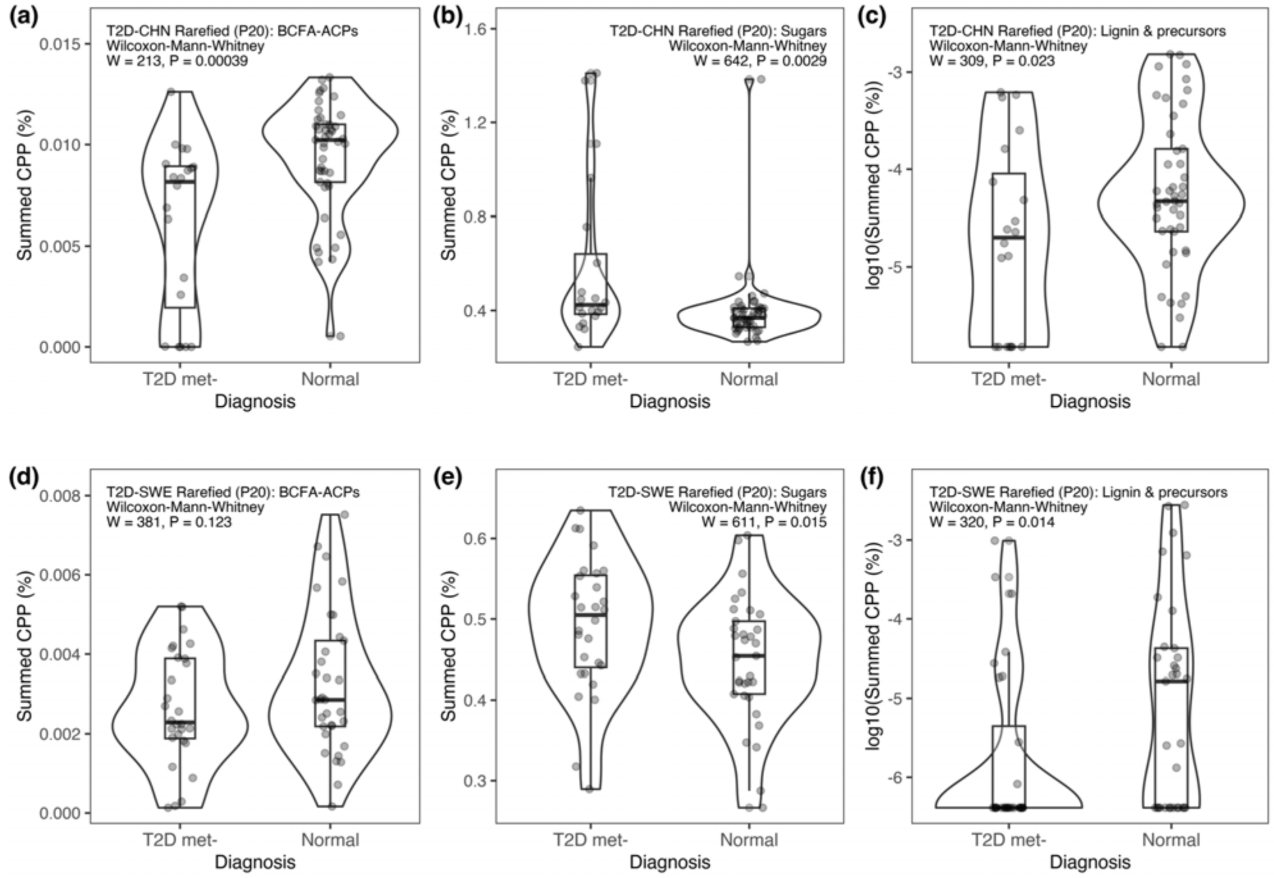


**Fig. S54.** Visualization and difference testing (Wilcoxon) between untreated T2D and normal healthy subjects for the CPP trend groups (n = 36 BCFA-ACPs, n = 5 sugars, n = 3 lignin & precursors) in samples that are rarefied to even sequencing depth at the **20th percentile of sequence counts** for respective datasets: (a-c) Chinese cohort (T2D-CHN; 9,261,724 sequences in each sample, total n = 65, n T2D = 20, n Normal = 45); and (d-f) Swedish cohort (T2D-SWE; 5,168,425 sequences in each sample, total n = 61, n T2D = 28, n Normal = 33).

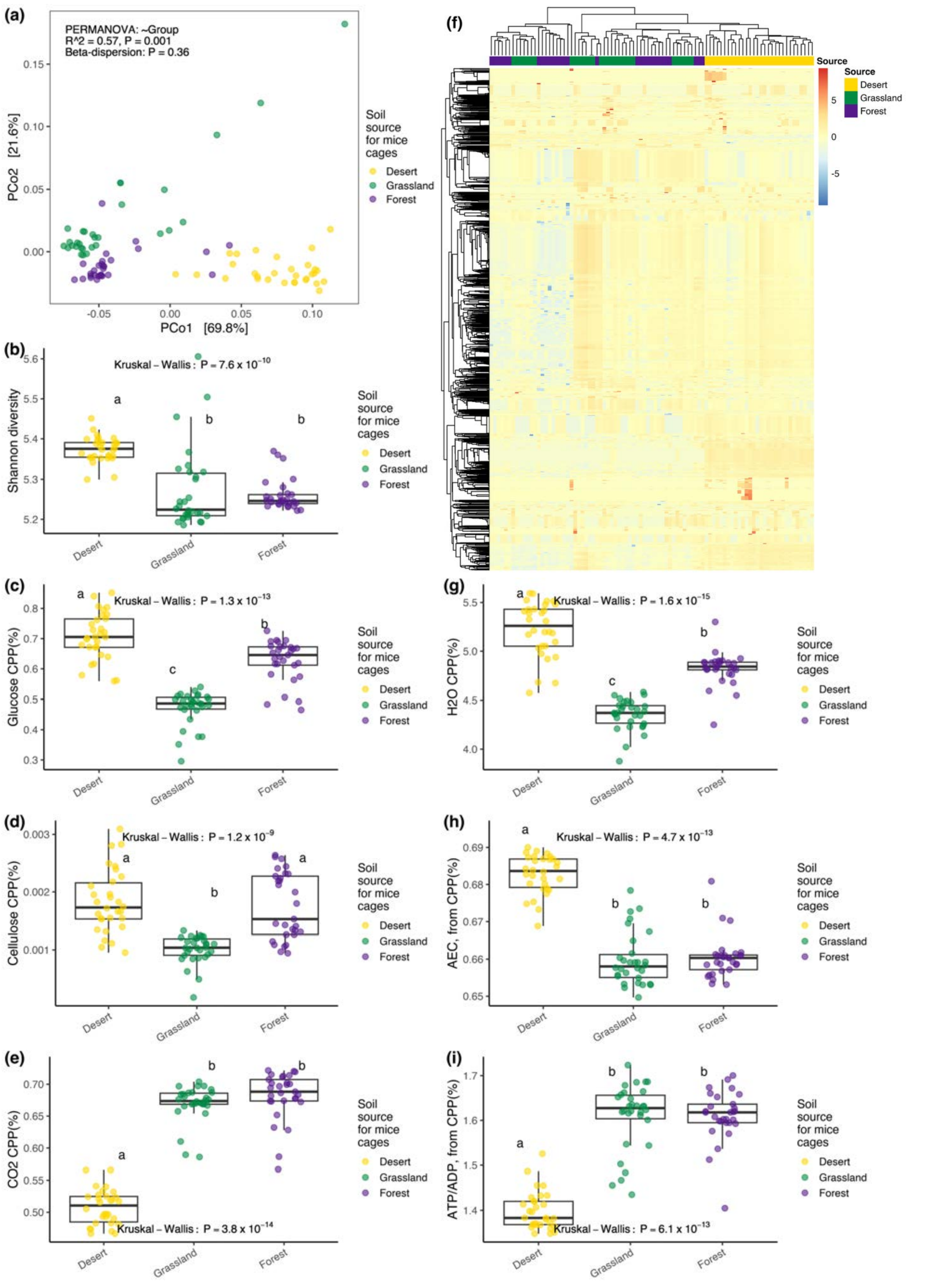


**Fig. S55.** CPP data from mice gut metagenomes with early life exposure to desert (n = 30), grassland (n = 30) and forest (n = 29) soils (Liu et al, 2021; SRA accession PRJNA542998). (a) Beta diversity, (b) alpha diversity, (c) glucose CPP, (d) cellulose CPP, (e) carbon dioxide CPP, (f) heat map for scaled log10(CPP) showing only >50$^{th}$ percentile variation (n = 3982 compounds), (g) water CPP, (h) adenylate energy charge (AEC = (ATP + 0.5xADP)/(ATP + ADP + AMP) based on CPP values, (i) ratio of adenosine triphosphate (ATP) to adenosine diphosphate (ADP) based on CPP values.

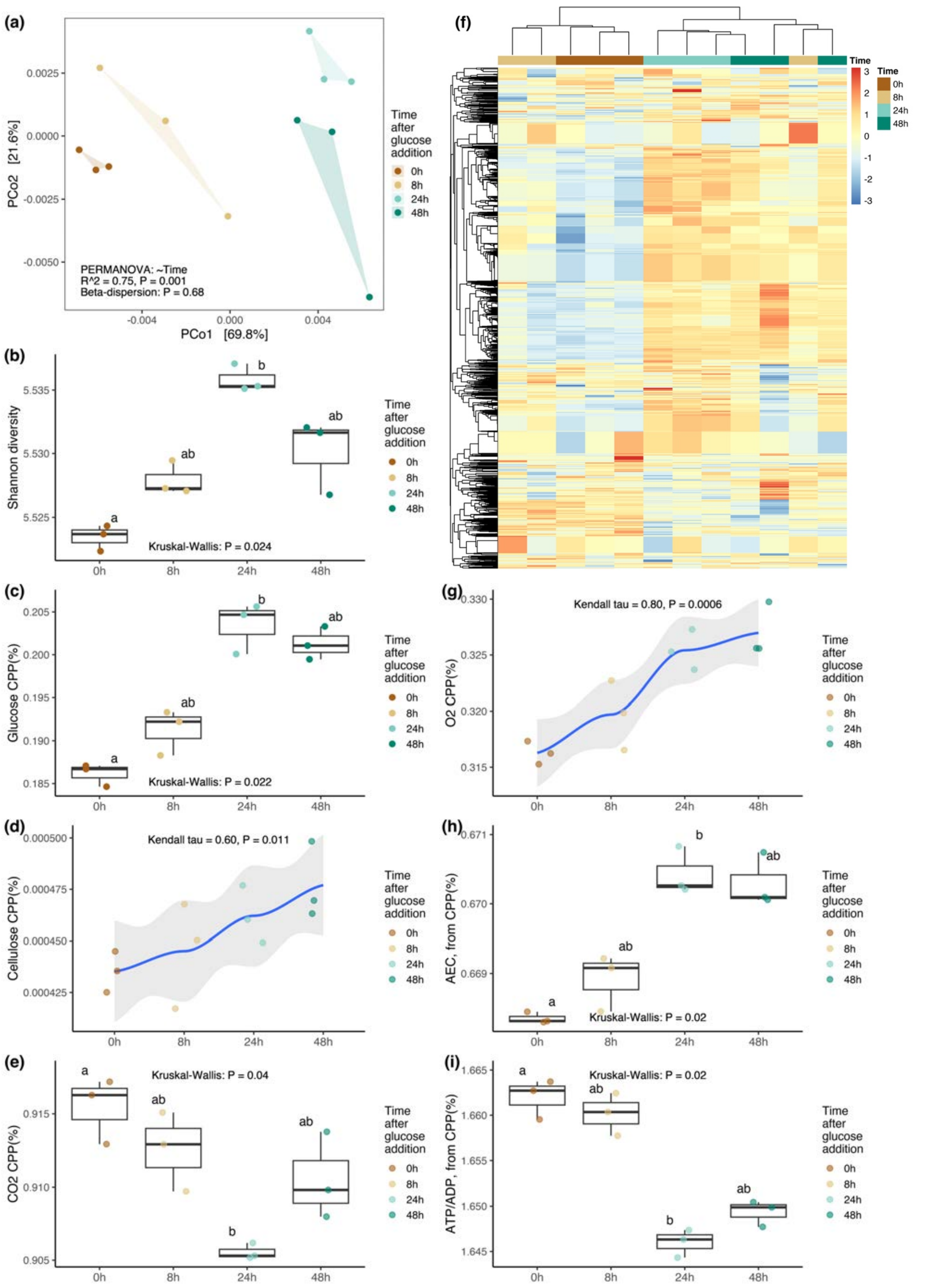


**Fig. S56.** CPP data for soil with glucose addition at time steps of 0, 8, 24 and 48h (n = groups of 3; Chuckran et al 2020). (a) Beta diversity, (b) alpha diversity, (c) glucose CPP, (d) cellulose CPP, (e) carbon dioxide CPP, (f) heat map for scaled log10(CPP) showing only >50$^{th}$ percentile variation (n = 4232 compounds), (g) oxygen CPP, (h) adenylate energy charge (AEC = (ATP + 0.5xADP)/(ATP + ADP + AMP) based on CPP values, (i) ratio of adenosine triphosphate (ATP) to adenosine diphosphate (ADP) based on CPP values.

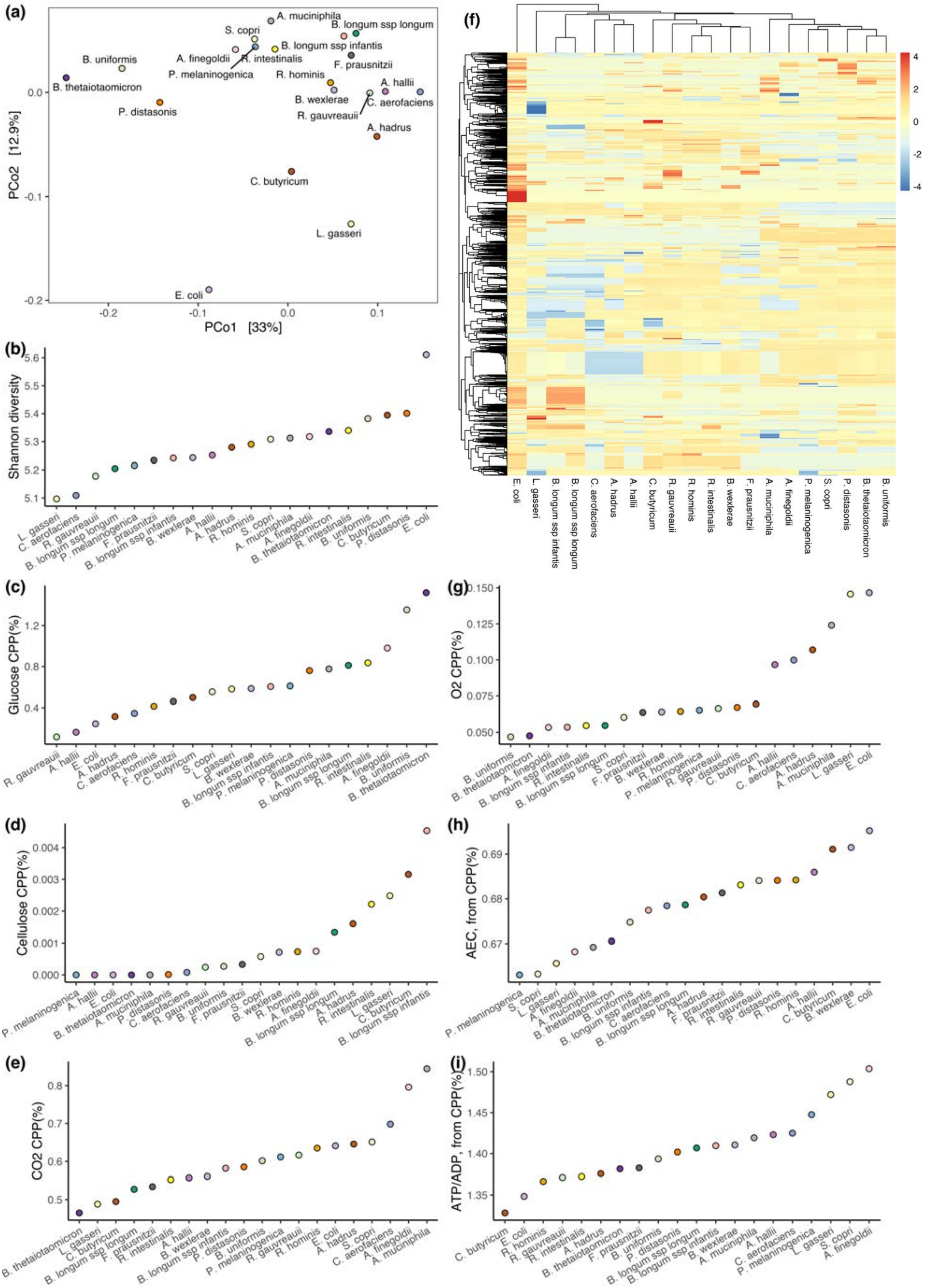


**Fig. S57.** CPP data for National Institute for Biological Standards and Control (NIBSC) bacterial cultures (Amos et al, 2020; SRA accession PRJNA622674). (a) Beta diversity, (b) alpha diversity, (c) glucose CPP, (d) cellulose CPP, (e) carbon dioxide CPP, (f) heat map for scaled log10(CPP) showing only >50$^{th}$ percentile variation (n = 3095 compounds), (g) oxygen CPP, (h) adenylate energy charge (AEC = (ATP + 0.5xADP)/(ATP + ADP + AMP) based on CPP values, (i) ratio of adenosine triphosphate (ATP) to adenosine diphosphate (ADP) based on CPP values.

**Table S1.** Results from rarefying analyses on T2D-CHN and T2D-SWE datasets. Key signals generally persisted when datasets were reanalysed with rarefaction to even read depths. Reanalysis following rarefaction of cleaned non-host sequences was performed at the minimum library size and using data subsets that were rarefied at the 5th, 10th, 15th, and 20th sequence count percentiles (samples with sequence counts below thresholds were excluded), yielding 25 out of 30 results (83%) with P value ≤0.05.

| Dataset | Rarefying threshold | No. of sequences used in rarefying reanalysis | Sample size | | | P-value of Wilcoxon-Mann-Whitney test T2D Met- vs Normal | | |
|---|---|---|---|---|---|---|---|---|
| | | | T2D Met- | Normal | Total | BCFA-ACPs | Sugars | Lignin and precursors |
| T2D-CHN | Minimum library | 758,416 | 30 | 52 | 82 | 4.56E-06 | 0.003 | 0.079 |
| T2D-SWE | Minimum library | 1,223,102 | 33 | 43 | 76 | 0.033 | 0.004 | 0.040 |
| T2D-CHN | 5th percentile | 5,962,972 | 27 | 50 | 77 | 1.26E-05 | 4.59E-04 | 0.057 |
| T2D-SWE | 5th percentile | 2,454,297 | 30 | 42 | 72 | 0.034 | 0.008 | 0.035 |
| T2D-CHN | 10th percentile | 7,356,437 | 24 | 49 | 73 | 4.49E-05 | 0.001 | 0.022 |
| T2D-SWE | 10th percentile | 4,330,969 | 30 | 38 | 68 | 0.056 | 0.009 | 0.007 |
| T2D-CHN | 15th percentile | 8,050,681 | 22 | 47 | 69 | 2.36E-04 | 0.001 | 0.011 |
| T2D-SWE | 15th percentile | 5,018,997 | 29 | 35 | 64 | 0.103 | 0.009 | 0.009 |
| T2D-CHN | 20th percentile | 9,261,724 | 20 | 45 | 65 | 3.87E-04 | 0.003 | 0.023 |
| T2D-SWE | 20th percentile | 5,168,425 | 28 | 33 | 61 | 0.123 | 0.015 | 0.014 |
